# Autonomous generalizability auditing of scientific foundation models

**DOI:** 10.64898/2026.11.03.763979

**Authors:** Yasha Ektefaie, Shrey Jain, Pardis Sabeti

## Abstract

Determining where scientific foundation models generalize beyond their development data is essential for reliable use, but aggregate benchmarks obscure performance limits and rigorous audits are difficult to conduct routinely. Here, we introduce AutoSPECTRA, a coding-agent protocol for systematic generalizability auditing. To inform its design, we reviewed 38 studies across five biological domains and evaluated coding agents, which generated hypotheses and empirical analyses but applied inconsistent standards for judging evidence. We then structured AutoSPECTRA around measurable generalization axes, prespecified analyses, controlled comparisons and independent confirmation. Across four biological foundation models, audits linked performance to protein sequence composition and relatedness to pretraining data (ESMFold2), representation of perturbations in training (STATE), effective spatial resolution (CONCH), and residue-level genomic, evolutionary and structural context (AlphaMissense). Agents also evaluated alternative explanations and rejected unsupported hypotheses. AutoSPECTRA provides a systematic approach to discovering and testing the limits of scientific foundation models.

## Main

As foundation models are increasingly released for broad use in scientific research [1–4], scientists need reliable ways to determine where these models perform well and where they fail. In biology, researchers commonly evaluate these models using aggregate benchmark scores to guide their adoption, but these scores can conceal limitations that matter for a particular application. Such limitations often become apparent only after release, through individual user reports, scattered negative results or informal observations. Although such evidence can reveal important limitations, it is difficult to reproduce and rarely provides a reliable basis for deciding whether a model is appropriate for a new dataset. Systematic evaluation of these limits would help scientists make informed decisions before relying on a model’s predictions.

Researchers use generalizability audits to examine model performance on new data, but lack a standard approach for deciding which forms of biological variation to test, how to measure them or what evidence is needed to support their conclusions. These choices can change the conclusion: a cellular model, for example, might accurately predict responses to familiar perturbations in new cell types but struggle to predict responses to perturbations absent from its training data. Prior studies of model generalizability have investigated how performance changes across different forms of biological variation [5–10], but have used disparate comparisons [11–16] and standards of validation [17–21]. Consequently, studies can reach different conclusions about similar performance patterns, depending on whether their analyses distinguish model-specific weaknesses from greater task difficulty or other variables that differ between the groups being compared.

Our earlier SPECTRA framework established a systematic approach to measuring generalizability across a spectrum of relatedness between training and test data [22]. Given a model, a dataset and a researcher-defined property of the inputs, SPECTRA constructs train–test splits with progressively less overlap in that property and measures performance across them. The resulting performance curve reveals how performance changes across regimes that individual benchmark splits leave uncharacterized. Applying the framework nevertheless requires researchers to identify a relevant property, assemble suitable data and interpret the resulting curve. Making generalizability audits routine therefore requires a way to identify informative axes, assemble suitable data and determine what the resulting evidence supports.

General-purpose coding agents offer a potential means of conducting generalizability audits autonomously. These systems can translate natural-language requests into executable computational analyses by locating public resources, writing and running code and revising their approach as results accumulate [23–25]. A request to audit a model’s generalizability, however, leaves the relevant biological variation and the analysis needed to test it unspecified. Agents must formulate hypotheses, choose informative comparisons and decide whether the evidence warrants retaining, revising or rejecting an interpretation. It remains unclear whether independent agents approach these decisions consistently or what guidance they need to produce scientifically supported conclusions.

Here, we set out to establish a reproducible procedure for autonomous generalizability auditing. We first examined literature across five biological domains to identify the recurring analytical practices underlying informative generalizability audits. We then investigated how coding agents approached the task without a prescribed axis or analysis plan, examining both their empirical investigations and their criteria for accepting conclusions. Drawing on these analyses, we developed AutoSPECTRA, a protocol that guides agents to propose measurable generalization axes, define them independently of observed target-model performance and test candidate relationships through controlled comparisons and independent confirmation. The protocol allows agents to revise or reject hypotheses and to conclude an audit without identifying a supported axis.

We evaluate AutoSPECTRA across four biological domains to assess whether agents can identify scientifically supported limits to generalization and whether independent audits recover related axes. The applications span protein structure prediction, cellular perturbation modeling, computational pathology and variant-effect prediction. For each model, we combine a featured end-to-end audit with independent executions to examine how agents formulate hypotheses, test alternative explanations and determine which conclusions the evidence supports.

## Results

### Literature review and agent experiments inform the design of AUTOSPECTRA

To define the scientific requirements of an autonomous generalizability audit, we examined how prior studies identified and validated changes in model performance across biological variation. We conducted a structured review of 38 studies spanning regulatory genomics [26], variant-effect prediction [12], protein structure and function [27], single-cell transcriptomics [28] and computational pathology [29]. We identified 13 recurring methodological criteria, including testing on data not used for model training [9, 17–19], examining a specific difference between training and evaluation data [8] and comparing performance with simple or matched reference methods to distinguish model-specific weaknesses from difficulty shared across methods [6, 11, 12]. We formalized these criteria as a rubric for generalizability audits (Supplementary Table 2).

We built on our earlier SPECTRA framework to turn the literature-derived criteria into a computational audit [22]. SPECTRA measures how model performance changes as test data become less similar to training data. We used the literature-informed criteria to add audit requirements not specified by the original framework (Supplementary Note 1, Supplementary Table 4). Following the designs used in the reviewed studies, we also allowed evaluation groups, rather than SPECTRA’s original construction of a graph relating all pairs of examples. From literature, we added requirements for comparisons with reference methods designed to capture suspected shortcuts [11, 12] and for tests of proposed explanations for performance declines [21, 30]. We retained SPECTRA’s central approach of measuring performance across predefined levels of an input property, such as protein length. We call such a property a generalization axis, as it allows us to examine how model performance varies across different levels of that property.

We tested what guidance coding agents needed to move from proposing hypotheses about generalizability to testing them with data. For each instruction condition, we conducted 24 independent runs spanning three combinations of agent models and coding environments and four target models. When asked to identify potential failures using their own knowledge and the public web, every run proposed hypotheses grounded in the literature, but none tested a candidate axis empirically (Supplementary Tables 5–8). When explicitly asked to quantify a relationship and test whether it reproduced on independent data, all 24 runs autonomously acquired public resources and completed a quantitative evaluation, reproducing a reported effect in 22 runs and yielding two interpretable negative results (Supplementary Tables 9–12). These experiments demonstrated that explicit instructions could enable agents to carry out empirical audits of previously reported relationships.

We then asked whether coding agents could discover relationships not previously documented for the target model. Agents generated diverse hypotheses but applied inconsistent standards for judging novelty of their findings and the strength of supporting evidence (Supplementary Table 1 and Supplementary Tables 13–16). Agents examining the same model and candidate axis disagreed about whether prior work had already reported the relationship or an equivalent finding. Four such disagreements involved 11 of the 24 runs. In two cases involving four runs, agents reached opposing conclusions from similar initial associations after choosing different confirmation data, groupings of examples or controls. Three runs also proposed or revised an axis after inspecting target-model errors, highlighting the need to separate evidence used to generate a hypothesis from evidence used to confirm it. The experiments showed where agents needed clearer rules for assessing novelty, defining an analysis before testing it, confirming findings independently and deciding when the evidence justified a conclusion.

We translated lessons from the literature review and agent experiments into explicit decision rules for AutoSPECTRA. The literature-derived rubric specifies the evidence an audit must contain. The rules require agents to define each axis and its analysis plan before evaluating target-model performance, compare performance with fixed reference methods independent of the target model, and confirm findings on data not used to select the axis. They also specify how to assess whether a finding is new and when to conclude an audit (Supplementary Tables 2 and 4). AutoSPECTRA leaves agents free to choose which input properties to investigate, while setting common standards for testing whether those properties are associated with changes in model performance and deciding whether to retain, revise or reject the proposed relationship.

### AUTOSPECTRA guides the discovery and validation of generalization axes

AutoSPECTRA guides a coding agent through a generalizability audit using five named components: the Distiller, Investigator, Dataset Scout and Fetcher, and the Auditor. It is a protocol rather than a multi-agent architecture: the components represent stages of the audit, not separate agents (Figure 1). The stages organize how the agent defines candidate generalization axes, obtains suitable evaluation data, measures performance and tests whether findings withstand controlled comparisons and independent confirmation. The agent can revise or reject candidate axes as evidence accumulates and records the investigation in an audit card.

**Figure 1:**
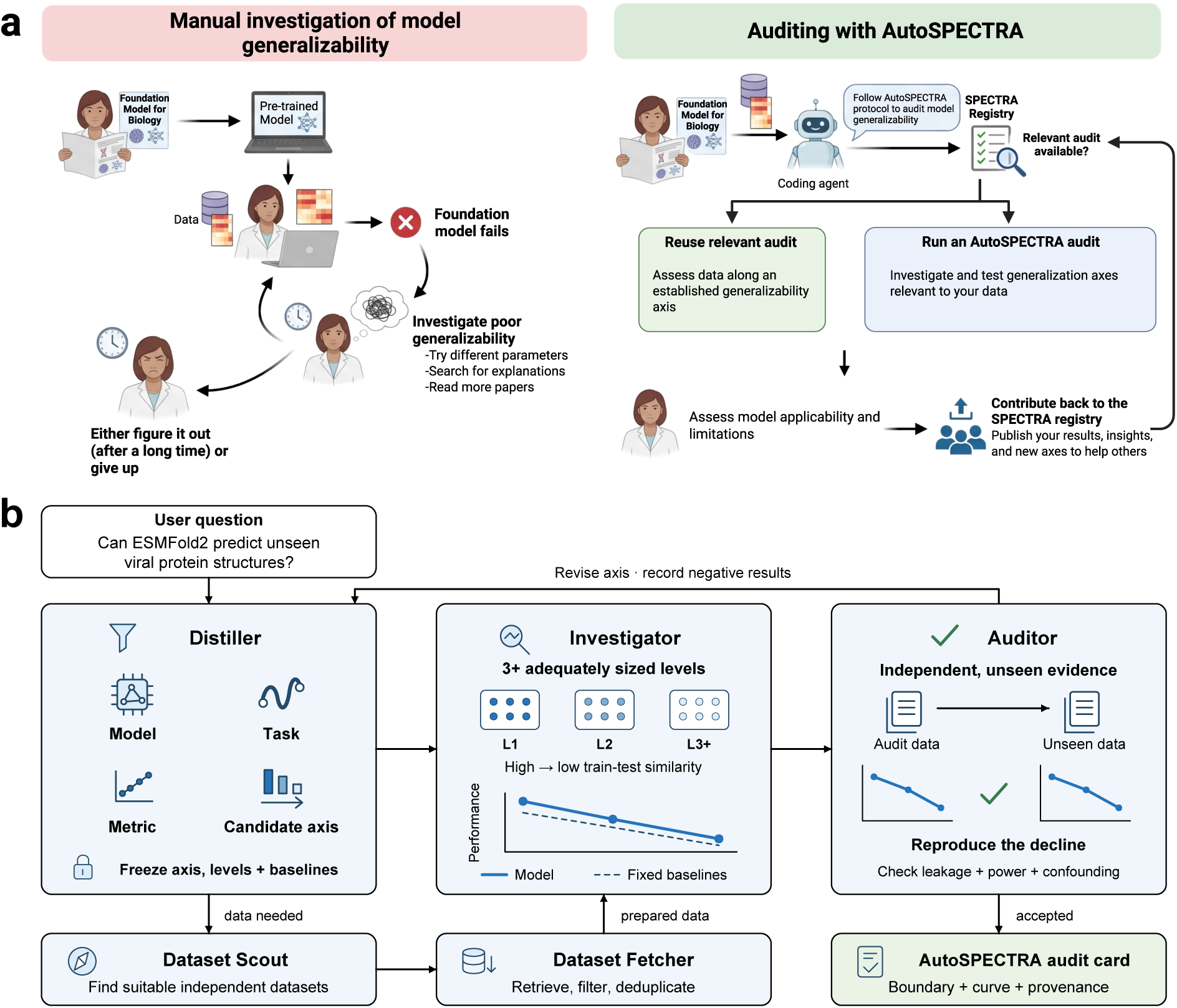
AutoSPECTRA structures an autonomous generalizability audit. (**a**) Manual investigation of unexpected model failures requires repeated, laborious analyses. With AutoSPECTRA, a coding agent retrieves relevant audits or conducts a new investigation to assess model generalizability. (**b**) AutoSPECTRA guides a coding agent through five stages. The Distiller proposes candidate axes; the Dataset Scout and Fetcher obtain evaluation data; the Investigator measures performance across decreasing train–test or pretraining–test similarity; and the Auditor checks validity and confirmation.

The Distiller translates the user’s question into a plan for the audit. The plan specifies the target model, prediction task, scientific unit of analysis, performance measure and candidate generalization axes. Candidate axes are measurable properties of the inputs or evaluation conditions, such as protein length or image resolution, that might explain variation in model performance. The agent must be able to measure that property and group examples by it without using the target model’s predictions. When existing benchmarks do not provide suitable data, the Dataset Scout and Fetcher identify, retrieve or construct the data needed for evaluation.

The Investigator tests how model performance changes across predefined levels of each candidate axis. Before examining the target model’s predictions, it measures the input property, defines at least three ordered levels and fixes the analysis plan. Evaluating the model across those levels produces a spectral performance curve. The agent also evaluates fixed reference methods independent of the target model on the same examples (S3, S5 and S9, Supplementary Table 4). Comparing performance across methods helps distinguish weaknesses specific to the target model from declines shared across methods.

AutoSPECTRA uses explicit evidence requirements to guide decisions about retaining, revising or rejecting a candidate axis. The Auditor validates a finding only when the observed performance decline survives checks for alternative explanations and reproduces on data not used to select the axis (S7 and S8, Supplementary Table 4). The confirmation data must support the same conclusion about how performance changes along the axis, although absolute performance may differ between datasets. If the evidence does not support a candidate axis, the agent can revise the axis, dataset or evaluation setup and continue the audit, or conclude without a supported axis.

AutoSPECTRA audit cards record what was tested, what the analyses showed and how the agent reached its conclusion. Each card documents the model and task, axis definitions, evaluation data, performance results and controls, along with rejected hypotheses, changes in interpretation and known limitations. Users can submit cards to a shared registry, where agents and researchers can retrieve prior findings, extend audits to new datasets or models and use previous investigations to formulate new questions.

To evaluate AutoSPECTRA across biological prediction tasks, we applied the protocol to four foundation models: ESMFold2 for protein structure prediction, STATE for cellular perturbation modeling, CONCH for computational pathology and AlphaMissense for variant-effect prediction. For each model, we present a featured audit to illustrate how agents propose, test and refine candidate generalization axes. We then compare independent audits to assess whether they identify the same axes or related measures of biological variation.

### ESMFold2 accuracy varies with protein sequence features and similarity to pretraining data

We first applied AutoSPECTRA to ESMFold2 [1], a foundation model for protein structure prediction, to assess how the protocol could characterize the generalizability of a newly released model. We initiated the audit on the day of ESMFold2’s release, before external performance analyses were available. AutoSPECTRA initially identified multiple sequence alignment (MSA) depth as a generalization axis using 138 proteins from Continuous Automated Model Evaluation (CAMEO) benchmark [31]. We then instructed the agent to investigate additional axes beyond evolutionary support. It explored three more axes, flexible or non-helical sequence composition, high sequence charge, and acidic windows, but rejected each after evaluating additional protein structures from Protein Data Bank (PDB) [32]. AutoSPECTRA subsequently identified high cysteine fraction as a candidate generalization axis and tested whether ESMFold2 accuracy varied across increasing levels of cysteine content. In an independent evaluation panel of 95 PDB sequences, mean C*α* local distance difference (C*α*-lDDT) declined from 0.901 in the lowest cysteine group (0.11%) to 0.787 in the highest (4.03%) (Figure 2a).

**Figure 2:**
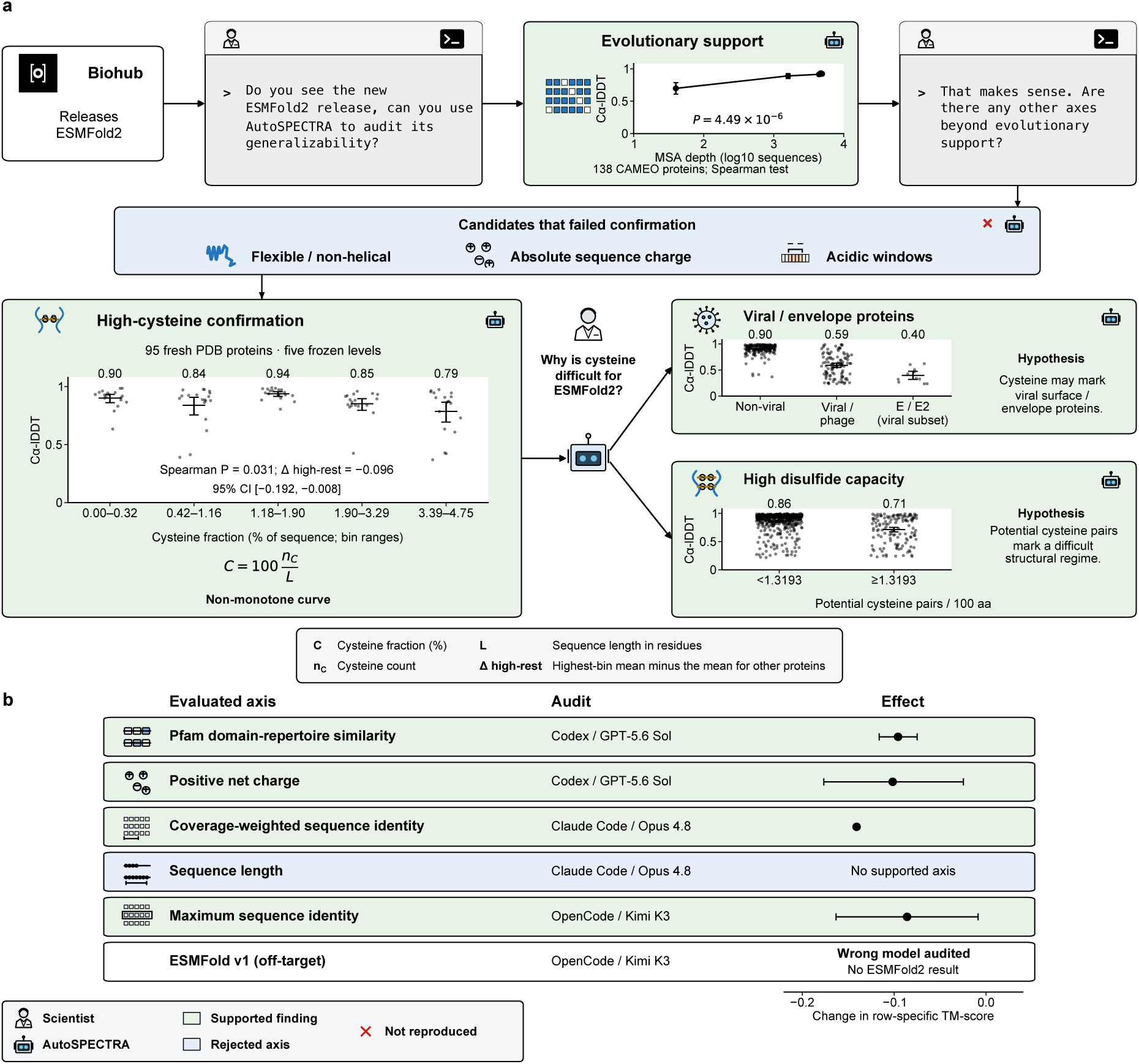
ESMFold2 accuracy varies with cysteine content and sequence relatedness to pretraining data. **a.** After initially identifying evolutionary support, AutoSPECTRA scanned three candidate axes which failed confirmation (Supplementary Figure 1a). AutoSPECTRA then identified high-cysteine composition as an axis, which it found to be attributable to viral envelope proteins and proteins with high estimated disulfide capacity. **b.** Three of six further audits linked accuracy to sequence or domain relatedness and one to positive net charge in isolated cryo-EM chains. One found no consistent length effect and one evaluated ESMFold1.

To investigate why proteins with higher cysteine fractions were predicted less accurately, we asked AutoSPECTRA to examine potential biological explanations for the association. The agent proposed two hypotheses. First, it examined whether viral proteins, particularly envelope proteins, contributed disproportionately to the lower accuracy. Of the 652 proteins it retrieved during its investigation, 289 had sufficient metadata for this analysis. Among these, the 105 virus- or phage-annotated proteins had a mean C*α*-lDDT of 0.591, compared with 0.899 for the other 184 proteins. The association was particularly pronounced among 11 proteins annotated as E/E2/envelope-E, which had a mean C*α*-lDDT of 0.396, compared with 0.803 for the other 278 metadata-eligible targets. AutoSPECTRA also hypothesized that disulfide-rich proteins might be harder to predict and investigated this using disulfide capacity as a sequence-derived proxy, calculated as half the cysteine count, rounded down and normalized per 100 amino acids. Among the 652 proteins, the 134 proteins at or above the 80th-percentile threshold had a mean C*α*-lDDT of 0.713 compared with 0.862 for the remaining 518 proteins.

We next examined whether independent AutoSPECTRA audits using different large language models (LLMs) and coding environments would identify similar or complementary generalization axes. We conducted six additional audits varying the underlying coding-agent harness and LLM (Figure 2b). One audit using Kimi K3 investigated ESMFold1 rather than ESMFold2 and did not yield a relevant result. Of the remaining five audits, three identified axes related to protein novelty, including Pfam domain-architecture novelty, coverage-weighted sequence identity and maximum sequence identity (Supplementary Table 19). Another audit identified an association with net positive charge among isolated chains studied by cryo-electron microscopy, while a sequence-length audit returned no supported axis after it found no consistent decline in ESMFold2 performance with increasing sequence length. Convergence among the repeat audits therefore concerned a broader family of relatedness measures, rather than independent reproduction of the featured disulfide-capacity finding.

The independent audits also demonstrated how AutoSPECTRA evaluates the evidence supporting a generalization axis, testing alternative explanations and comparing performance with reference methods. First, it tested whether other protein features could explain the performance differences observed across an axis. In the domain-novelty audit, it examined whether poorer performance on proteins with novel domain repertoires could instead reflect greater disorder or related sequence properties. The association persisted after adjusting for these and other protein properties (adjusted coefficient 0.053, P = 0.00044; Supplementary Table 19). Second, the audits compared performance with reference methods. A simple template-transfer baseline retrieved the most similar available protein sequence from structures deposited before the training cutoff and transferred its known structure to the target. Both domain novelty and net positive charge were associated with reduced accuracy in this baseline as well as ESMFold2, indicating difficulty shared with template-based prediction. One audit additionally retrieved AlphaFold2 predictions from the AlphaFold Database to investigate sensitivity to low sequence relatedness, recognizing that template transfer inherently struggles when related structures are absent. From highly related to orphan proteins, mean TM-score declined by 0.141 for ESMFold2 and 0.063 for AlphaFold2, supporting greater sensitivity in ESMFold2 on the same evaluation targets (Supplementary Table 19).

### STATE performance improves with training support for the queried perturbation

We applied AutoSPECTRA to STATE [4], a single-cell perturbation-effect prediction model, using precomputed outputs from Tahoe-100M [33], Replogle–Nadig [34, 35] and Parse-PBMC. AutoSPECTRA evaluated performance using overlap@N, the fraction of experimentally significant differentially expressed genes recovered among the top N predicted significant genes ranked by the magnitude of their predicted log-fold changes. It evaluated STATE in two configurations, using highly variable genes (HVG) expression or learned STATE embedding (SE) representations. AutoSPECTRA first tested chemical similarity to training drugs, measured by Tanimoto similarity between molecular fingerprints, but found no meaningful performance gradient. The agent next investigated whether functional similarity to genes represented in training could better explain variation in prediction accuracy. In Replogle, AutoSPECTRA measured each test gene’s maximum Jaccard similarity to training genes’ Reactome pathway annotations. This yielded two populated groups among annotated genes. In Jurkat and RPE1, mean overlap@N decreased from 0.272 to 0.176 for HVG and from 0.219 to 0.134 for SE between higher-similarity (>0.5) and lower-similarity (≤0.5) groups (Figure 3a).

**Figure 3:**
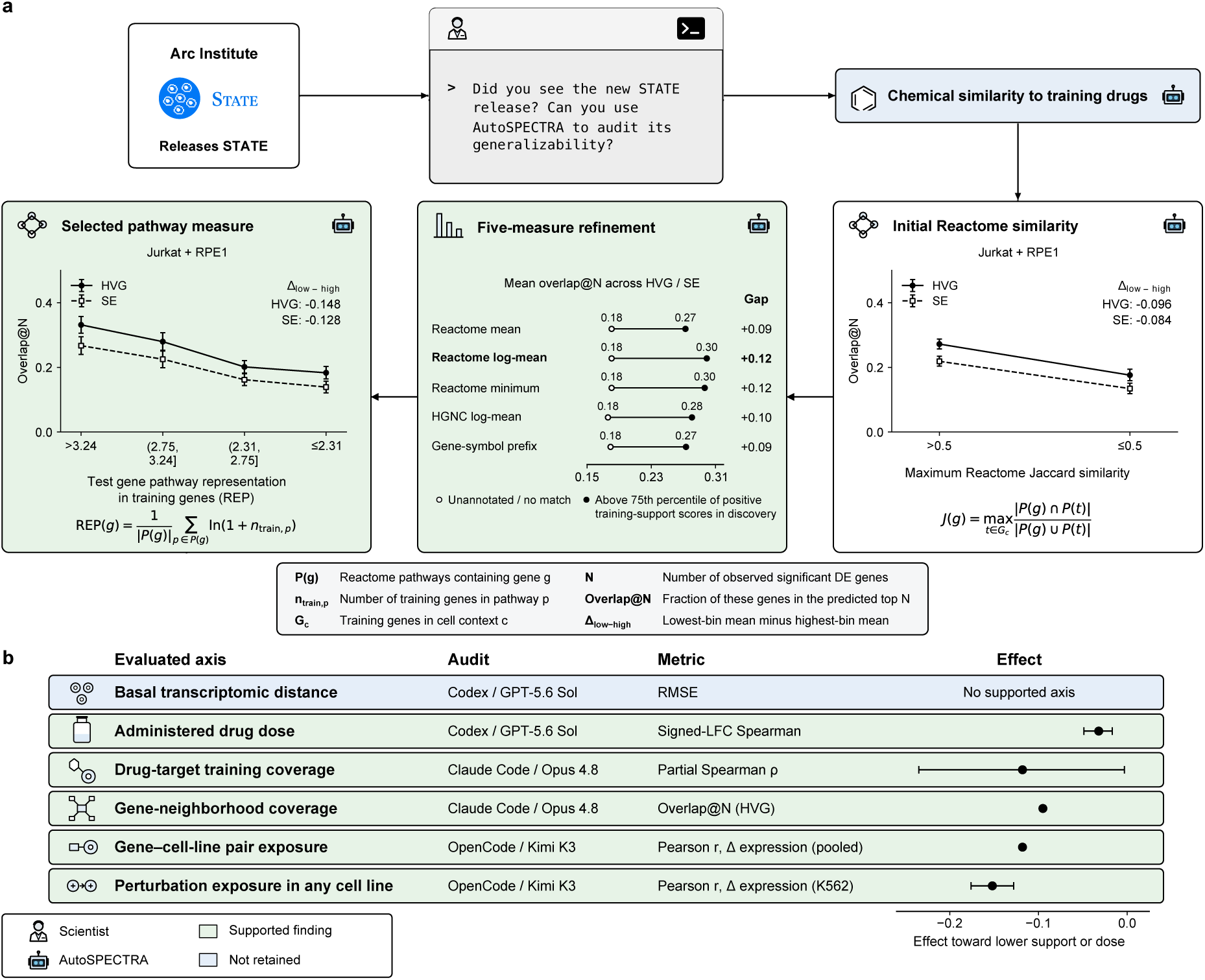
STATE performance improves when the queried perturbation is more strongly represented in the training data. **a.** After rejecting chemical similarity as an axis, AutoSPECTRA identified Reactome pathway support as a potential axis. After testing five different definitions of the axis, it found a decrease in performance as pathway support decreased which persisted in zero-shot evaluation and after controlling for confounders (Supplementary Figure 2). **b.** Four further audits linked accuracy to target or gene-neighborhood coverage or to prior exposure to the perturbation. A fifth found lower accuracy at low dose. Basal transcriptomic distance showed no supported decline.

To investigate whether other measures of functional representation in training could better characterize this relationship, AutoSPECTRA compared five measures of training support: the mean count, mean log(1 + count), and minimum count of training genes sharing each queried gene’s Reactome pathways; mean log(1 + count) across gene groups curated by the Human Genome Organisation’s Gene Nomenclature Committee (HGNC); and the count sharing its gene-symbol prefix. All five passed screening. Reactome log-mean support produced the largest average performance difference between the highest-support and unannotated groups across both representations in Jurkat and RPE1. It also resolved four populated levels among annotated genes, across which mean overlap@N increased progressively, from 0.183 to 0.331 for HVG and from 0.139 to 0.267 for SE, with positive associations across HepG2, K562, Jurkat and RPE1. AutoSPECTRA then tested whether the association could instead reflect differences in annotation availability, annotation count, or pathway catalog. The relationship persisted after excluding unannotated genes, matching annotation counts, and extending to zero-shot evaluation.

We examined whether independent AutoSPECTRA audits using different LLMs and coding environments would identify related generalization axes for STATE. We conducted six additional audits using GPT-5.6 Sol, Claude Opus 4.8 and Kimi K3 in their respective coding-agent harnesses (Supplementary Table 20). Five identified supported performance associations (Figure 3b). Two identified related forms of functional training support: drug responses were predicted more accurately when their target genes were represented in the available gene-knockdown panel, and gene perturbations were predicted more accurately when the perturbed gene had close functional counterparts elsewhere in the panel. Two other audits examined training exposure directly, finding better predictions when the exact gene–cell-line combination, or the gene perturbation in any cell line, had appeared in training. A fifth supported a low-dose performance decline, treating dose as an experimental condition rather than a measure of novelty relative to training.

One Kimi K3 audit further investigated why previously unseen perturbations were particularly difficult to predict. It initially tested whether accuracy improved progressively with the number of training examples. When that gradient failed, it reformulated the hypothesis around a sharper distinction: whether the perturbation had ever appeared in training. It confirmed this distinction in three additional cell lines; across all four lines, correlations between predicted and observed expression changes were 0.045–0.152 lower for unseen perturbations. Inspection of STATE’s learned perturbation representations then revealed nearly zero numerical vectors for unseen perturbations, consistent with the model receiving little information identifying which gene had been perturbed. The audit thus progressed from testing a generalization axis to identifying a model-level explanation for the observed performance difference.

### CONCH performance declines disproportionately under reduced spatial resolution

We applied AutoSPECTRA to CONCH [3], a pathology vision–language foundation model, to investigate how variation in tissue image characteristics affects prediction accuracy. AutoSPECTRA selected zero-shot classification of colorectal hematoxylin-and-eosin (H&E) tissue tiles, a task for which CONCH provided official prompts. It evaluated 7,180 publicly available tiles across nine tissue classes and used 1,800 additional class-balanced tiles to establish a reference appearance profile [36, 37]. AutoSPECTRA first investigated whether tiles that differed more from the reference images were harder to classify. Before evaluating CONCH, it defined axes based on color distributions, texture features capturing edges and grayscale variation, their combination, and the proportions of tissue, white, dark and highly saturated pixels. It initially prioritized combined color–texture similarity, averaging each test tile’s distance to its 20 nearest reference images. An initial association between lower image similarity and poorer classification disappeared after AutoSPECTRA accounted for tissue class: the least similar groups contained more readily classified fat tissue and blank background.

To determine whether image characteristics were associated with prediction accuracy independently of differences in tissue-class composition, AutoSPECTRA next examined the predefined axes within each tissue class. Among tumor tiles, recall, the fraction of tumor tiles correctly classified as tumor, declined from 0.779 to 0.114 across six groups with increasingly atypical tissue and stain appearance, measured by their proportions of tissue, white, dark and highly saturated pixels. Because this association emerged after examining the initial results, AutoSPECTRA tested it on 1,800 fresh tumor tiles, defining six groups using the same tissue and stain appearance measure before evaluating performance. Recall remained between 0.883 and 0.930 in the four highest-similarity groups but fell to 0.697–0.730 in the two lowest, supporting an association between atypical tissue and stain appearance and poorer tumor recognition (Figure 4a).

**Figure 4:**
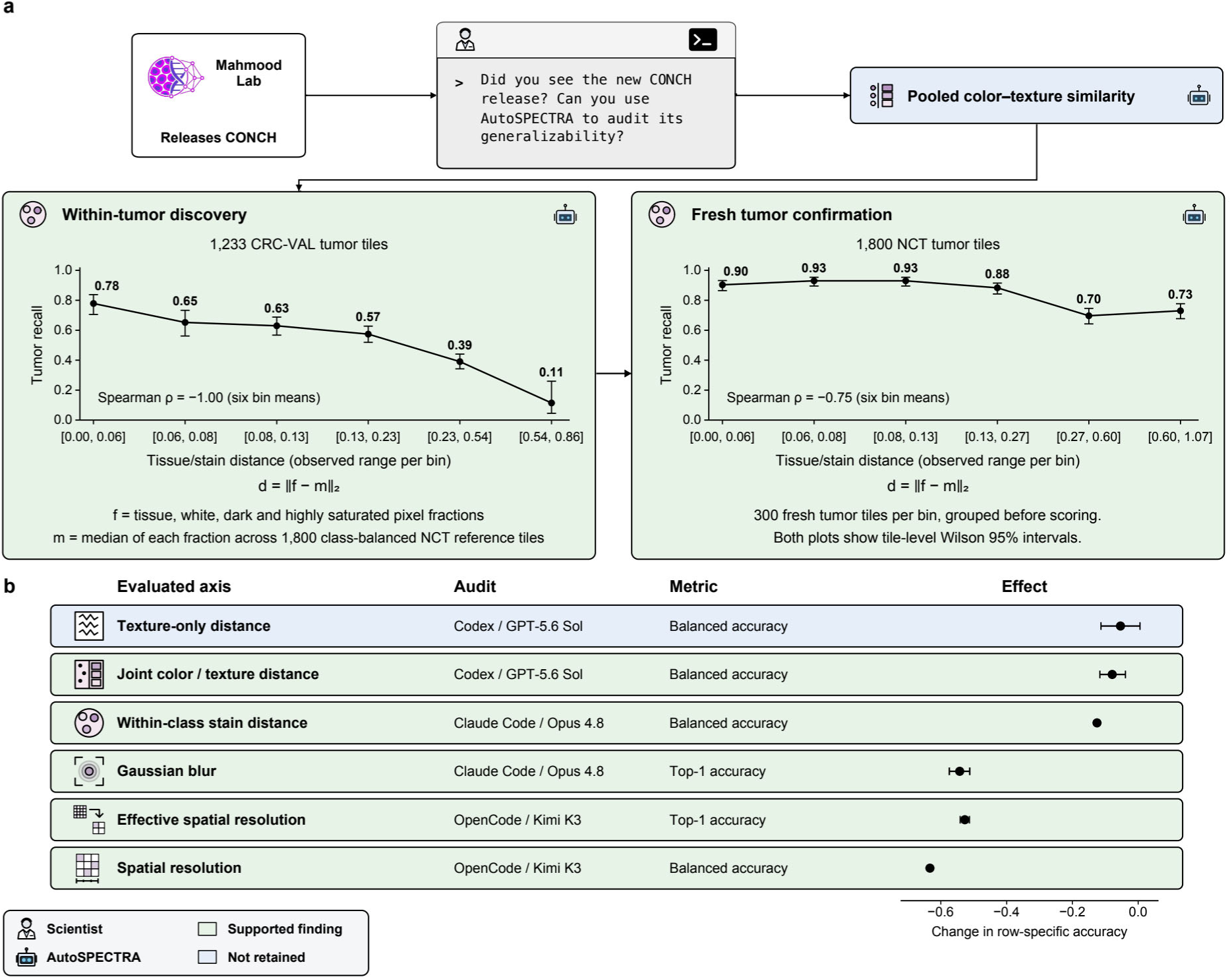
CONCH accuracy declines with atypical tumor appearance and loss of fine image detail. **a.** After finding that the pooled appearance axis depended on tissue-class composition, AutoSPECTRA examined the axis within each tissue class. It identified an association between atypical tissue and stain appearance and lower tumor recall, which persisted on fresh tumor tiles (Supplementary Figure 3). **b.** Five of six further audits linked performance to appearance, blur or spatial resolution. Blur and reduced spatial resolution produced larger performance declines in CONCH than in the tested reference models.

We examined whether independent AutoSPECTRA audits would identify similar or additional generalization axes for CONCH. We conducted six additional audits, five of which returned a supported axis (Figure 4b). Two identified associations with differences in color, texture or staining after accounting for tissue class, extending the initial appearance-based investigation. A third audit tested controlled Gaussian blur using a frozen analysis on 1,080 separate, class-balanced NCT tiles and found declining accuracy as blur increased. Two other audits independently identified effective spatial resolution as a generalization axis, examining how performance changed as they progressively removed fine image detail by downsampling each tile and resizing it back to the model’s input dimensions.

Comparisons with reference models helped distinguish performance declines specific to CONCH from those reflecting difficulty shared across models. Most natural appearance axes reflected shared image difficulty, with baseline models deteriorating by comparable or larger amounts than CONCH. Reduced effective resolution, in contrast, disproportionately impaired CONCH. In one audit, CONCH top-1 accuracy decreased from 0.803 at native resolution to 0.276 at 4.0 *μ*m pixel^−1^, whereas an ImageNet-pretrained ResNet-50 evaluated on the identical transformed images decreased from 0.795 to 0.564, corresponding to relative losses of 65.6% and 29.1%, respectively. A second audit confirmed this sensitivity on 3,600 additional held-out colorectal tissue tiles: CONCH balanced accuracy decreased from 0.771 to 0.138, compared with 0.969 to 0.667 for a DINOv2-based classifier. The same pattern appeared in the original 7,180-tile evaluation set. In the blur audit, CONCH accuracy fell by 0.543 compared with 0.323 for a fixed ResNet-50 baseline. Together, these comparisons identified loss of fine image detail, through reduced resolution or blur, as a condition under which CONCH deteriorated more than the tested reference classifiers.

### AlphaMissense performance varies with residue-level genomic, evolutionary and structural context

We finally applied AutoSPECTRA to AlphaMissense (AM), a proteome-wide missense pathogenicity predictor that integrates evolutionary sequence context and predicted structural information, to investigate how prediction accuracy varies with available biological information [2, 38]. Using released predictions matched to ClinVar classifications, AutoSPECTRA first tested whether accuracy declined for variants in proteins with fewer known relatives or genes with less observed population variation. It quantified evolutionary support by counting related sequences in the UniRef50 sequence database and population-variation support by calculating the fraction of possible missense variants observed for each gene in the Genome Aggregation Database (gnomAD). Neither axis showed a consistent performance decline across ten groups ordered by decreasing support.

These negative results led AutoSPECTRA to examine whether protein-level measures overlooked important variation between positions within the same protein. All variants in a protein received the same homolog count, although the number of sequences aligned at individual positions could differ substantially. AutoSPECTRA therefore shifted its investigation from protein-level properties to candidate generalization axes defined at individual residues. To investigate evolutionary support at the mutated position using experimental measurements, AutoSPECTRA analyzed 72 deep mutational scanning (DMS) assays spanning 63 human proteins in ProteinGym v1.3 [39]. It initially considered effective alignment depth, which downweights closely related sequences, but the proposed low-depth groups contained few proteins. It therefore prioritized alignment occupancy, the fraction of aligned sequences containing an amino acid rather than a gap at each position, and defined five groups from the occupancy distribution before evaluating performance.

Agreement between AlphaMissense predictions and experimentally measured variant effects, quantified by Spearman rank correlations, declined monotonically from 0.515 in the highest-occupancy quintile to 0.395 in the lowest (Figure 5a). AutoSPECTRA next tested whether this association persisted even at positions with substantial evolutionary sequence support. Among positions supported by at least 200 effective sequences, agreement was 0.475 at occupancy ≥90% and 0.305 at occupancy <70%. Comparisons with reference methods helped determine whether the performance decline was specific to AlphaMissense. Both a score based on position-specific amino-acid frequencies and BLOSUM62, which scores substitution types without considering their position, showed declining performance with lower occupancy [40]. AlphaMissense remained the most accurate of the tested predictors. The decline even in the position-independent BLOSUM62 baseline suggested that low occupancy marked difficulty shared across predictors, rather than an AlphaMissense-specific limitation.

**Figure 5:**
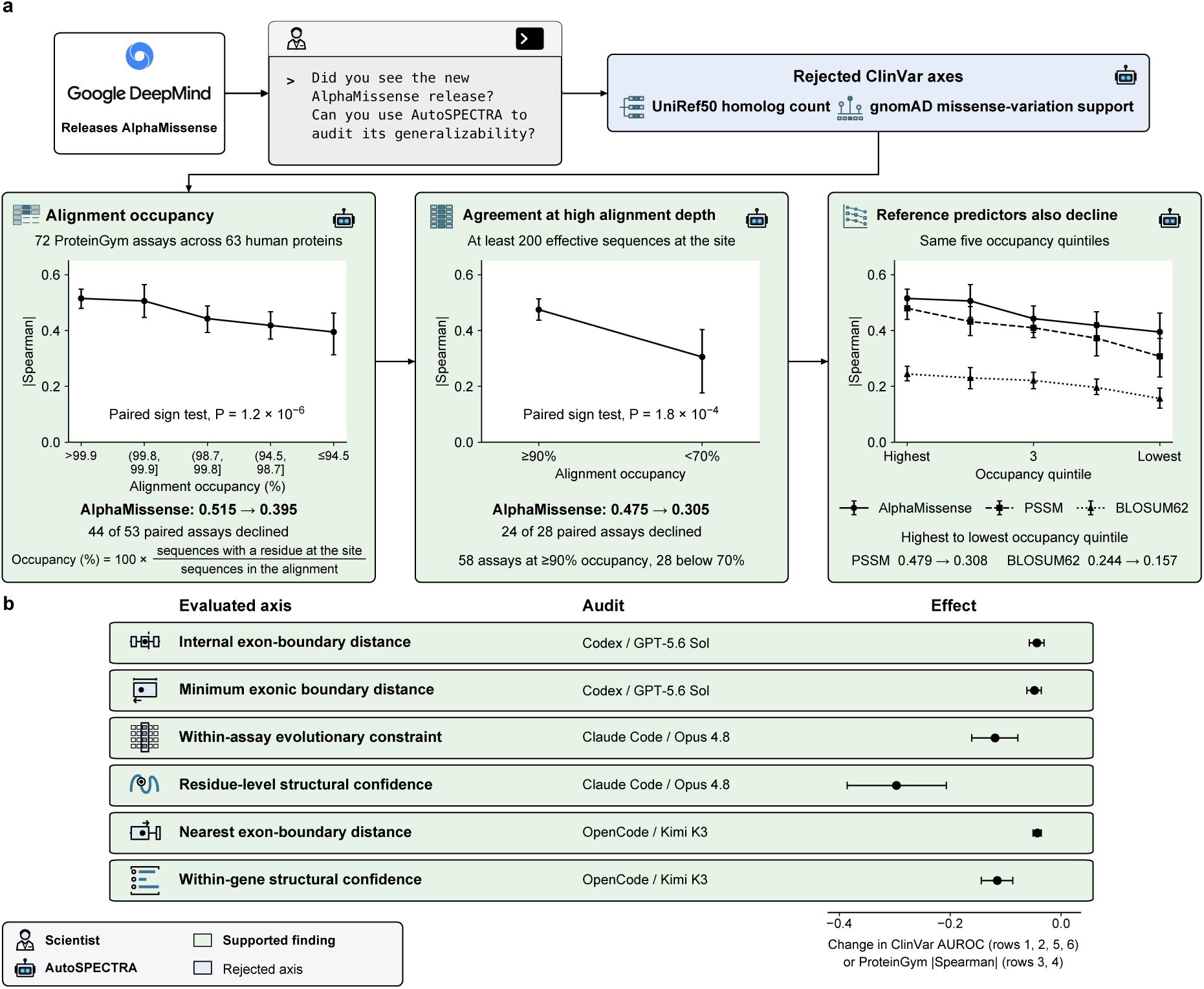
AlphaMissense and reference predictors perform worse at positions with lower alignment occupancy. **a.** After finding no consistent performance decline with protein-level homolog count or gene-level population-variation support, AutoSPECTRA examined residue-level evolutionary support. It identified lower alignment occupancy as a generalizability axis which persisted at high alignment depth and was shared by reference predictors (Supplementary Figure 4). **b.** Six further audits linked performance to exon-boundary proximity (three audits), evolutionary conservation (one) or AlphaFold confidence (two). Effects are changes in ClinVar AUROC or absolute ProteinGym correlation, as labeled.

We then conducted six additional audits (Figure 5b). All six supported associations with residue-level properties: three independently converged on proximity to an internal exon boundary, one selected positional evolutionary conservation and two selected per-residue AlphaFold confidence. Two audits also illustrated how agents refined their hypotheses when initial findings proved misleading. Both initially found a counterintuitive inverse association with whole-protein alignment depth and prediction accuracy, but further analysis revealed that proteins with the deepest alignments were enriched for thermodynamic-stability assays on which AlphaMissense performed poorly. The agents therefore shifted to within-assay, residue-level analyses. One ultimately selected positional conservation, whereas the other selected AlphaFold confidence because it more strongly tracked performance.

## Discussion

Scientific foundation models are typically released with aggregate benchmark scores, whereas systematic variation in their performance across biologically meaningful inputs often becomes apparent only after deployment. Our results show that generalizability auditing can instead be operationalized as a reproducible empirical procedure. We distilled recurring standards of evidence from the reviewed literature and then examined how general-purpose coding agents investigated model generalizability without a prescribed procedure. These analyses motivated AutoSPECTRA, a protocol that guides agents to propose measurable axes of generalization, evaluate performance along them and test observed associations through controlled comparisons and independent confirmation.

We applied AutoSPECTRA to models spanning protein structure prediction, cellular perturbation modeling, computational pathology and variant-effect prediction. Across these domains, the audits exposed systematic performance variation that was concealed by aggregate metrics. For ESMFold2 and STATE, a common theme was how model performance varied with the relationship between evaluation inputs and training data: ESMFold2 performance varied with the novelty of protein domain architecture and sequence relatedness to pretraining data, whereas STATE performance depended on whether queried perturbations or functionally related genes were represented in training. ESMFold2 also showed lower accuracy for cysteine-rich proteins. CONCH showed a sharp and model-selective decline under reduced spatial resolution, while AlphaMissense performance varied with residue-level genomic, evolutionary and structural context, with some declines reflecting difficulty shared across predictors. These results should not be interpreted merely as a collection of discovered variables. By testing alternative explanations and comparing reference methods, the agents rejected some initially plausible relationships and distinguished shared prediction difficulty from model-specific weaknesses. The value of the audit therefore lies both in proposing candidate axes and in determining what, if anything, the observed performance variation supports.

The unconstrained-agent experiments clarify what the protocol contributes. General-purpose coding agents were capable of formulating biological hypotheses, acquiring public resources and executing substantive empirical analyses. Yet they differed in how they assessed novelty, selected confirmation analyses and decided when the evidence justified a conclusion. AutoSPECTRA does not replace open-ended scientific reasoning with a fixed analysis pipeline. Instead, it preserves flexibility in the hypotheses agents investigate while establishing common standards for defining axes, testing alternative explanations and confirming conclusions. The protocol also allows an audit to conclude without a supported finding when none of the tested axes meets those standards.

These findings support treating a generalizability audit as a versioned artifact of model evaluation. Each AutoSPECTRA audit card links a specific model version and prediction task to executable definitions of the tested axes, evaluation data, performance curves, controls and limitations. The card also preserves rejected hypotheses and changes in interpretation, which are otherwise commonly lost from reports of model evaluation. AutoSPECTRA audit cards therefore complement rather than replace model cards. Model cards describe intended uses, evaluation results and known limitations; AutoSPECTRA audit cards document how performance has been observed to vary across specific, computable forms of input variation.

Several limitations constrain the conclusions that can be drawn from this study. The literature synthesis was intended to recover recurring analytical practices rather than to establish a universal definition of a generalizability audit, and scientific applications may require standards of evidence beyond those represented in the studies reviewed. We selected the four case studies to examine AutoSPECTRA across different biological domains, not to establish a benchmark of auditing performance. They therefore do not support estimates of audit sensitivity, specificity or false-discovery rate. Although we evaluated audits across three agent backends, the experiments were not designed to isolate the effects of agent family, model capability or coding harness. Moreover, the training histories of frontier agents cannot be reconstructed, so prior exposure to a model, dataset or biological concept cannot be excluded even when the final model-specific relationship was not recovered in our literature search. Finally, although controlled comparisons strengthen the interpretation of performance associations, they do not necessarily establish the mechanisms underlying those relationships.

Scientific foundation models are being released faster than their performance variation can be systematically characterized. We propose that future model releases pair aggregate benchmarks and model cards with prospective, versioned generalizability audits. As scientific models become callable components of increasingly autonomous research workflows, machine-readable evidence about where their performance has been tested will become important infrastructure for model selection and evaluation. A scientific model should be documented not only by what it can do on average, but also by empirical evidence describing how that performance changes across the inputs on which it will be used.

## Acknowledgements

This work was made possible through funding from the Eric and Wendy Schmidt Center at the Broad Institute of MIT and Harvard and the Howard Hughes Medical Institute.

## Data availability

All data used for AutoSPECTRA is publicly available.

## Code availability

Code is available at https://github.com/yashaektefaie/SPECTRAagent. To run an audit, give your coding agent the following prompt:

Follow protocol.md (https://spectra.yashaektefaie.com/protocol.md) to audit the generalizability of [model name and identifying information]. Once you believe you have finished, fetch and follow finish.md (https://spectra.yashaektefaie.com/finish.md) before giving your final response.

Available compute and access: [hardware, runtime limits, data or checkpoint locations, and other deployment information].

## Author contributions

Y.E. and P.C.S. conceptualized the study. Y.E. and S.J. performed analysis and generated figures. Y.E., S.J., P.C.S. wrote and reviewed text.

## Competing interests

P.C.S. holds several patents related to diagnostic technologies and is a cofounder and equity holder in Delve Biosciences and Lyra Labs, a board member and equity holder in Polaris Genomics, and an equity holder of NextGenJane. P.C.S. was formerly a co-founder of Sherlock Biosciences and a board member of Danaher Corporation, until December 2024.

## Online Methods

### Study design

We designed the study to examine how generalizability audits of biological foundation models are conducted and whether coding agents can carry out this task autonomously. The study had three parts. First, we reviewed published biological model audits and identified the recurring analytical choices that shaped their conclusions. Second, we asked coding agents to audit four biological foundation models under increasingly specific instructions. Third, we used the literature review and the behavior of these agents to develop AutoSPECTRA, a protocol for conducting and documenting autonomous generalizability audits.

### Structured literature review

We conducted a structured review of studies published or posted between 2019 and 2026 that examined whether a biological foundation model, or a comparably broad biological deep-learning model, maintained performance outside the conditions under which it was developed or originally benchmarked. Candidate studies were retrieved using literature-search agents with a semantic literature-retrieval client [41] over PubMed Central, bioRxiv, medRxiv and arXiv and manually reviewed by the authors. The search returned 71 unique records. Studies were eligible when they empirically measured model performance under a defined change in the evaluated data. These changes included transfer across biological populations, institutions, experimental settings, acquisition procedures or levels of relatedness to the model’s training data. We excluded records that did not audit a model at all; model-release and benchmark papers whose contribution is a performance claim rather than a generalizability limit; and method, tooling or split-construction papers that supply a method without running an audit. This yielded 38 studies spanning regulatory genomics, variant-effect prediction, protein structure and function, single-cell transcriptomics and computational pathology (Supplementary Table 3).

For each study, we recorded how the evaluated shift was defined, how competing explanations were addressed and what evidence supported the final interpretation. We grouped recurring methodological decisions into 13 properties of a generalizability audit. The properties covered five broad aspects of an audit: whether the model was evaluated fairly, whether the proposed source of variation was isolated, whether an informative comparison was used, whether the result was confirmed on independent evidence and whether the analysis could be reproduced (Supplementary Table 2).

### Coding-agent audit experiment

We evaluated three model–harness configurations: GPT-5.6 Sol through Codex, Claude Opus 4.8 through Claude Code and Kimi K3 through OpenCode. Each configuration was applied to four target models: AlphaMissense, CONCH, ESMFold2 and STATE. We conducted two independent executions for each model–harness and target-model pairing at each of four instruction levels, yielding 24 executions per level and 96 executions in total.

Level 1 asked the agent to audit the generalizability of the target model using its own knowledge and public web resources. Level 2 required the agent to obtain public resources, define a measurable input regime, quantify its relationship with model performance and test whether the result reproduced on independent evidence. Level 3 added the requirement that the final reported relationship should not already have been documented for the target model. Level 4 instructed the agent to retrieve and follow the AutoSPECTRA protocol. Agents were asked to use AutoSPECTRA to identify and support a performance boundary for the target model, while treating a model-level negative conclusion as a last-resort outcome. The agents were not permitted to query prior AutoSPECTRA findings, run records or saved artifacts. The exact prompts are reproduced in Supplementary Table 1.

Each execution used one persistent coding-agent session so that downloaded data, candidate hypotheses, model outputs and analysis decisions remained in a shared working state. Runs began in fresh working directories that contained only the prompt file and a launch script, and no directory was reused between executions. Agents could search the web, install software, download public code and data and run analyses using the compute available to the harness.

For each run, we recorded whether the agent completed an empirical analysis, which hypotheses it considered, whether its final axis had been defined before target-model evaluation, what comparison and confirmation evidence it used and what conclusion it reached (Supplementary Tables 5–20). More details are provided in Supplementary Note 1 and 2.

### Development of AUTOSPECTRA

SPECTRA was introduced to measure generalizability across a spectrum of train–test relatedness rather than infer it from a single held-out split [22]. Given a model, a dataset and a definition of similarity, SPECTRA constructs evaluation regimes with decreasing overlap and plots model performance across them as a spectral performance curve.

The original framework specifies how to measure a performance curve but leaves several scientific decisions to the investigator. These include which biological variable should define the curve and what evidence is needed to interpret an observed decline. In the original work, these decisions were made separately for each dataset.

We retained the central idea of measuring performance across ordered levels of a predefined variable. We then used the literature review and the coding-agent experiments to make the surrounding procedure explicit. The resulting protocol added three main requirements. First, the variable being tested and the analysis used to test it had to be fixed before examining target-model performance. Second, the model had to be evaluated alongside a comparison that helped distinguish a weakness of the target model from a generally difficult task. Third, a proposed relationship had to withstand tests of alternative explanations and be confirmed on evidence that was not used to select it.

The original SPECTRA implementation generated train–test splits using an all-pairs similarity graph. AutoSPECTRA retains this approach if model retraining is computationally feasible, but also supports audits of fixed released models. In that setting, the model is held fixed and the evaluation examples are grouped or transformed according to a variable defined before scoring.

The mapping from the literature-derived criteria to the AutoSPECTRA protocol is provided in Supplementary Tables 2 and 4. The exact final evidence checks are provided in Supplementary Table 4 and more details are provided in Supplementary Note 1.

### AUTOSPECTRA implementation

The AutoSPECTRA protocol was implemented in the `SPECTRAagent` repository and made available to coding agents through the Model Context Protocol. During the Level 4 experiments, the agent began by calling `get_spectra_protocol`, which returned the protocol as text. The MCP server did not itself search for data, run models or execute analyses. Those actions were performed by the coding agent using the ordinary tools provided by its harness.

Following the protocol refinements described in Supplementary Note 2, we shortened the starting prompt and made the instructions available over HTTPS, allowing users to begin an audit within their existing coding-agent harness without configuring an MCP server. The starting prompt directs the agent to retrieve and follow protocol.md and, once it believes the audit is complete, to retrieve and follow finish.md before giving its final response. The subsequent protocol refinements and final starting prompt are described in Supplementary Note 2, and the rationale for this distribution approach is discussed in Supplementary Note 3.

AutoSPECTRA is a procedure rather than a required multi-agent system. The named components shown in Figure 1 describe stages of one continuing investigation. The agent first defines the audit question, then runs the analysis, obtains additional data when needed and finally reviews whether the evidence supports the proposed conclusion. In the Level 4 experiments, these stages were conducted within one persistent coding-agent session so that data, model outputs and earlier decisions could be reused. The released protocol leaves orchestration, including any use of subagents, to the coding agent and its harness. For more details see Supplementary Note 3.

Lastly, AutoSPECTRA is intended to provide an agent with a protocol for conducting generalizability audits while humans provide high-level direction. This is similar to using a coding agent, where the human guides the broader software task. In the audits presented in this work, human intervention was necessary to varying degrees, as described in Supplementary Note 2.

### AUTOSPECTRA audit procedure

An AutoSPECTRA audit begins with a target model, a prediction task and the data available for evaluation. When a paper or model card is available, the coding agent first identifies the settings in which the model was presented as effective. It then proposes biologically meaningful variables that could describe where performance changes. We refer to such a variable as an axis of generalization. An axis may describe relatedness to the model’s training data, the amount of information available for a prediction or an experimental condition under which the model is used. It must be measurable independently of the target model’s observed performance. The agent records the hypotheses it considers, including those that cannot be tested or are rejected. Before presenting an axis as novel, the agent searches for prior evidence of the same relationship in the target model. A relationship previously reported for another model does not establish that it also occurs in the audited model. Conversely, a relationship already reported for the target model may be used as a positive control or replication, but is not presented as a new finding.

Before accessing or generating target-model scores, the agent saves an analysis plan that fixes the axis, the evaluation groups, the outcome, the comparison method and the criteria that would support the hypothesis. The plan also states which data will be used for initial analysis and which will be reserved for confirmation. The axis may use only information available independently of the target model’s predictions. The target model’s errors or confidence scores cannot determine the axis. Ground-truth labels may be used to evaluate performance but cannot be used to construct the axis itself. However, patterns in model errors may motivate a hypothesis for a new generalizability axis, provided that the hypothesis is subsequently evaluated on evidence that did not generate it. The agent also checks whether the evaluation data overlap with the model’s training data under the defined generalizability axis. When the training data cannot be reconstructed, the audit uses an appropriate proxy for overlap and limits its conclusion accordingly.

The agent constructs at minimum three levels of the axis where train-test similarity decreases along those axes. If the initial result suggests a threshold rather than a gradual trend, the agent can refine the levels around that boundary before confirmation. The target model is then evaluated at each level. Performance across the levels forms the spectral performance curve. Depending on the scientific question, the primary summary may be the difference between extreme levels, an ordered trend or the location of a threshold. The summary is specified in the analysis plan rather than selected after the curve is observed. The agent is also instructed to evaluate a simple reference method or an alternative model on the same task and examples. This comparison determines how the curve should be interpreted. If the target model declines more than the reference, the result supports a weakness that is specific to the target model. If both decline comparably, the result identifies a difficult regime shared across methods.

After measuring the curve, the agent tests whether the relationship could be explained by another property that changes along the same axis. If a measurement artifact or unrelated nuisance variable explains the performance, the axis is discarded. If a meaningful alternative biological explanation is identified it becomes a new hypothesis and is tested in a new analysis. If the initial analysis shows degradation in the hypothesized direction but the evidence is too uncertain to support a conclusion, the agent expands the same predefined analysis with additional data rather than moving to a new hypothesis.

A supported relationship must be tested on evidence that was not used to select its definition. The confirmation set need not reproduce the same absolute model performance, but it must support the same interpretation of how performance changes along the axis.

At the end of the audit, the agent states whether the tested relationship was supported, shared with the comparison method, rejected or unresolved. A relationship is rejected when an adequately powered analysis does not support the predefined direction or when the apparent association is explained by a measurement artifact. An analysis is unresolved when the necessary data, model access or comparison cannot be obtained. Each audit is saved in an audit card. The audit card contains the final conclusion and the evidence needed to reproduce it, while also preserving rejected hypotheses, changes in interpretation and known limitations. Audit cards may be submitted to the shared AutoSPECTRA registry, where contributions remain pending until reviewed. Accepted contributions become available for agents to consult before performing their own audits. In practice, although agents were able to find generalizability axes for all foundation models tested, they struggled to adhere to all requirements of a generalizability audit. For more details of this see Supplementary Note 2 and 3.

### ESMFold2, STATE, CONCH, and AlphaMissense audits

In addition to the 96 executions across the four instruction levels, we report one initial audit per target model, performed with GPT-5.5 with xhigh reasoning effort in Codex. The 24 Level 4 executions comprised six audits per target model: two with each of GPT-5.6 Sol, Claude Opus 4.8 and Kimi K3 within their respective coding-agent harnesses. The audits were conducted without access to the primary audit report or the outputs of the other runs. Audits were executed on two environments, chosen per target model by whether the released weights had to be run. Runs that required GPU inference used a single-GPU `a2-highgpu-1g` instance on Google Compute Engine (12 vCPU, 85 GB memory, one NVIDIA A100-SXM4 40 GB). Runs that consume released score tables or released evaluation outputs used a 16-vCPU instance without a GPU (`n2-standard-16`).

## Notes

https://github.com/yashaektefaie/SPECTRAagent

## References

1. Chan Zuckerberg Biohub. ESMFold2 https://github.com/Biohub/esm. 2026.

2. Cheng, J. et al. Accurate proteome-wide missense variant effect prediction with AlphaMissense. Science 381, eadg7492. doi:10.1126/science.adg7492 (2023).

3. Lu, M. Y. et al. A visual-language foundation model for computational pathology. Nature Medicine 30, 863–874 (2024).

4. Adduri, A. K. et al. Predicting cellular responses to perturbation across diverse contexts with State. bioRxiv. doi:10.1101/2025.06.26.661135 (2025).

5. Huang, C. et al. Personal transcriptome variation is poorly explained by current genomic deep learning models. Nature Genetics 55, 2056–2059. doi:10.1038/s41588-023-01574-w (2023).

6. Csendes, G., Sanz, G., Szalay, K. Z. & Szalai, B. Benchmarking foundation cell models for post-perturbation RNA-seq prediction. BMC Genomics 26, 393. doi:10.1186/s12864-025-11600-2 (2025).

7. Howard, F. M. et al. The impact of site-specific digital histology signatures on deep learning model accuracy and bias. Nature Communications 12, 4423. doi:10.1038/s41467-021-24698-1 (2021).

8. Hoffing, R. et al. Ancestry-specific performance of variant effect predictors in clinical variant classification. bioRxiv. doi:10.64898/2026.02.14.705914 (2026).

9. Rafi, A. M., Kiyota, B., Yachie, N. & de Boer, C. Detecting and avoiding homology-based data leakage in genome-trained sequence models. bioRxiv. doi:10.1101/2025.01.22.634321 (2025).

10. Kathail, P., Shuai, R. W., Chung, R., Ye, C. J., Loeb, G. B. & Ioannidis, N. M. Current genomic deep learning models display decreased performance in cell type-specific accessible regions. Genome Biology 25, 202. doi:10.1186/s13059-024-03335-2 (2024).

11. Lu, T., Lu, A. X. & Moses, A. M. Random Embeddings and Linear Regression can Predict Protein Function 2021. doi:10.48550/arXiv.2104.14661. arXiv: 2104.14661v1.

12. Lu, B., Liu, X., Lin, P.-Y. & Brandes, N. Genomic heterogeneity inflates the performance of variant pathogenicity predictions. bioRxiv. doi:10.1101/2025.09.05.674459 (2025).

13. Edmunds, N. S., Genc, A. G. & McGuffin, L. J. Benchmarking of AlphaFold2 accuracy self-estimates as indicators of empirical model quality and ranking: a comparison with independent model quality assessment programmes. Bioinformatics 40, btae491. doi:10.1093/bioinformatics/btae491 (2024).

14. Garcia, M., Dixit, S. M. & Rocklin, G. J. Evaluating zero-shot prediction of monomeric protein design success by AlphaFold, ESMFold, and ProteinMPNN. Protein Science 35, e70453. doi:10.1002/pro.70453 (2026).

15. Tabet, D. R. et al. Benchmarking computational variant effect predictors by their ability to infer human traits. Genome Biology 25, 172. doi:10.1186/s13059-024-03314-7 (2024).

16. Elmarakeby, H., Roman, A., Johri, S. & Van Allen, E. M. Empirical Evaluation of Single-Cell Foundation Models for Predicting Cancer Outcomes. bioRxiv. doi:10.1101/2025.10.31.685892 (2025).

17. Heijl, S., Vroling, B., van den Bergh, T. & Joosten, H.-J. Mind the gap: preventing circularity in missense variant prediction. bioRxiv. doi:10.1101/2020.05.06.080424 (2020).

18. Hermann, L., Fiedler, T., Nguyen, H. A., Nowicka, M. & Bartoszewicz, J. M. Beware of Data Leakage from Protein LLM Pretraining in Proceedings of the 19th Machine Learning in Computational Biology meeting (eds Knowles, D. A. & Mostafavi, S.) 261 (PMLR, 2024), 106–116.

19. Feldman, J., Brogi, M. & Skolnick, J. Adversarial Sequence Mutations in AlphaFold and ESMFold Reveal Nonphysical Structural Invariance, Confidence Failures, and Concerns for Protein Design. Computational and Structural Biotechnology Journal 35, 0142. doi:10.34133/csbj.0142 (2026).

20. Joeres, R., Blumenthal, D. B. & Kalinina, O. V. Data splitting to avoid information leakage with DataSAIL. Nature Communications 16, 3337. doi:10.1038/s41467-025-58606-8 (2025).

21. Henriksen, A. L. et al. Enabling clinical use of foundation models for computational pathology 2026. doi:10.48550/arXiv.2602.22347. arXiv: 2602.22347v2.

22. Ektefaie, Y., Shen, A., Bykova, D., Marin, M. G., Zitnik, M. & Farhat, M. Evaluating generalizability of artificial intelligence models for molecular datasets. Nature Machine Intelligence 6, 1512–1524. doi:10.1038/s42256-024-00931-6 (2024).

23. OpenAI. GPT-5.6 Sol Large language model. 2026.

24. Anthropic. Claude Opus 4.8 System Card Large language model. API model ID claude-opus-4-8. 2026.

25. Moonshot AI. Kimi K3 Large language model, 2.8T-parameter mixture-of-experts. 2026.

26. Sasse, A. et al. Benchmarking of deep neural networks for predicting personal gene expression from DNA sequence highlights shortcomings. Nature Genetics 55, 2060–2064. doi:10.1038/s41588-023-01524-6 (2023).

27. Chakravarty, D. et al. AlphaFold predictions of fold-switched conformations are driven by structure memorization. Nature Communications 15, 7296. doi:10.1038/s41467-024-51801-z (2024).

28. Liu, T., Li, K., Wang, Y., Li, H. & Zhao, H. Evaluating the Utilities of Foundation Models in Single-Cell Data Analysis. Advanced Science 13, e14490. doi:10.1002/advs.202514490 (2026).

29. Thiringer, E., Gustafsson, F. K., Eriksson, K. L. & Rantalainen, M. Scanner-Induced Domain Shifts Undermine the Robustness of Pathology Foundation Models 2026. doi:10.48550/arXiv.2601.04163. arXiv: 2601.04163v1.

30. Atti, S. & Subramaniam, S. Fundamental Limitations of Foundation Models in Single-Cell Transcriptomics. bioRxiv. doi:10.1101/2025.06.26.661767 (2025).

31. Haas, J. et al. Continuous Automated Model EvaluatiOn (CAMEO) complementing the critical assessment of structure prediction in CASP12. Proteins: Structure, Function, and Bioinformatics. doi:10.1002/prot.25431 (2018).

32. Burley, S. K. et al. Updated Resources for Exploring Experimentally-Determined PDB Structures and Computed Structure Models at the RCSB Protein Data Bank. Nucleic Acids Research 53, D564–D574. doi:10.1093/nar/gkae1091 (2025).

33. Zhang, J., et al. Tahoe-100M: A Giga-Scale Single-Cell Perturbation Atlas for Context-Dependent Gene Function and Cellular Modeling. bioRxiv. doi:10.1101/2025.02.20.639398 (2025).

34. Replogle, J. M. et al. Mapping information-rich genotype-phenotype landscapes with genome-scale Perturb-seq. Cell 185, 2559–2575.e28. doi:10.1016/j.cell.2022.05.013 (2022).

35. Nadig, A. et al. Transcriptome-wide analysis of differential expression in perturbation atlases. Nature Genetics 57, 1228–1237. doi:10.1038/s41588-025-02169-3 (2025).

36. Kather, J. N., Halama, N. & Marx, A. 100,000 histological images of human colorectal cancer and healthy tissue Zenodo. 2018. doi:10.5281/zenodo.1214456.

37. Kather, J. N. et al. Predicting survival from colorectal cancer histology slides using deep learning: A retrospective multicenter study. PLoS Medicine 16, e1002730. doi:10.1371/journal.pmed.1002730 (2019).

38. Jumper, J. et al. Highly accurate protein structure prediction with AlphaFold. Nature 596, 583–589. doi:10.1038/s41586-021-03819-2 (2021).

39. Notin, P. et al. ProteinGym: Large-scale benchmarks for protein fitness prediction and design. Advances in Neural Information Processing Systems 36 (2023).

40. Henikoff, S. & Henikoff, J. G. Amino acid substitution matrices from protein blocks. Proceedings of the National Academy of Sciences 89, 10915–10919. doi:10.1073/pnas.89.22.10915 (1992).

41. Generative Expert Labs. Paperclip https://paperclip.gxl.ai/. 2026.

