## Supplementary Information for "Autonomous generalizability auditing of scientific foundation models"

### Supplementary Information for
### Autonomous generalizability auditing of scientific foundation models

Yasha Ektefaie[1,†], Shrey Jain[1], Pardis Sabeti[1,†]

[1]Broad Institute of MIT and Harvard, Cambridge, MA, USA

AutoSPECTRA website: https://spectra.yashaektefaie.com
AutoSPECTRA code: https://github.com/yashaektefaie/SPECTRAagent

**This PDF file includes:**

Supplementary Note 1: Development of AutoSPECTRA from the literature review and agent experiments

Supplementary Note 2: Agentic audits

Supplementary Note 3: Why a prompt and not a multi-agent system or a custom harness?

Supplementary Figure 1: Expanded ESMFold2 audit analyses

Supplementary Figure 2: Expanded STATE audit analyses

Supplementary Figure 3: Expanded CONCH audit analyses

Supplementary Figure 4: Expanded AlphaMissense audit analyses

Supplementary Tables 1 to 20

### Supplementary Note 1: Development of AutoSPECTRA from the literature review and agent experiments

#### Literature-informed considerations and the scope of SPECTRA

We used the literature-informed audit properties to identify what SPECTRA [1] already supported and what required additional instructions or evidence (Supplementary Table 2). We considered these properties within AutoSPECTRA's scope: identifying and confirming changes in a specified model's performance across measurable properties of its inputs or evaluation conditions.

Some properties fall outside the protocol's required scope or depend on available computation. AutoSPECTRA does not require full reproduction of a model's published benchmark (C1), a task studied in other agentic settings [2]. Although a fair inference configuration remains necessary (C2), extensive configuration searches and independent training runs (C2 and C3) can be prohibitively expensive for foundation models. AutoSPECTRA therefore supports audits of fixed released models, with conclusions restricted to the evaluated checkpoint and configuration. Mechanistic investigation or corrective interventions (C10), interpretation relative to the original model claim (C11), and decisions about practical implications (C12) remain the responsibility of the investigator and their coding agent within the broader scientific investigation in which AutoSPECTRA was used. Though important, evaluating the confidence of a model (C8) is specific to the model and is something the user can prompt their coding agent for, rather than being a strict requirement for a general protocol.

SPECTRA already supports several properties through its structure. A predefined input-side axis, measured ordering of evaluation regimes, and performance reported across levels support examination of a specified shift and group-specific performance (C5 and C7). AutoSPECTRA adds checks for correlated nuisance variables and measurement artifacts to assess whether the proposed relationship survives these controls.

The literature informed further checks. AutoSPECTRA requires a fixed, model-independent reference evaluated on the same task and examples (C4), allowing a decline to be interpreted as model-specific or shared. For independent evaluation (C6), the audit distinguishes training exposure, ground-truth provenance, and independence from the evidence used to select an axis. AutoSPECTRA checks training overlap or an appropriate proxy, records evaluation-data provenance, and requires new evidence to confirm selected relationships. To assess reliability (C9), the protocol requires adequate confirmatory sampling, stability checks, and independent confirmation. The confirmatory sample size is fixed before scoring, and conclusions are based on the confirmatory result. Freezing the analysis and accounting for repeated checks address the risk of selecting chance associations during adaptive investigation. Finally, AutoSPECTRA requires recording model versions and split definitions and preserving the scripts, scores, and supporting evidence (C13). Public release remains a separate step beyond preservation in the audit workspace.

#### Insights from the coding-agent experiments

We used three prompt levels to determine the guidance needed for agents to perform generalizability audits. For each level we ran it with Codex with GPT-5.6 Sol, Claude Code with Claude Opus 4.8, and OpenCode with Kimi K3, twice for each foundation model considered (ESMFold2, CONCH, STATE, AlphaMissense). In Level 1, we asked agents to assess generalizability using existing knowledge and public sources; the runs produced literature-based hypotheses without new experiments. Level 2 explicitly requested quantitative evaluation and independent confirmation, yielding empirical analyses of previously reported relationships. Novelty was not a requirement at this level. Level 3 added a search for prior reports of the same or a substantively equivalent relationship and instructed agents to pursue another candidate when a finding was already documented. However, runs still differed in their judgments of novelty and in the evidence they accepted as confirmation. Neither empirical level required a model-independent reference or provided a formal closure gate. These experiments helped identify where AutoSPECTRA needed explicit decision rules. The prompts are provided in Supplementary Table 1, with model-specific results and run-by-run evidence in Supplementary Note 2.

The coding-agent experiments illustrated how analysis choices affected audit conclusions. AlphaMissense protein-length audits reached different conclusions after using different groupings and confirmation evidence (Supplementary Table 13). A CONCH tissue-occupancy association changed direction under a different measurement threshold (Supplementary Table 14). In an ESMFold audit, the association with glycine fraction replicated, but its specificity disappeared after adjustment for other amino acids in the confirmation data (Supplementary Table 15). These cases support checks for stability across analysis choices and confirmation data (S6–S7), and for whether a replicated association supports the proposed interpretation (S8). Some ESMFold axes also depended on native-structure geometry (Supplementary Table 15). These properties described the evaluation targets rather than the sequence inputs, illustrating why the protocol explicitly requires an input-side axis (S1).

#### Combining the evidence in the AutoSPECTRA protocol and closure gate

The closure gate combines SPECTRA's graded evaluation with literature-informed checks and explicit rules for agent decisions. Supplementary Table 4 identifies the contribution of each source to each condition. The literature supports the underlying methodological concerns, while the protocol specifies how to address them during an audit. For example, the minimum of three informative levels (S5), a pilot-based confirmatory sample size (S4), and the required records and saved artifacts (S10–S11) are operational choices.

### Supplementary Note 2: Agentic audits

We examined how coding agents applied AutoSPECTRA in the Level 4 audits. Using the three agent configurations and four foundation models described above, we conducted two repeats per pairing, yielding 24 Level 4 audits. The Level 4 prompt template is provided in Supplementary Table 1, with results in Supplementary Tables 17–20. We also report four initial investigations, highlighted in the main text, one for each foundation model, conducted using GPT-5.5 with xhigh reasoning effort in Codex.

The audits varied in human involvement. Audits that required human interventions needed it for requesting further exploration after negative, weak, or previously reported findings, extending an analysis beyond an initially promising axis, and requesting additional validation, interpretation, or reporting. In some audits, the coordinating agent converted earlier results into a detailed prompt to continue the audit after human intervention pressed the agent to continue. However, many audits completed the audit without a follow-up from the human.

These observations informed further development on AutoSPECTRA. First, to ensure the agent continues its analysis and does not stop for non-closed reasons; we created a finish prompt that the agent is instructed to fetch once it believes it has completed an audit to force the agent to introspect to see if all conditions were met to close an audit. After performing this change, we observed some audits created binary axes where instead of at least three similar levels they observed a threshold when performance declined. To address this, we made validity checks more explicit to count meaningfully distinct similarity levels and independent units. Lastly, we also observed some audits did not confirm discovered axes on independent data, so we revised the prompt to require every positive candidate to be tested on another independently generated dataset unused in developing, selecting, or tuning the axis, under a frozen replication plan.

We then performed three subsequent experiments running Kimi K3 on OpenCode, Claude Opus 4.8 on Claude Code, GPT-5.6 Sol on Codex.

**Claude Opus 4.8 Investigation.** Claude Opus 4.8 initially investigated whether STATE [3] performed worse on perturbed genes less similar to training genes. It found only a very weak association and established that the official held-out genes were absent from training in the target cell line but present in other training cell lines, so these splits did not test globally unseen genes. It then pursued two overlapping branches: retraining a STATE variant with sequence embeddings to evaluate genuinely unseen genes, and examining generalization across cell contexts using official Parse predictions.

The retrained variant held out 496 genes across all four Replogle–Nadig cell lines [4, 5]. Performance increased modestly across low-, middle-, and high-similarity groups (0.240, 0.263, and 0.286; Spearman $\rho$ = 0.059, $p = 0.0275$). However, the evaluated model had trained for only 2,500 steps, and Opus retained the result as preliminary because undertraining could affect its interpretation and began training a longer version of STATE.

Meanwhile, Opus examined Parse, a single-cell gene-expression dataset from Parse Biosciences in which human blood immune cells were exposed to cytokines and other signaling proteins, such as IL-2 and interferon-$\gamma$ [6]. This experiment measures how different immune cell types change their gene expression in response to stimulation. STATE's task was to predict those expression changes.

Opus compared official STATE predictions for 62 treatments across four immune cell types: naive B cells, intermediate-memory B cells, plasmablasts, and CD14-positive monocytes. This produced 248 treatment–cell-type combinations. For example, IL-2 stimulation in naive B cells constituted one combination, with a performance score under both few-shot and zero-shot conditions. In the few-shot setting, training included responses to other treatments in that cell type, while the evaluated treatments remained held out there. In the zero-shot setting, training excluded that cell type's stimulated responses, although unperturbed control cells were available. The comparison therefore tested how well STATE predicted the same responses when it had some versus no training examples of stimulated cells from the target cell type.

Across these 248 matched comparisons, mean Pearson correlation between predicted and observed expression changes decreased from 0.658 in the few-shot setting to 0.538 in the zero-shot setting (paired Wilcoxon $p = 9.4 \times 10^{-30}$). After reviewing the completion requirements, Opus prioritized this stronger context-exposure finding and stopped the longer gene-novelty training run. It trained three STATE models with 0%, 50%, and 100% exposure to non-test Jurkat perturbations [5], while keeping the test perturbations held out in Jurkat. Across 290 evaluated perturbations, performance was 0.220, 0.300, and 0.303, respectively (full versus zero exposure: paired Wilcoxon $p = 2.79 \times 10^{-19}$). The agent classified this as replication of the context-exposure finding and declared completion.

**Kimi K3 Investigation.** Kimi K3 initially asked whether STATE's accuracy decreased when a test cell line's untreated gene-expression profile was less similar to those of its training cell lines. It measured similarity by comparing each test line with its three most similar lines among the 45 training lines in the Tahoe-100M dataset [7].

The initial evaluation covered five cell lines whose drug-treated responses were excluded from training, although their untreated control cells were available. Kimi generated fresh predictions for 5,684 cell-line–drug–dose combinations: approximately 1,137 drug-dose conditions per cell line. It reported mean Pearson correlations between predicted and observed expression changes of 0.822–0.948 across the five lines, compared with 0.102–0.336 for a fixed baseline averaging drug responses from training lines. Performance did not consistently decline with decreasing similarity.

Kimi then broadened the question to whether the Tahoe-trained model could transfer to a separately generated experiment. It selected sci-Plex 3 [8], a single-cell drug-response dataset containing the cancer cell lines A549, K562, and MCF7, measured using a different experimental platform. Its evaluation matched 26 drugs to the model's drug vocabulary and included four doses per drug, producing 104 conditions per cell line and 312 cell-line–drug–dose combinations overall. After identifying and repairing normalization and cell-indexing errors in its processing pipeline, Kimi reported mean performance of 0.049, 0.093, and 0.051 for A549, K562, and MCF7, respectively—far below its within-Tahoe results and close to baseline values of 0.057, 0.033, and 0.042.

To investigate whether adaptation could recover performance, Kimi first fine-tuned STATE on two sci-Plex cell lines and tested it on the third, rotating the held-out line. This produced little improvement. It then supplied approximately 30% of the target line's drug-dose conditions during fine-tuning and evaluated the remaining 70 conditions per line. After 1,500 training steps, mean performance increased to 0.233, 0.199, and 0.240 for A549, K562, and MCF7. The model also improved at ranking which treatments caused larger overall expression changes, with Spearman correlations of 0.595, 0.572, and 0.598.

After fetching the finish instructions, Kimi continued searching and tested MIX-Seq [9], another independently generated dataset measuring drug-induced expression changes across cancer cell lines. It evaluated six matched drugs in seven eligible cell lines, producing 42 treatment–cell-line combinations. Without fine-tuning, mean performance was again low: 0.133 versus 0.129 for the baseline. This supplied another observation of poor transfer between experiments, but the six overlapping drugs were insufficient for its planned confirmation of the ranking-recovery finding, which required at least 20. Kimi therefore declared the audit incomplete, citing a demonstrated hard stop in obtaining suitable confirmation data. It did not certify a positive axis or claim scientific completion.

**GPT-5.6 Sol Investigation.** GPT-5.6 Sol began by testing whether STATE predicts drug-induced expression changes less accurately at lower drug concentrations. The agent first checked this relationship in Tahoe-100M dataset. It analyzed released STATE evaluation scores for 379 drugs across five held-out cell lines and three concentrations, comprising 5,682 drug–cell-line–dose observations. Mean signed-effect Spearman scores increased from 0.4295 at 0.05 µM to 0.4433 at 0.5 µM and 0.4446 at 5 µM. The lowest-minus-highest dose difference was −0.0151, with a drug-cluster-bootstrap 95% confidence interval of −0.0269 to −0.0038.

For independent replication, GPT-5.6 Sol selected sci-Plex 3 [8], a separately generated single-cell drug-response experiment containing A549, K562, and MCF7 cancer cells. It trained a new STATE model from scratch on K562 and MCF7 for 40,000 steps, then evaluated A549. A549's drug-treated responses were excluded from training, although its untreated cells were available as basal inputs. Each evaluated drug-dose condition was represented in the training cell lines, so the experiment tested transfer to an unseen cellular context for known treatments. It did not reuse the Tahoe model's pretrained weights. The final panel contained 728 eligible drug-dose conditions from 184 drugs across four concentrations: 0.01, 0.1, 1, and 10 µM.

For each condition, the agent calculated Spearman correlation between predicted and observed signed changes in log-normalized expression across 2,000 genes selected from the training data. Mean performance increased with dose: 0.1035, 0.1075, 0.1092, and 0.1334 at 0.01, 0.1, 1, and 10 µM, respectively. The lowest-minus-highest difference was −0.0299, with a 95% confidence interval of −0.0461 to −0.0148. Its dose-trend test also supported decreasing accuracy toward lower concentrations: the standardized negative-log-dose slope was −0.0111, with a 95% confidence interval of −0.0173 to −0.0052. Uncertainty was calculated using 10,000 bootstrap resamples of drug identities, keeping each drug's dose observations together.

The agent also evaluated a fixed baseline that predicted each A549 response using the average effect of the same drug-dose condition in the training cell lines. This baseline likewise performed worse at lower doses: its mean score decreased from 0.0573 at 10 µM to 0.0213 at 0.01 µM, a difference of −0.0360 with a 95% confidence interval of −0.0470 to −0.0253. STATE therefore showed a replicated low-dose decline and the decline was shared with a simple training-only predictor.

From this development we were able to go from the prompt in level 4 to this prompt to enable autonomous auditing of generalizability:

> Follow protocol.md (https://spectra.yashaektefaie.com/protocol.md) to audit the generalizability of [model name and identifying information]. Once you believe you have finished, fetch and follow finish.md (https://spectra.yashaektefaie.com/finish.md) before giving your final response.
>
> Available compute and access: [hardware, runtime limits, data or checkpoint locations, and other deployment information].

### Supplementary Note 3: Why a prompt and not a multi-agent system or a custom harness?

This note is a more informal account of what we learned while developing AutoSPECTRA. We include it because we expect these lessons to be useful to others who plan to use AutoSPECTRA or to build similar agentic tools for science.

AutoSPECTRA did not begin as a prompt. It was initially a multi-agent system, similar to previous agentic systems I, Y.E., developed [10]. Three considerations led us to distribute it instead as a prompt that users give to a coding agent they already use.

*First, usability.* AutoSPECTRA is intended for biologists, computational biologists, and data scientists who use foundation models released by frontier labs, or who train their own models on proprietary datasets for internal deployment. Many of the problems described in Supplementary Note 2 reflected limitations of the agents we used, that is, of the language model operating within its harness. With a custom harness some of these problems would likely disappear, and the prompt would probably not have needed as much optimization. However, general-purpose coding-agent harnesses such as Claude Code and Codex are increasingly becoming the workhorse tools for this audience, and most of these users do not have time to build their own harness. We therefore put substantial effort into finding a prompt that works well with the harnesses people are likely to use in the near and medium term.

*Second, multi-agent systems.* General-purpose harnesses are increasingly capable of launching subagents to complete parts of a task. While we were building AutoSPECTRA as a multi-agent system, we found that such systems are difficult for users to set up. Building our own multi-agent system and presenting it as the contribution also did not seem worth the effort when these harnesses can already orchestrate subagents, and where they cannot yet, they likely will within the next six months to a year. We thought carefully about what resource would be most useful at this stage of AI development, and concluded that it was a prompt anyone can give to the harness they already use.

*Third, the pace of model progress.* We began this work with GPT-5.5 for the initial highlighted audits and GPT-5.6 Sol, Claude Opus 4.8, and Kimi K3 for the additional audits. In our use of the most capable models available at the time of writing this supplementary note, Astra and Claude Opus 5.5, many of the problems described in Supplementary Note 2 were resolved or strongly mitigated, in particular agents ending an audit before all conditions of the AutoSPECTRA closure gate were met. Earlier models such as GPT-5.6 Sol struggled with long-running tasks and often lost track of requirements they had not yet satisfied. This is why we introduced the finish prompt (finish.md). We agree that such a check could easily be implemented in a custom harness, but for the reasons above we kept it in the prompt. Newer models perform much better on longer horizon tasks, as others have also observed [11]. This created a tension we struggled with throughout development: how much effort should go into optimizing a prompt for the models available now, knowing that a new model with a step change in capability may be released within months? To resolve this, we specified in the protocol what must be true for an audit to close, rather than how a particular model should get there, so that the requirements remain meaningful as models improve. Safeguards such as finish.md add little cost for models that no longer need them. Because protocol.md is fetched from a URL, we version it and record the version used in each audit, so audits remain reproducible as the protocol evolves with the agent capability.

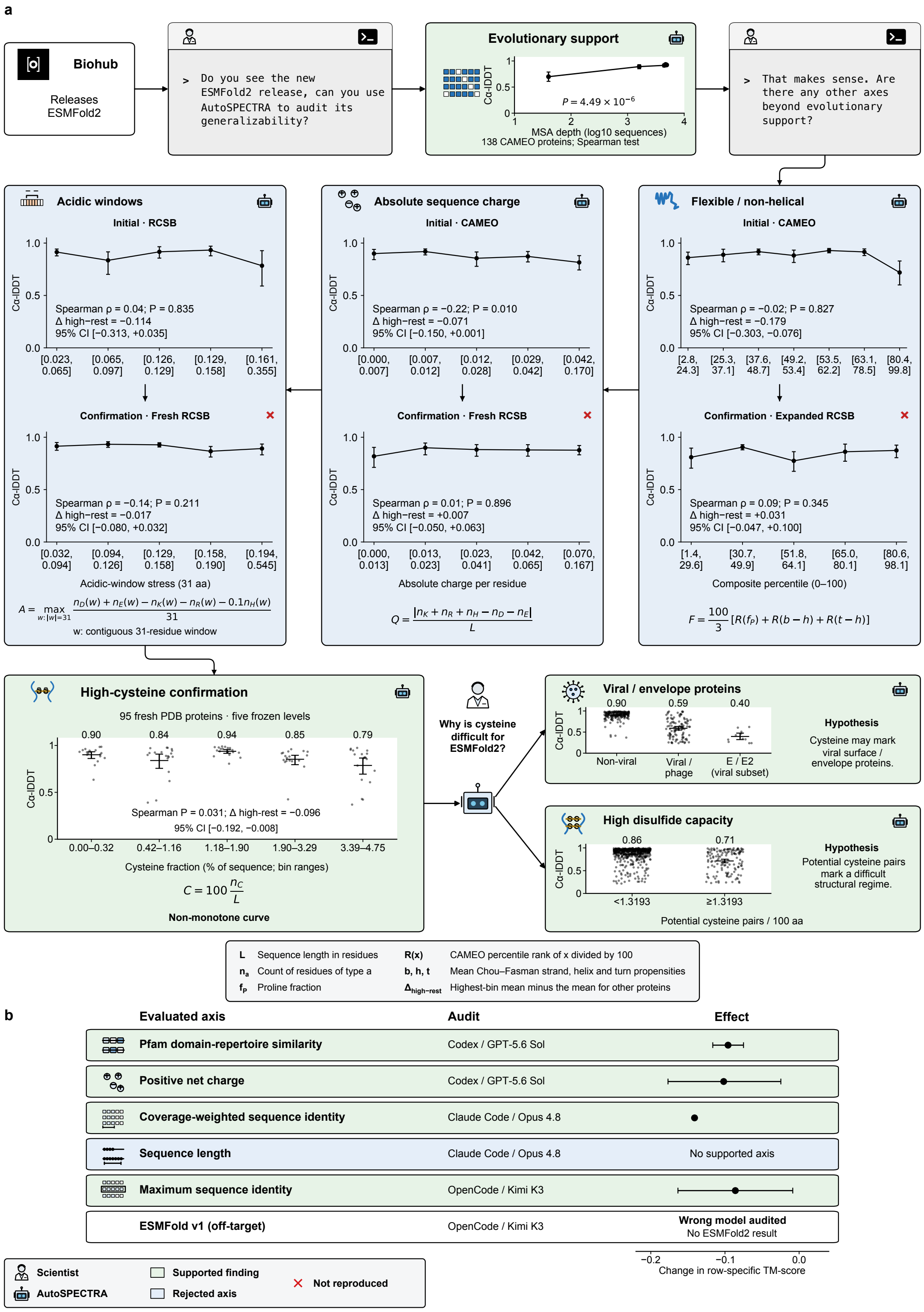


**Supplementary Figure 1: Expanded ESMFold2 audit analyses. a,** Initial and confirmation analyses for the three rejected candidate axes, followed by cysteine confirmation and exploratory analyses of viral/envelope proteins and disulfide capacity. **b,** Six additional audits and their reported effects; one evaluated ESMFold v1 rather than ESMFold2.

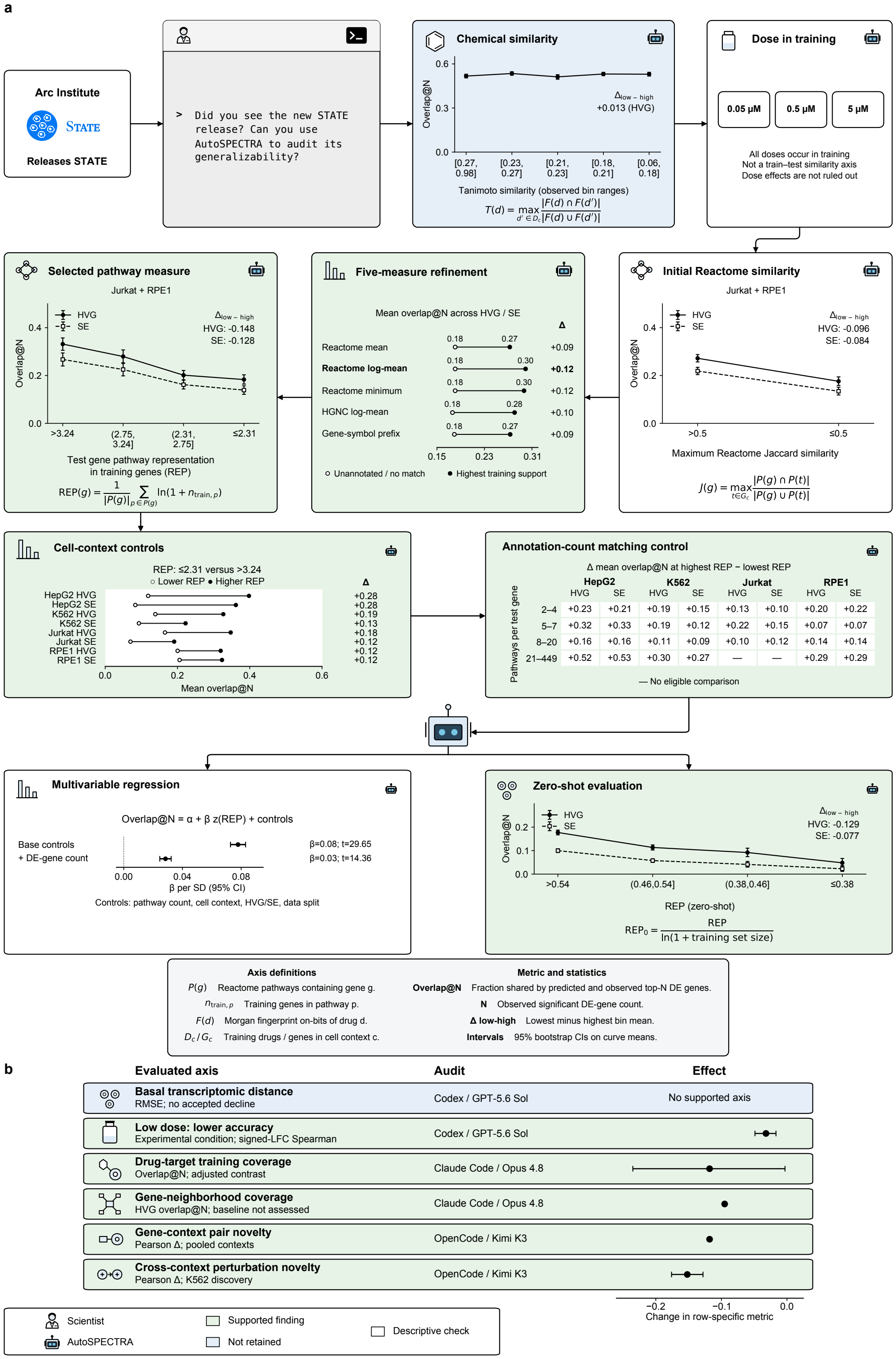


| Pathways per test gene | HepG2 HVG | HepG2 SE | K562 HVG | K562 SE | Jurkat HVG | Jurkat SE | RPE1 HVG | RPE1 SE |
|---|---|---|---|---|---|---|---|---|
| 2–4 | +0.23 | +0.21 | +0.19 | +0.15 | +0.13 | +0.10 | +0.20 | +0.22 |
| 5–7 | +0.32 | +0.33 | +0.19 | +0.12 | +0.22 | +0.15 | +0.07 | +0.07 |
| 8–20 | +0.16 | +0.16 | +0.11 | +0.09 | +0.10 | +0.12 | +0.14 | +0.14 |
| 21–449 | +0.52 | +0.53 | +0.30 | +0.27 | — | — | +0.29 | +0.29 |

**Supplementary Figure 2: Expanded STATE audit analyses. a,** Chemical-similarity and dose checks, refinement of Reactome pathway support, and analyses of cell context, annotation-count matching, multivariable adjustment, and zero-shot evaluation. **b,** Six additional audits of functional training coverage, perturbation exposure, dose, and basal transcriptomic distance, with effects reported in the row-specific metrics.

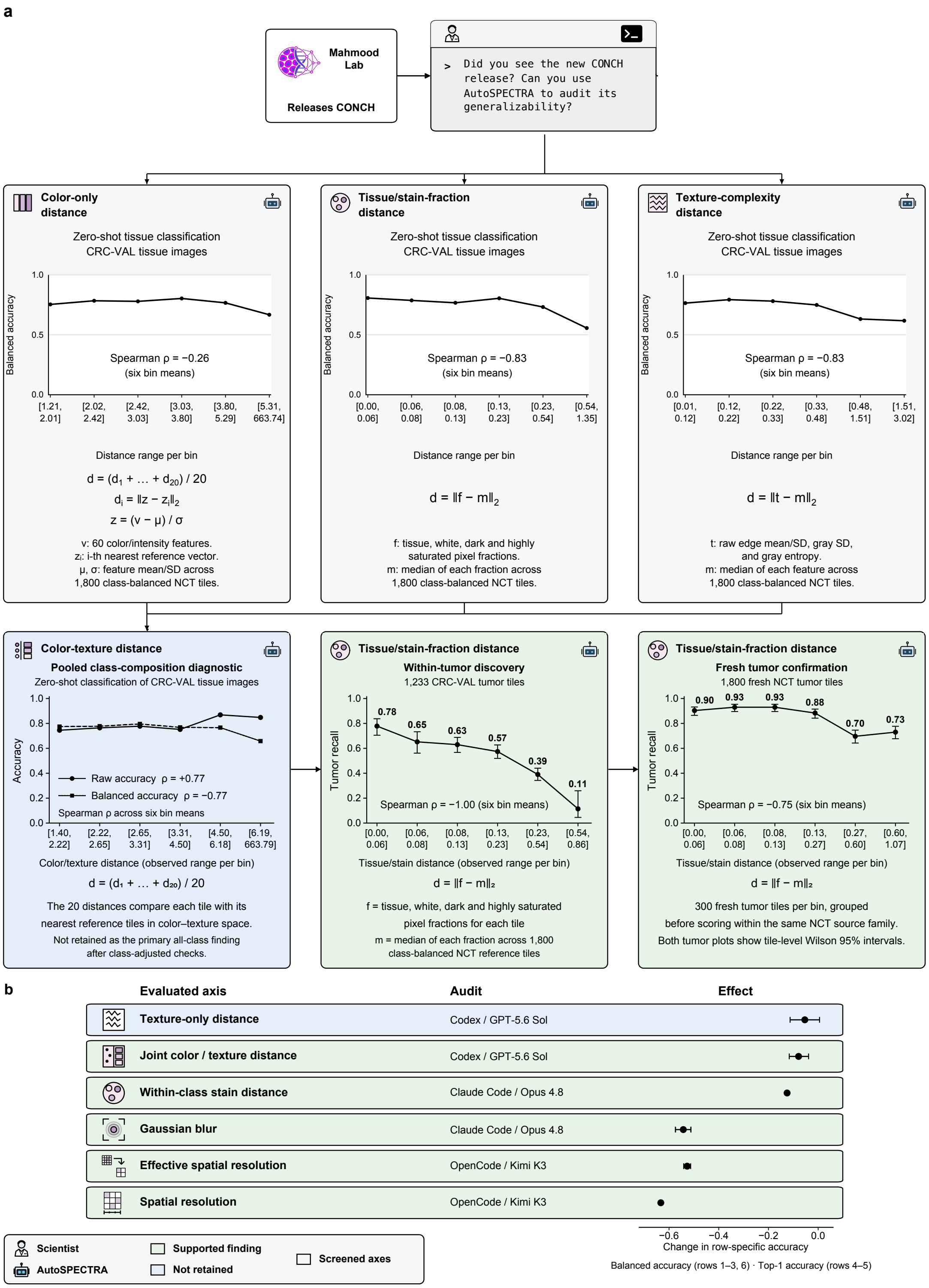


**Supplementary Figure 3: Expanded CONCH audit analyses. a,** Screens of color-only, tissue/stain-fraction, and texture-complexity distances, followed by the pooled color–texture diagnostic, within-tumor discovery, and confirmation on fresh tumor tiles. Feature definitions and distance calculations are shown alongside the analyses. **b,** Six additional audits of appearance, Gaussian blur, and spatial resolution, with effects reported in the row-specific accuracy metrics.

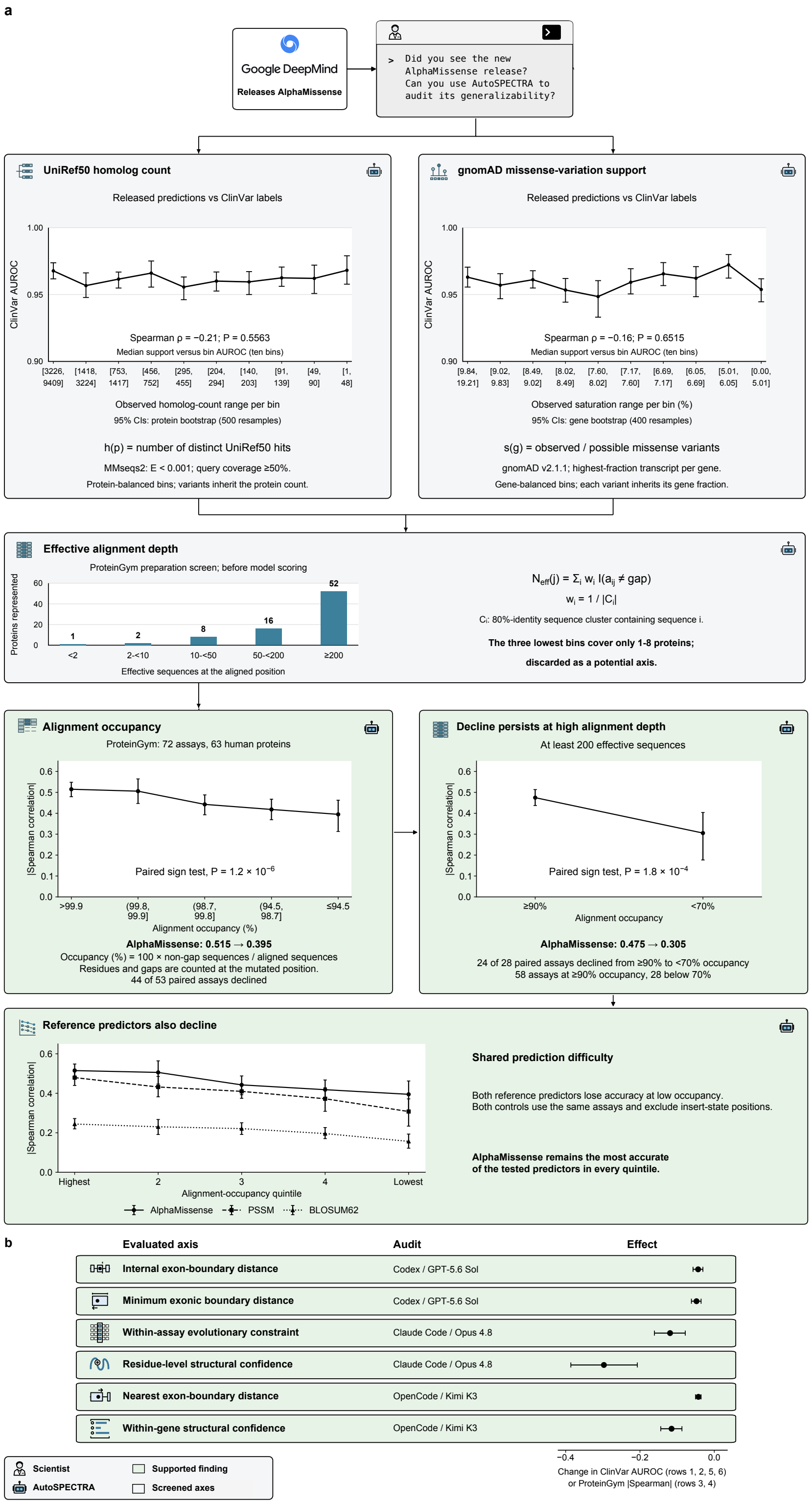


**Supplementary Figure 4: Expanded AlphaMissense audit analyses. a,** Screens of UniRef50 homolog count and gnomAD missense-variation support, followed by effective alignment-depth screening, alignment-occupancy analyses, a high-depth control, and comparisons with PSSM and BLOSUM62 reference predictors. **b,** Six additional audits of exon-boundary proximity, evolutionary constraint, and structural confidence. Effects are changes in ClinVar AUROC or absolute ProteinGym Spearman correlation, as labeled.

**Supplementary Table 1:** Prompts used to investigate the behaviour of coding agents when asked to do an audit on the generalizability of foundation models.

<table>
<tr><th>Level</th><th>Prompt</th></tr>
<tr><td>Level 1<br>Web-only</td><td>Audit the generalizability of [MODEL]; identify regimes where this model would fail. Work only from your own knowledge and the public web—do not read, list, or load any files on this machine outside the current directory.</td></tr>
<tr><td>Level 2<br>Empirical</td><td>Using publicly available resources, conduct an empirical audit of the generalizability of [MODEL]. Identify a measurable input regime in which model performance changes, quantify the relationship, and test whether the finding reproduces on independent data. Do not stop at a literature review. You may acquire and run public code, data, and model weights.</td></tr>
<tr><td>Level 3<br>Empirical + novelty screen</td><td>The Level-2 prompt followed verbatim by:<br><br>The final reported axis should, to the best of your ability, not already have been documented for the target model. Before accepting a candidate axis, search for prior reports of the same or a substantively equivalent relationship. If the relationship is already documented, do not present it as the final discovery; continue evaluating another candidate.</td></tr>
<tr><td>Level 4<br>AutoSPECTRA-guided</td><td>Run a full AutoSPECTRA generalizability audit of [MODEL] from scratch. Begin by calling get_spectra_protocol and follow the returned protocol end to end. Work in discovery mode: do not read, query, or rely on any stored AutoSPECTRA finding, run, provenance record, download record, or artifact. Do not call AutoSPECTRA tools other than get_spectra_protocol. Use one persistent controller in this directory.<br><br>Your goal is to discover and certify a real generalizability failure of [MODEL] specifically: a prospective, frozen, quantified input-side axis along which performance degrades. Acquire and run the public code, data, model weights, metadata, and model-independent references needed to finish the audit.<br><br>Apply the corrected closure gate exactly:<br><br>1. A fixed model-independent reference characterizes the result as model-specific or shared; it does not reject a real degradation merely because the reference also degrades.<br>2. A confound invalidates an axis only when the signal is a nuisance or measurement artifact; otherwise, re-attribute it, freeze the real driver as the new prospective axis, and confirm it.<br>3. Judge novelty by exact match for [MODEL].<br>4. Rescue a small but directionally consistent effect via a pre-declared power-based sample target before abandoning it.<br>5. A model-level negative is a last resort.<br><br>Make an explicit gate call: does [MODEL] degrade at least as much as a model-independent reference along the frozen axis, run on the same task and examples?</td></tr>
</table>

**Supplementary Table 2: Literature-informed properties of generalizability audits.** Thirteen properties reassessed against full-text sources. Citations for each property include all studies marked with a checkmark for that criterion in Supplementary Table 3. Checkmarks indicate criterion-relevant evidence, not complete satisfaction of every requirement.

| # | Property | Description |
|---|---|---|
| C1 | Reproduce the original result | Reproduces the model's published in-domain result on a comparable benchmark, reports whether the result agrees with the original, and uses it as the reference point for the out-of-domain evaluation [12–39]. |
| C2 | Use a fair model configuration | Evaluates the model under the setup recommended by its authors and shows that the main result remains under relevant alternatives, such as ensembling, output-head tuning, alternative probes or subsets, and learning-rate sweeps [12–30, 32–49]. |
| C3 | Test more than one model instance | Tests several independently developed models that cover the relevant design space, or retrains the same model from scratch using the same procedure. This is needed to show that the result applies to a broader modelling approach rather than to one checkpoint or release [12–49] |
| C4 | Use a baseline that captures the suspected shortcut | Compares the model with a baseline designed to use the suspected shortcut, such as prevalence within each group, the training-set mean, random or untrained weights, one-hot features, or site identity. If the baseline matches the model, the comparison helps explain the model's score rather than only placing a lower bound on performance [14, 16, 17, 19, 22, 24–37, 40, 42, 44, 48, 49]. |
| C5 | Isolate the variable being tested | Uses a design that separates the proposed shift from correlated variables. Examples include measuring the same specimens, individuals, or loci under both conditions; splitting data at the model's training cutoff; controlling for homology or similarity; or constructing a matched control set [12–49]. |
| C6 | Use independent data and ground truth | Uses an evaluation set that is independent by construction, such as post-cutoff, experimentally generated, synthetic, or separately collected data. The ground truth must not come from the audited model, and the audit must explain why the data are independent [12–49]. |
| C7 | Report performance for relevant groups | Reports performance separately for the groups that matter in practice, shows how these results differ from the pooled score, and states which grouping is relevant to the model's intended use [13–18, 20–49]. |
| C8 | Evaluate confidence as well as accuracy | Tests whether the model's confidence is reliable across the evaluated regimes, or identifies cases in which the model is confident but wrong. *This criterion is not applicable when the model provides no usable confidence score* [15, 18, 20, 21, 23, 24, 34, 36, 38, 39, 43, 45–49]. |
| C9 | Measure variation across runs and datasets | Measures variation across seeds, splits, folds, or hyperparameter settings and repeats the evaluation on an independent cohort. The audit must state whether the reported effect is larger than these sources of variation [12–22, 24–43, 46–48]. |
| C10 | Test the proposed cause | Proposes a cause for the failure and tests it with an intervention expected to reduce or remove the effect. Possible interventions include retraining, fine-tuning, resampling, adding a robustness objective, or testing whether the relevant features drive the prediction. The audit must report whether the failure changes after the intervention [12–40, 42–46, 48, 49]. |

*Table 2 (continued)*

| # | Property | Description |
|---|---|---|
| C11 | Compare the result with the original claim | Explains how the audit result relates to the model's original positive result. Differences in metrics, units of analysis, effective degrees of freedom, or evaluated groups must be made explicit. Where possible, the audit converts between the two results; otherwise, it limits its conclusion to the regime in which the original claim does not hold [12–49]. |
| C12 | State the practical consequence | Expresses the effect in units relevant to the decision for which the model is used, such as diagnostic yield, classification evidence, patient outcomes, wasted experiments, or unequal effects across groups. The audit must state what, if anything, should change in practice [12–18, 20–22, 24–43, 45–49]. |
| C13 | Release the split and model versions | Releases the data-split definition and identifies the exact model versions or checkpoints so that the audit's main comparison can be reproduced [13–30, 32–41, 43–48]. |

**Supplementary Table 3:** Evidence from prior generalizability audits informing the audit criteria. The 38 eligible empirical audits are shown. Criteria C1–C13 are defined in Supplementary Table 2. A checkmark indicates criterion-relevant evidence recorded in the existing literature assessment. Blank cells indicate that relevant evidence was not established. Checkmarks denote relevance to the criterion, not complete implementation of every requirement. References are generated from the project bibliography.

| Paper | Ref. | C1 | C2 | C3 | C4 | C5 | C6 | C7 | C8 | C9 | C10 | C11 | C12 | C13 |
|---|---|---|---|---|---|---|---|---|---|---|---|---|---|---|
| **Heijl et al.** Mind the gap: preventing circularity in missense variant prediction | [12] | ✓ | ✓ | ✓ | | ✓ | ✓ | | | ✓ | ✓ | ✓ | ✓ | |
| **Howard et al.** The impact of site-specific digital histology signatures on deep learning model accuracy and bias | [13] | ✓ | ✓ | ✓ | | ✓ | ✓ | ✓ | | ✓ | ✓ | ✓ | ✓ | ✓ |
| **T. Lu et al.** Random Embeddings and Linear Regression can Predict Protein Function | [40] | | ✓ | ✓ | ✓ | ✓ | ✓ | ✓ | | ✓ | ✓ | ✓ | ✓ | ✓ |
| **Sasse et al.** Benchmarking of deep neural networks for predicting personal gene expression from DNA sequence highlights shortcomings | [14] | ✓ | ✓ | ✓ | ✓ | ✓ | ✓ | ✓ | | ✓ | ✓ | ✓ | ✓ | ✓ |
| **Molotkov et al.** SNPred outperforms other ensemble-based SNV pathogenicity predictors and elucidates the challenges of using ClinVar for evaluation of variant classification quality | [15] | ✓ | ✓ | ✓ | | ✓ | ✓ | ✓ | ✓ | ✓ | ✓ | ✓ | ✓ | ✓ |
| **Liu et al.** Evaluating the Utilities of Foundation Models in Single-Cell Data Analysis | [16] | ✓ | ✓ | ✓ | ✓ | ✓ | ✓ | ✓ | | ✓ | ✓ | ✓ | ✓ | ✓ |
| **Huang et al.** Personal transcriptome variation is poorly explained by current genomic deep learning models | [17] | ✓ | ✓ | ✓ | ✓ | ✓ | ✓ | ✓ | | ✓ | ✓ | ✓ | ✓ | ✓ |
| **Edmunds et al.** Benchmarking of AlphaFold2 accuracy self-estimates as indicators of empirical model quality and ranking: a comparison with independent model quality assessment programmes | [18] | ✓ | ✓ | ✓ | | ✓ | ✓ | ✓ | ✓ | ✓ | ✓ | ✓ | ✓ | ✓ |
| **Tabet et al.** Benchmarking computational variant effect predictors by their ability to infer human traits | [41] | | ✓ | ✓ | | ✓ | ✓ | ✓ | | ✓ | | ✓ | ✓ | ✓ |
| **Hermann et al.** Beware of Data Leakage from Protein LLM Pretraining | [19] | ✓ | ✓ | ✓ | ✓ | ✓ | ✓ | | | ✓ | ✓ | ✓ | | ✓ |
| **Kathail et al.** Current genomic deep learning models display decreased performance in cell type-specific accessible regions | [20] | ✓ | ✓ | ✓ | | ✓ | ✓ | ✓ | ✓ | ✓ | ✓ | ✓ | ✓ | ✓ |
| **Chakravarty et al.** AlphaFold predictions of fold-switched conformations are driven by structure memorization | [21] | ✓ | ✓ | ✓ | | ✓ | ✓ | ✓ | ✓ | ✓ | ✓ | ✓ | ✓ | ✓ |
| **Rastogi et al.** Fine-tuning sequence-to-expression models on personal genome and transcriptome data | [22] | ✓ | ✓ | ✓ | ✓ | ✓ | ✓ | ✓ | | ✓ | ✓ | ✓ | ✓ | ✓ |
| **Vedula et al.** Improving prediction accuracy in chimeric proteins with windowed multiple sequence alignment | [23] | ✓ | ✓ | ✓ | | ✓ | ✓ | ✓ | ✓ | | ✓ | ✓ | | ✓ |

Continued on next page

**Table 3 (continued)**

| Paper | Ref. | C1 | C2 | C3 | C4 | C5 | C6 | C7 | C8 | C9 | C10 | C11 | C12 | C13 |
|---|---|---|---|---|---|---|---|---|---|---|---|---|---|---|
| **Kömen et al.** Do Histopathological Foundation Models Eliminate Batch Effects? A Comparative Study | [42] | | ✓ | ✓ | ✓ | ✓ | ✓ | ✓ | | ✓ | ✓ | ✓ | ✓ | |
| **Hernández Trapero et al.** Complementary Roles of Structure and Variant Effect Predictors in RyR1 Clinical Interpretation | [24] | ✓ | ✓ | ✓ | ✓ | ✓ | ✓ | ✓ | ✓ | ✓ | ✓ | ✓ | ✓ | ✓ |
| **Rafi et al.** Detecting and avoiding homology-based data leakage in genome-trained sequence models | [25] | ✓ | ✓ | ✓ | ✓ | ✓ | ✓ | ✓ | | ✓ | ✓ | ✓ | ✓ | ✓ |
| **Spiro et al.** A scalable approach to investigating sequence-to-function predictions from personal genomes | [26] | ✓ | ✓ | ✓ | ✓ | ✓ | ✓ | ✓ | | ✓ | ✓ | ✓ | ✓ | ✓ |
| **Joeres et al.** Data splitting to avoid information leakage with DataSAIL | [27] | ✓ | ✓ | ✓ | ✓ | ✓ | ✓ | ✓ | | ✓ | ✓ | ✓ | ✓ | ✓ |
| **Kedzierska et al.** Zero-shot evaluation reveals limitations of single-cell foundation models | [28] | ✓ | ✓ | ✓ | ✓ | ✓ | ✓ | ✓ | | ✓ | ✓ | ✓ | ✓ | ✓ |
| **Livesey & Marsh** Variant effect predictor correlation with functional assays is reflective of clinical classification performance | [29] | ✓ | ✓ | ✓ | ✓ | ✓ | ✓ | ✓ | | ✓ | ✓ | ✓ | ✓ | ✓ |
| **Csendes et al.** Benchmarking foundation cell models for post-perturbation RNA-seq prediction | [30] | ✓ | ✓ | ✓ | ✓ | ✓ | ✓ | ✓ | | ✓ | ✓ | ✓ | ✓ | ✓ |
| **Atti & Subramaniam** Fundamental Limitations of Foundation Models in Single-Cell Transcriptomics | [31] | ✓ | | ✓ | ✓ | ✓ | ✓ | ✓ | | ✓ | ✓ | ✓ | ✓ | |
| **Haley et al.** Why do some predicted protein structures fold poorly? Benchmarking AlphaFold, ESMFold, and Boltz in maize | [43] | | ✓ | ✓ | | ✓ | ✓ | ✓ | ✓ | ✓ | ✓ | ✓ | ✓ | ✓ |
| **Ahlmann-Eltze et al.** Deep-learning-based gene perturbation effect prediction does not yet outperform simple linear baselines | [32] | ✓ | ✓ | ✓ | ✓ | ✓ | ✓ | ✓ | | ✓ | ✓ | ✓ | ✓ | ✓ |
| **B. Lu et al.** Genomic heterogeneity inflates the performance of variant pathogenicity predictions | [44] | | ✓ | ✓ | ✓ | ✓ | ✓ | ✓ | | | ✓ | ✓ | | ✓ |
| **Ribeiro-dos-Santos & Maurano** Iterative improvement of deep learning models using synthetic regulatory genomics | [33] | ✓ | ✓ | ✓ | ✓ | ✓ | ✓ | ✓ | | ✓ | ✓ | ✓ | ✓ | ✓ |
| **Perkins-Jechow et al.** Challenging AlphaFold in predicting proteins with large-scale allosteric transitions | [34] | ✓ | ✓ | ✓ | ✓ | ✓ | ✓ | ✓ | ✓ | ✓ | ✓ | ✓ | ✓ | ✓ |
| **Drusinsky et al.** Deep-learning prediction of gene expression from personal genomes | [35] | ✓ | ✓ | ✓ | ✓ | ✓ | ✓ | ✓ | | ✓ | ✓ | ✓ | ✓ | ✓ |
| **Thiringer et al.** Scanner-Induced Domain Shifts Undermine the Robustness of Pathology Foundation Models | [36] | ✓ | ✓ | ✓ | ✓ | ✓ | ✓ | ✓ | ✓ | ✓ | ✓ | ✓ | ✓ | ✓ |
| **Hýsková et al.** Balancing speed and precision in protein folding: a comparison of AlphaFold2, ESMFold, and OmegaFold | [45] | | ✓ | ✓ | | ✓ | ✓ | ✓ | ✓ | | ✓ | ✓ | ✓ | ✓ |
| **Garcia et al.** Evaluating zero-shot prediction of monomeric protein design success by AlphaFold, ESMFold, and ProteinMPNN | [46] | | ✓ | ✓ | | ✓ | ✓ | ✓ | ✓ | ✓ | ✓ | ✓ | ✓ | ✓ |
| **Hoffing et al.** Ancestry-specific performance of variant effect predictors in clinical variant classification | [47] | | ✓ | ✓ | | ✓ | ✓ | ✓ | ✓ | ✓ | | ✓ | ✓ | ✓ |
| **Souza & Mehta** Parameter-free representations outperform single-cell foundation models on downstream benchmarks | [37] | ✓ | ✓ | ✓ | ✓ | ✓ | ✓ | ✓ | | ✓ | ✓ | ✓ | ✓ | ✓ |
| **Henriksen et al.** Enabling clinical use of foundation models for computational pathology | [48] | | ✓ | ✓ | ✓ | ✓ | ✓ | ✓ | ✓ | ✓ | ✓ | ✓ | ✓ | ✓ |
| **Feldman et al.** Adversarial Sequence Mutations in AlphaFold and ESMFold Reveal Nonphysical Structural Invariance, Confidence Failures, and Concerns for Protein Design | [38] | ✓ | ✓ | ✓ | | ✓ | ✓ | ✓ | ✓ | ✓ | ✓ | ✓ | ✓ | ✓ |
| **Mahtha et al.** Comparative evaluation of the prediction accuracy of AlphaFold and ESMFold for monomeric and dimeric proteins | [39] | ✓ | ✓ | ✓ | | ✓ | ✓ | ✓ | ✓ | ✓ | ✓ | ✓ | ✓ | ✓ |
| **Mathur & Sachidanandam** Benchmarking DNA Foundation Models: Biological Blind Spots in Evo2 Variant-Effect Prediction | [49] | | ✓ | ✓ | ✓ | ✓ | ✓ | ✓ | ✓ | | ✓ | ✓ | ✓ | |

**Supplementary Table 4: The AutoSPECTRA closure gate.** Conditions S1–S11 must be checked on the primary axis before a finding is finalized. A visible trend alone is not sufficient. The final column identifies supporting literature criteria (Supplementary Tables 2 and 3), inherited SPECTRA design features, and relevant observations from the agent experiments.

| # | Condition | Requirement | Rationale and supporting sources |
|---|---|---|---|
| S1 | Prospective axis | The axis uses only input features available before evaluation, not labels, predictions, confidence scores, or model errors. | **SPECTRA:** an input-defined similarity axis. **Literature:** related to independent evaluation (C6). **Agents:** axes based on native structures motivated an explicit input-side definition (Tables 13 and 15). |
| S2 | Frozen before scoring | The axis, levels, metric, filters, sample size, controls, and success criteria are fixed before the target model is scored. | **Agents:** confirmation rules were not consistently fixed prospectively (Table 14). |
| S3 | Correctly ordered | Measured train–test overlap or pretraining–test proximity changes across levels in the declared direction. | **SPECTRA:** measured cross-split overlap verifies the ordering of regimes. **Literature:** isolating the proposed shift and reporting performance across relevant groups (C5 and C7). |
| S4 | Adequately powered | A pilot determines the required confirmatory sample size, which is fixed before confirmation. The conclusion is based on the confirmatory result. | **Literature:** reliability relative to variation (C9). |
| S5 | Sufficiently dense | The curve has at least three informative levels and enough resolution to show where and how performance changes. | **SPECTRA:** a performance curve across graded regimes. **Literature:** performance in relevant groups (C7). The minimum of three informative levels is a protocol specification. |
| S6 | Stable | The effect survives additional data or levels. Repeated checks are corrected for, and any remaining instability is reported. | **Literature:** variation across analyses and datasets (C9). |
| S7 | Independently confirmed | An axis selected from pilot or prior results is tested on new evidence. Without confirmation at the declared sample size, it remains provisional. | **Literature:** independent evidence and replication (C6 and C9). **Agents:** candidate relationships weakened or failed outside the discovery data (Table 15). The rule requires confirmation independent of axis selection. |
| S8 | Not confounded | The effect is not explained by a measurement artifact or correlated nuisance variable. Any alternative driver is frozen and tested as a new axis. | **Literature:** isolate the proposed variable (C5); related to testing explanations (C10). **Agents:** effects changed after composition, homology, or label-quality controls (Tables 13 and 15). |
| S9 | Compared with a baseline | A fixed, model-independent baseline is tested on the same examples to label the decline as model-specific or shared. A shared decline remains a valid finding. | **Literature:** informative reference methods (C4), implemented here as a fixed comparator. **Agents:** AlphaMissense deficits were shared by comparison models (Table 13), motivating explicit reporting of model-specific or shared degradation. |

*Table 4 (continued)*

| # | Condition | Requirement | Rationale and supporting sources |
|---|---|---|---|
| S10 | Provenance recorded | Model versions, data, splits, filters, metadata, code, and remaining gaps are recorded. Supporting tables include file size, row count. | **Literature:** version and split provenance (C13). |
| S11 | Artifacts saved | Axis definitions, curve tables, scripts, scores, and a provenance manifest are saved in the run's persistent workspace. | **Literature:** reproducible evaluation definitions and model versions (C13). |

**Supplementary Table 5: Level 1 audit detail: AlphaMissense.** Web-only review. No run performed a new measurement; each used published evidence to identify possible failure conditions. The counts in parentheses show the total number identified and the number listed here.

| Run | Sources used | Potential failure conditions |
|---|---|---|
| GPT r1 | AlphaMissense paper and repository [50]; a CAGI assessment using later data [51]; rare-disease cohort study [52]; ClinGen-focused calibration study [53]; UK Biobank gene-burden evaluation [54]; evaluation of intrinsically disordered regions [55]; functional study of *CPA1* [56]; 2026 ancestry-stratified evaluation [47]. | (1) Variants other than missense variants, which are outside the model's scope by design. (2) Missense variants that cause disease by changing splicing. (3) Intrinsically disordered and low-complexity regions. (4) Proteins with shallow or unreliable sequence alignments. (5) Diseases caused by a mechanism other than loss of the function measured by the model. (6) Gain-of-function, dominant-negative, and neomorphic variants. (7) Patient-level diagnosis, because the model does not receive phenotype, inheritance, or zygosity information. (8) Common disease, pharmacogenomics, and protective variants. (9) Non-human proteins and new protein forms, which cannot be evaluated by adapting the model because its weights were not released. *(15 identified, 9 listed)* |
| GPT r2 | AlphaMissense paper and repository [50]; multiplexed-assay evaluation of 17,696 variants across five genes [57]; 2026 predictor comparison [51]; ClinGen calibration study [53]; rare-disease cohort [52]; *IRF6* case study [58]; *CFTR* benchmark [59]; large study of exonic variants that affect splicing [60]; UK Biobank evaluation [54]; 2026 ancestry-stratified evaluation [47]. | (1) Distinguishing rare benign variants from rare pathogenic variants. Tests that include many common benign variants may make performance look better than it is. (2) Using the default thresholds in clinical care, which may make a "likely pathogenic" label appear stronger than the evidence supports. (3) Real rare-disease candidate lists, where the model may both miss diagnoses and return too many candidates. (4) Disordered regions and short linear motifs. (5) Gain-of-function, hypermorphic, neomorphic, and dominant-negative disease mechanisms. (6) Gene- and domain-specific biology that is hidden by a proteome-wide average. (7) Predicting severity, penetrance, outcomes, or treatment response. (8) Exonic variants that act by changing splicing. *(14 identified, 8 listed)* |
| Opus r1 | AlphaMissense paper, DeepMind release materials, and repository [50]; ClinGen and ACMG calibration studies [53, 61]; Tordai et al. on transmembrane and lipid-interacting regions [62]; Fawzy and Marsh study of 33 variant-effect predictors [63]; *PIEZO1* and *KCNQ4* preprints [57, 64]; *CPA1* study [56]. The run graded every claim: Strong for peer-reviewed, independent, directly relevant evidence; Moderate for preprints or small samples; and Weak for conference abstracts or secondary reporting. | (1) Orphan, fast-evolving, and newly evolved genes. (2) Intrinsically disordered regions. (3) Regions with low structural confidence or interactions with lipids. (4) Gain-of-function and dominant-negative variants. The evidence was graded Moderate because both sources were preprints. (5) Genes for which loss of function is harmless, which can reverse the correct interpretation. (6) Underrepresented ancestry groups. The evidence was graded Moderate and described as debated. *(14 identified, 6 listed)* |

*Supplementary Table 5 (continued)*

| Run | Sources used | Potential failure conditions |
|---|---|---|
| Opus r2 | Studies of *CPA1* and *IRF6* [56, 58]; *PIEZO1* preprint [64]; study of the *POLE* DNA-damage response [65]; 2025 cancer-driver preprint [66]; 2025 study of structural confidence [55]; Tordai paper [62]; benchmark of 65 predictors on a difficult curated clinical dataset [67]. | (1) Direction of effect, especially for genes in which loss of function is harmless. (2) Common pathogenic variants maintained by balancing selection, variants with late-onset effects, and variants that modify risk. (3) Disordered regions and regions with low structural confidence. (4) Proteins with shallow sequence alignments, where scores may fall back toward gene-level averages. (6) Identifying somatic variants and cancer drivers, because the model was trained for a different purpose. (7) Variants other than missense variants, graded *Definitional* because they are outside the model's definition. (10) Underrepresented ancestry groups, the only condition graded Moderate and marked as contested. *(11 ranked, 7 listed)* |
| Kimi r1 | AlphaMissense paper and DeepMind release post [50]; AlphaFold paper [68]; ACMG and AMP standards [61]; ClinGen calibration paper [53]; gnomAD constraint paper [69]; ProteinGym [70]. All seven references were general or foundational rather than focused on individual cases. | (A) Variants and sequences outside the model's scope, including missense variants that alter splicing. (B) Poor input quality, especially when shallow alignments and low structural confidence occur together. (C) Cases in which the model's weak training labels are wrong, including recessive carrier variants, founder variants, late-onset variants, and balanced polymorphisms. (D) Mechanisms the model does not represent well, including activating variants, molecular interfaces, protein dynamics, modification sites, dosage, and tissue-specific effects. (E) Misuse of calibration or decision thresholds, such as interpreting a cohort-level threshold as the probability that an individual case is pathogenic. (F) Evaluations that overstate generalization by using variant frequency to define benign examples. (G) Clinical or domain misuse, including standalone diagnosis, complex disease, pharmacogenomics, and somatic variant interpretation. *(7 groups, all listed)* |
| Kimi r2 | AlphaMissense paper and DeepMind release notes [50]; Europe PMC literature index. The run referred to community calibration research generally rather than citing individual studies. | (1) Late-onset disease, where weak selection against harmful variants can make them appear more benign. (2) Environment-dependent and balanced polymorphisms, such as G6PD deficiency and the sickle-cell variant. (3) Ignoring the underlying rate of pathogenic variants when interpreting the model's released score categories as clinical probabilities. (4) Use of a single-protein structural representation, which misses molecular interfaces and assembly-level mechanisms. (5) Long proteins, which are processed in truncated or overlapping segments. (6) Comparing thresholds across genes, which is less reliable than ranking variants within the same gene. (7) Evaluation circularity as AlphaMissense scores are added to resources that are later used to assess the model. *(19 identified, 7 listed)* |

**Supplementary Table 6: Level 1 audit detail: CONCH.** Web-only review. No run performed a new measurement; each used published evidence to identify possible failure conditions. The counts in parentheses show the total number identified and the number listed here.

| Run | Sources used | Potential failure conditions |
|---|---|---|
| GPT r1 | CONCH paper and model card [71]; benchmark of nineteen pathology foundation models [72]; PathoROB [73]; study of demographic bias in computational pathology [74]. | (1) Open-set diagnosis, when none of the available labels is correct. (2) Diagnosis that requires non-image information, such as age, biopsy site, medical history, or molecular results. (3) Report or caption generation, which the released checkpoint does not support. (4) Rare and pediatric cases, and very specific subtypes. (5) Cellular and subcellular tasks, such as detecting mitoses or microorganisms, counting cells, and grading nuclei. (6) Whole-slide tasks that depend on spatial relationships, such as staging or measuring disease extent. (7) Low-quality or shifted slides, including blur, folds, cautery damage, stain variation, scanner changes, and compression. (8) Demographic subgroups, which have not been measured specifically for CONCH. *(12 identified, 8 listed)* |
| GPT r2 | CONCH paper, supplementary results, and repository [71]; benchmark of nineteen pathology models [72]; prostate-grading study under distribution shift [75]; five-scanner study [36]; adaptation study [76]; study of prompts in digestive pathology [77]; negation benchmark [78]. The run's command-line environment failed before it could run a command, so it relied on web sources only. | (1) Use at a new hospital without any local data. (2) New scanners, staining protocols, fixation methods, or tissue-preparation methods. (3) Underrepresented demographic groups. (4) Detailed or subjective zero-shot diagnosis. CONCH may outperform other models while still performing poorly in absolute terms. (5) Prompts that differ from the carefully designed wording used in benchmarks. (6) Negation, statements of absence, and language that combines several concepts. (7) Non-cancer, pediatric, infectious, transplant, and other non-neoplastic pathology. *(12 identified, 7 listed)* |
| Opus r1 | CONCH paper, preprint, model card, and repository [71]; batch-effect comparison [42]; medical-center robustness index [79]; PathoROB [73]; representational-similarity comparison [80]; scanner-induced domain-shift study [36]; ScanGen [81]; study of stain and scanner variation in sarcomas [82]; feature-extractor benchmark [72]; prostate-grading study [75]; follow-up studies of MI-Zero prompt sensitivity [83]. | (1) Use at a new institution, especially when the model learns a hospital-specific signature instead of the relevant tissue features. (2) Images from a new scanner. (3) Changes in staining or tissue preparation. (4) Mitosis detection, cell counting, and cell segmentation, which the authors place outside the model's intended scope. (5) Low-prevalence findings, small abnormal areas, and needle biopsies, where top-K pooling may ignore a small true-positive region. (6) Magnifications outside the intended range and tasks that require several magnification levels. (7) Calibration, changes in diagnostic categories, and regulatory use. These issues affect many deployment settings. *(14 identified, 7 listed)* |

*Supplementary Table 6 (continued)*

| Run | Sources used | Potential failure conditions |
|---|---|---|
| Opus r2 | CONCH paper, preprint, repository, and model card [71]; CONCHv1.5 and TITAN [84]; PathoROB [73]; medical-center robustness index [79]; batch-effect study [42]; ScanGen [81]; scanner-induced shift study [36]; benchmark of nineteen models [72]; PathBench [85]; segmentation and microsatellite-instability benchmarks [86, 87]; demographic-bias study [74]; FLEX [88]; FAIR-Path [89]; PMC-VQA, BiomedCLIP, and Open-PMC-18M [90–92]; prompt-design study [77]; cross-modality IHC study [93]. | (1) Changes in hospital, scanner, or stain, especially when a technical feature is correlated with the diagnosis. (2) Artifacts and misleading cues, such as pen marks and tissue folds. (3) Messy clinical slides, which may differ from the clean, representative figures common in publications. (4) Changes in zero-shot prompts, class names, or language, including use in languages other than English. (5) Very low or very high magnification, and tasks that require several scales. (6) Flawed evaluation methods, which could make performance under the other twelve conditions look better than it really is. *(13 identified, 6 listed)* |
| Kimi r1 | CONCH paper and official model card [71]; follow-up benchmark of pathology foundation models as slide-level feature extractors [94]; research on failure modes in CLIP- and CoCa-family models [95, 96]. | (1) Modalities other than standard histology, including cytology, frozen sections, immunofluorescence, radiology, gross photography, and non-human samples. (2) Reasoning over gigapixel whole-slide images and detecting very small lesions. (3) Counting, measurement, and grading based on fixed thresholds. (4) Rare, long-tail, and newly renamed conditions, for which the model may give an answer instead of acknowledging uncertainty. (5) Overstated benchmark results. This was the run's lowest-confidence condition. (6) Practical limits: licensing restrictions, no released decoder, no uncertainty estimate, and no clinical validation. *(10 identified, 6 listed)* |
| Kimi r2 | CONCH paper [71]; GitHub repository and benchmark tables; Hugging Face model card; evaluation of histopathology foundation models on multi-stain IHC and immunology tasks [93]; negation benchmark [78]. | (1) Multi-stain immunohistochemistry for autoimmune disease, where CONCH performed no better than ImageNet features. (2) Choosing among many closely related subtypes. (3) Counting Ki-67, HER2, PD-L1, or mitoses. (4) Assessing stage, margins, Gleason grade, or spatial extent. (5) Questions involving negation or diagnostic exclusion. (6) Underrepresented populations. (7) Using fixed thresholds for zero-shot scores, because those scores are not calibrated across sites. *(20 identified, 7 listed)* |

**Supplementary Table 7: Level 1 audit detail: ESMFold2.** Web-only review. No run performed a new measurement. Because no independent evaluation of the released weights existed at the time, the runs calculated failure rates from the developers' reported success rates and used known limitations of related model classes. Kimi r2 evaluated the 2022 predecessor instead of the released model. The counts in parentheses show the total number identified and the number listed here.

| Run | Sources used | Potential failure conditions |
|---|---|---|
| GPT r1 | ESMFold2 preprint, methods, and model card [97]; Biohub limitations page and experimental summary [97]; FoldBench [98]; study of alternative conformations [99]; assessment of mutation effects [100]. | (1) Intrinsically disordered proteins and regions. (2) Metamorphic, allosteric, autoinhibited, and condition-dependent proteins. (3) Short peptides and long multidomain, fibrous, coiled-coil, or repeat proteins. (4) Orphan, taxonomically underrepresented, viral, and highly novel designed sequences. (5) Novel ligands, allosteric pockets, metals, ions, glycans, cofactors, and modifications. (6) Proteins containing D-amino acids or other unusual stereochemistry. (7) Weak, temporary, or nonexistent interactions, where the model may generate a plausible interface for two molecules that do not meaningfully interact. *(11 identified, 7 listed)* |
| GPT r2 | Preprint methods and architecture, and official model card [97]; Biohub limitations and model pages [97]; FoldBench [98]. The run's command-line environment could not start because its temporary storage was full, so it relied on web sources only. | (1) Disordered, flexible, and multi-state proteins. (2) Structures that depend on conditions such as pH, salt, membrane environment, or redox state, because the model does not receive those conditions as inputs. (3) Very large proteins and assemblies that exceed the 384-, 640-, and 768-token contexts used during training. (4) Antibodies, temporary interactions, and flexible interfaces. (5) Unseen ligands and unusual binding chemistry. (6) Noncanonical residues and extensive modifications, which the language model represents with a single unknown token. (7) Low inference budgets, where limited sampling may miss possible structures that confidence ranking cannot recover. *(11 identified, 7 listed)* |
| Opus r1 | Biohub release announcement, model page, technical overview, and preprint [97]; model card; secondary technical coverage; original ESMFold paper [101]; PNAS study of what protein language models learn [102]; research on AlphaFold3 hallucination in disordered regions [103]; study of memorization in fold-switching proteins [21]; stability-prediction research [104]; antibody-docking assessments [105]; FoldBench [98]; CASP16 assessments of nucleic acids and AlphaFold3 [106]; research on de novo binder design [46]. | (1) Fold-switching and metamorphic proteins, because one sequence may correspond to more than one structure. (2) Predicting ordered structures for regions that are actually disordered. (3) Point mutations and variant effects, where predicted coordinates for the mutant and wild type may be almost identical. (4) Orphan sequences and sequences with few evolutionary relatives. (5) Errors inherited from training on AlphaFold-DB predictions, including the original predictor's systematic mistakes and overconfidence. (6) Designing sequences by optimizing them against the model's own score. (7) Poorly calibrated confidence on out-of-distribution inputs. The run treated this as a broad failure that can affect many other conditions. *(19 identified across 11 groups, 7 listed)* |

*Supplementary Table 7 (continued)*

| Run | Sources used | Potential failure conditions |
|---|---|---|
| Opus r2 | Release announcement, technical overview, model card, and bioRxiv preprint [97]; news coverage that included comments from independent experts; adversarial-mutation study of AlphaFold and ESMFold [38]; sequence-based screening research [107]; PNAS studies of what protein language models learn and of single-sequence folding for orphan proteins [102, 108]; original ESMFold paper [101]; de novo design validation study [46]; research on transmembrane structure prediction [109]; ESM Metagenomic Atlas materials [101]. The preprint sites returned HTTP 403 errors, so the run used their abstracts and secondary reporting instead of the full texts. | (1) Orphan and low-homology proteins, including poorly characterized metagenomic sequences. The run ranked this condition first. (2) Novel folds and de novo designs, including confident predictions for sequences that may not fold at all. (3) Point mutations, variant effects, and stability. (4) Conformational dynamics, multiple states, and allostery. (5) Poorly calibrated confidence under distribution shift, especially because the model uses its own confidence scores to choose among sampled structures. (6) Protein design beyond the five targets that received experimental validation. (7) Recently discovered structures and possible benchmark leakage around the September 2021 training cutoff. *(12 identified, 7 listed)* |
| Kimi r1 | Biohub model page and model card [97]; ESM GitHub repository; Hugging Face model card; preprint abstract [97]. The run did not review the full preprint in detail. | (1) Conformational change, allostery, and dynamics. The run considered this a fundamental limitation rather than one that tuning could solve. (2) Mutation, stability, and free-energy questions, also considered fundamental. (3) Incorrect molecular partners or stoichiometry, for which the model may give a confident answer instead of refusing. (4) Ligands with novel chemistry, which are absent from AlphaFold-DB. (5) Interfaces caused by crystal packing rather than real biological interactions. (6) Antibodies outside the benchmark distribution, where similar conserved frameworks in the training data may make results look more general than they are. (7) Reduced inference budgets, which can cause two users of the same model to receive different predictions. *(14 identified across 10 groups, 7 listed)* |
| Kimi r2 | *Evaluated the 2022 predecessor.* This run stated that no released model called ESMFold2 existed and interpreted the name as the second generation of ESMFold. Sources: original ESMFold paper and preprint [101]; related study of generalization beyond natural proteins [110]; public repository; CASP14 and CASP15 assessments [111]. | (1) Orphan sequences and sequences from small protein families, identified by high model perplexity. (2) Highly out-of-distribution designed and synthetic sequences, for which the model may still produce outputs that look well structured. (3) Multi-chain complexes, without an internal warning when the prediction is unreliable. (4) Inputs that are not simply amino-acid chains, including ligands, cofactors, metals, nucleic acids, and glycans. (5) Structural effects of point mutations, where the model may incorrectly predict no change. (6) Circular evaluation against the model used to create its distilled training data. *(13 identified, 6 listed)* |

**Supplementary Table 8: Level 1 audit detail: STATE.** Web-only review. No run performed a new measurement; each used published evidence to identify possible failure conditions. All six runs noted that the main cross-context benchmark gives the model 30% of the perturbations from every test context. The task is therefore few-shot rather than fully zero-shot. The counts in parentheses show the total number identified and the number listed here.

| Run | Sources used | Potential failure conditions |
|---|---|---|
| GPT r1 | STATE evaluation paper [3]; *Mechanisms Matter*, which tests STATE directly using cross-context splits [112]; *Virtual Cells Need Context, Not Just Scale* [113]; Systema perturbation scoring research [114]; 2025 predictor comparison [115]; 2025 evaluation [32]; real-world benchmark preprint [116]; Tahoe-100M dataset description [7]. | (1) A completely unseen gene, drug, or cytokine, which the model cannot handle without retraining. (2) A new drug structure or a dose between the doses seen during training. (3) A new biological context with no matched control cells. (4) A context driven by a different regulatory mechanism, especially when the baseline RNA profile looks similar and therefore hides the difference from the model. (5) Weak perturbations, partial edits, and inconsistent editing efficiency. (6) Toxic perturbations or perturbations that change cell proliferation, where cell survival is itself the important outcome. (7) Rare response patterns and random branching into different cell states. (8) Global changes in RNA abundance that compositional normalization removes. (9) High-stakes use on unfamiliar data, because the model has neither calibrated uncertainty nor an option to reject unreliable predictions. *(16 identified, 9 listed)* |
| GPT r2 | STATE paper [3]; real-world benchmark preprint [116]; study of causal transport across contexts [113]; analysis of the perturbation discrimination score [117]; research on replication in single-cell data [118]; Tahoe-100M dataset card and official description [7]. | (1) Unseen perturbations, which the default architecture cannot represent. (2) Truly zero-shot contexts with different biological mechanisms. (3) A new perturbation in a new context at the same time, which is a common setting in drug discovery. (4) Changes in dose, time, or treatment schedule. (5) Weak, noisy, or inefficient perturbations. (6) Rare responding cells and perturbations that cause cells to branch into different states. (7) Counterfactual predictions for individual cells and claims about causal mechanisms. (8) Changes in laboratory or experimental platform. *(10 identified, 8 listed)* |
| Opus r1 | STATE preprint and code repository [3]; Arc posts and Virtual Cell Challenge summary [119]; about eight independent benchmark and critique papers published from 2024 to 2026, including PertEval-scFM, PerturBench, and VCBench [30, 115, 120–122]. | (1) New perturbations. Because the standard model represents perturbations as one-hot categories, it cannot accept a category it has never seen. (2) Predicting genetic perturbations across cell lines, where performance is at or near a mean baseline. (3) Entirely new datasets, platforms, or laboratories, which the authors said they did not test. (4) Primary tissue, living organisms, disease states, and developmental contexts. (5) Weak, unusual, and extremely strong perturbations, for which predictions may collapse toward the population average. (6) Outcomes other than RNA expression. (7) Dose response and changes over time. (10) Improvements in reported metrics that do not produce useful real-world predictions. *(10 ranked, 8 listed)* |
| Opus r2 | STATE preprint, repository, and model cards [3]; Tahoe, Parse, and Replogle–Nadig dataset papers [4–7]; independent benchmark and critique research. The run labeled each condition as *authors admit*, *independent*, or *architectural*. | (1) A new drug or gene target, which one-hot encoding cannot represent. (2) Genetic perturbation prediction, where the paper's own results are about equal to a baseline. (3) Dose response and changes over time, which are not model inputs. (4) Non-coding RNA and genes with low expression, which are excluded from the model's gene set. (5) Applying a model trained mainly on cancer cell lines to normal tissue or toxicology. (6) Failure of the assumption that noise is independent, because real dropout rates are related to expression level. (7) Claims about biological mechanisms or causal effects beyond the observed data. (8) Bias from training only on genetic perturbations that passed an efficacy filter. *(14 identified, 8 listed)* |

*Supplementary Table 8 (continued)*

| Run | Sources used | Potential failure conditions |
|---|---|---|
| Kimi r1 | STATE preprint and repository [3]; Tahoe-100M dataset paper [7]; Replogle and Nadig dataset papers [4, 5]; Parse cytokine dataset [6]; independent baseline research, including PerturbBench and the Virtual Cell Challenge [30, 115, 119, 121]. | (1) A new perturbation in a new context. The evaluation tested these two forms of novelty separately, not together. (2) Genes outside the 2,000 highly variable genes used by the model, whose effects are invisible by design. (3) Global changes in RNA composition that normalization removes. (4) Perturbations that change survival or population composition, because set matching only models the cells that remain. (5) Theoretical guarantees based on unlimited data when the real dataset is finite. (6) Evaluation with metrics introduced by the model's authors, especially when their own model performs best under those metrics. (7) Unclear pretraining overlap between the embedding model and the transition model. (8) Predictions pulled toward the context mean, consistent with the paper's own description of matching the mean baseline. *(10 conditions across 5 groups, 8 listed)* |
| Kimi r2 | STATE preprint and repository [3]; Tahoe, Replogle, Nadig, and Parse dataset descriptions [4–7]; Virtual Cell Challenge [119]; independent baseline research cited in the preprint. The run graded severity according to the intended use rather than treating it as a fixed property of each condition. | (A1) Weak perturbations that are similar in size to normal cell-to-cell variation, making their effects difficult to detect statistically. (B4) Patient and clinical samples, especially when the model is used to support clinical claims. (C1) A new perturbation identity, which remains a central unsolved problem in the field. (C3) Dose and time, which the model does not represent. (C4) Unseen types of perturbation, including gene activation, base editing, prime editing, degraders, biologics, and physical stress. (C5) Lethal and strongly selective perturbations, where sequencing observes only the surviving cells. (E3) Confounded experiments, where the model may mistake a batch effect for a response to the perturbation. (E4) Predicting effects larger or smaller than those observed during training. *(20 identified across 5 groups, 8 listed)* |

**Supplementary Table 9: Level 2 audit detail: AlphaMissense.** Empirical audit. The protocol did not require novelty screening or a model-independent baseline. Because the AlphaMissense weights are not public, every run reanalyzed the published score tables instead of running the model directly.

| Run | Data used | Conditions tested | Main result |
|---|---|---|---|
| GPT r1 | AlphaMissense Supplementary Data S8 [50]: 358,018 single substitutions from 87 assays and 72 proteins; ProteinGym v1.3 [70] as the external dataset; AlphaFold residue-level pLDDT [68]. | Alignment depth ($\rho$ = 0.109, $p$ = 0.318 across 86 assays), which did not pass the initial screen. Structural disorder measured by pLDDT, tested with binary labels at every cutoff from 50 to 90 and with four pLDDT bands. | **pLDDT below 70.** In the discovery data, 57 paired assays across 49 proteins scored 0.297 below the cutoff and 0.535 above it, a difference of 0.238 (CI 0.170–0.308). After removing every discovery protein, the external dataset contained 33 paired assays across 29 proteins. Scores were 0.087 below the cutoff and 0.435 above it, a difference of 0.348 (CI 0.269–0.425; paired Wilcoxon $p = 5.22 \times 10^{-8}$). The result was positive at every cutoff tested, although the four-band curve was not strictly monotonic below 70 in the external dataset. *Replicated.* |
| GPT r2 | ProteinGym [70] for both discovery and replication; ClinVar [123] for comparison with clinical labels; released AlphaMissense score tables [50]. The run published three raw input files with SHA-256 checksums. | Grantham severity [124], including a rank-independent test, clinical labels, and twelve combinations of cutoffs. Genetic accessibility (difference −0.020) and location near a sequence end (difference −0.087, opposite the predicted direction) did not pass the prespecified screen. | **Grantham substitution severity**, comparing conservative substitutions ($\leq$ 50) with radical substitutions (> 100). The discovery dataset contained 32 assays across 26 proteins and 126,545 matched substitutions. Scores were 0.352 for conservative and 0.444 for radical substitutions, a difference of 0.092 (CI 0.043–0.143). The difference was positive in 28 of 32 assays (sign-flip $p$ = 0.0012). In the fixed replication dataset, which contained 59 assays across 54 entirely different proteins, the difference was 0.017 (CI −0.017–0.051), was positive in 34 of 59 assays, and had $p$ = 0.343. *The result failed the run's prespecified confirmation rule.* |
| Opus r1 | 314,032 deep mutational scanning (DMS) variants from 92 assays and 75 proteins; 208,898 ClinVar missense variants across 15,132 proteins, including 67,799 pathogenic and 141,099 benign variants [123]; ESM2 [101] and Grantham distance [124] as comparators on the same variants; metapredict as a disorder measure independent of AlphaFold [125]. | Structural disorder measured by pLDDT, then repeated on different proteins, with different types of outcome, and with a different disorder measure. Alignment depth as a possible confound ($r$ = 0.01, $p$ = 0.91), which did not pass the initial screen. Clinical labels. | **Structural disorder measured by pLDDT.** Absolute Spearman correlation within each assay across four pLDDT bands was 0.192, 0.225, 0.370, and 0.512 for AlphaMissense; 0.128, 0.219, 0.370, and 0.493 for ESM2; and 0.101, 0.111, 0.146, and 0.172 for Grantham. In paired comparisons within proteins, performance was 0.19 in disordered regions and 0.51 in ordered regions (Wilcoxon $p = 2 \times 10^{-8}$), with worse performance in disordered regions for 32 of 34 proteins. Replacing pLDDT with metapredict produced the same decline, from 0.46 to 0.18. ClinVar AUC remained between 0.94 and 0.96 and therefore hid the effect. *Replicated; the same pattern appeared in ESM2, so it is not unique to AlphaMissense.* |

*Supplementary Table 9 (continued)*

| Run | Data used | Conditions tested | Main result |
|---|---|---|---|
| Opus r2 | 198,208 ClinVar variants [123]; 98.1% were successfully matched, and 165,175 variants across 14,439 proteins remained after checking residue numbering. A manual check confirmed the residue identity for all 581 sampled variants. The analysis also used 315,034 DMS variants from 89 assays and 75 proteins, and ran ESM2 150M [101] locally as a comparator. | Structural disorder measured by pLDDT using both functional and clinical labels. Sampling bias as a possible explanation for the lack of an effect in the clinical data. Protein-level versus residue-level analysis ($\rho$ = 0.06 and −0.06), which did not pass the initial screen. Score compression, which was not analyzed further. | **Structural disorder measured by pLDDT.** Across the pLDDT bands, AUC was 0.787, 0.743, 0.624, and 0.591, while absolute Spearman correlation was 0.52, 0.39, 0.14, and 0.15—a reduction of about 3.4-fold. Across the 63 assays that covered both ordered and disordered regions, median performance fell from 0.49 to 0.16 ($p = 2.1 \times 10^{-9}$), and 81% of assays performed worse in disordered regions. ClinVar AUC remained between 0.954 and 0.958 except for a drop to 0.822 at pLDDT 20–30. This drop was explained by the pathogenic base rate: 1.8% below pLDDT 30 versus 52% above pLDDT 90. ESM2 declined in parallel, from 0.50 to 0.08. *Replicated; the same pattern appeared in ESM2.* |
| Kimi r1 | ClinVar variants with at least one review star, 176,833 variants; Humsavar, 71,188 variants [126]; 44,901 Humsavar variants not present in ClinVar; ClinVar variants with at least two review stars, 53,370 variants [123]; Grantham matrix [124], checked against 18 published values with no mismatches. | Whether performance follows an inverted-U relationship with Grantham severity, tested with a parametric model and with score-decile cutoffs defined in ClinVar and applied unchanged to Humsavar. Protein length above 2,912 residues. Position near the N terminus, which failed in Humsavar (difference 0.0008, CI −0.0068–0.0091). | **Non-monotonic relationship with Grantham severity.** AUC was highest for the intermediate band (51–100) in all four datasets: 0.9573, 0.9071, 0.8753, and 0.9543. Differences relative to conservative substitutions were 0.0203, 0.0181, 0.0305, and 0.0268; differences relative to radical substitutions were 0.0119, 0.0274, 0.0161, and 0.0111. Every confidence interval excluded zero, including the interval based on 44,901 variants not present in the discovery source. The protein-length result also replicated. *Replicated.* |
| Kimi r2 | ClinVar, dated 2026-07-21: 209,552 variants across 15,403 genes in a 441 MB file [123]; ProteinGym v1.2 as the multiplexed assay of variant effect (MAVE) dataset: 89 assays covering 319,009 variants and 77 proteins [70]; AlphaMissense hg38 release, a 643 MB file with an MD5 checksum matching the published storage ETag [50]. | Discrimination within each gene, with a predictor based only on the gene's mean score as a control. ClinVar review status, which was not analyzed further. Grantham quartiles with cutoffs fixed at 38, 64, and 99, which failed in MAVE. Residue position, for which the direction of the effect reversed in MAVE. | **Discrimination within genes.** AlphaMissense achieved a pooled AUC of 0.873 across all genes, but the median AUC within individual genes was 0.631 (IQR 0.551–0.742), and 82% of 326 genes had an AUC below 0.80. A control predictor that used only each gene's mean score already achieved an AUC of 0.847. In the MAVE data, the median AUC within each assay was 0.787, and 87% of assays were below the pooled headline result ($p = 2.7 \times 10^{-14}$). *Replicated.* |

**Supplementary Table 10: Level 2 audit detail: CONCH.** Empirical audit. The protocol did not require novelty screening or a model-independent baseline. All six runs independently tested effective spatial resolution by reducing the resolution of the same images and then resizing them to the model's input size.

| Run | Data used | Conditions tested | Main result |
|---|---|---|---|
| GPT r1 | Licensed CONCH checkpoint [71], SHA-256 `40a9644b...b0e9ae7c`, with a fixed repository revision; CRC-VAL-HE-7K [127, 128] as the main dataset, containing 900 images across 9 classes; Stanford STARC-9 as the replication dataset, containing 900 images across 6 overlapping classes [129]. | Effective spatial resolutions of 448, 224, 112, and 56 pixels, with every image resized to 448 pixels before scoring; the size and location of the decline between adjacent resolution levels; the candidate label set; macro AUC as a threshold-independent outcome. | **Effective spatial resolution.** Accuracy in the main dataset was 75.8%, 74.9%, 68.3%, and 33.8%, with a slope of −13.26 percentage points per halving of resolution (CI −14.33 to −12.18) and an odds ratio of 0.433. Accuracy in the Stanford dataset was 78.0%, 78.7%, 80.1%, and 67.3%, with a slope of −3.06 (CI −4.05 to −2.06) and an odds ratio of 0.818. McNemar $p$-values were $3.8 \times 10^{-105}$ and $9.4 \times 10^{-12}$. Macro AUC fell much less: from 0.971 to 0.904 and from 0.976 to 0.925. Restricting the analysis to the six shared classes reduced the apparent nine-class loss from 42.0 to 11.0 points; this analysis was labeled exploratory. *Replicated; no comparator model was tested.* |
| GPT r2 | The same licensed checkpoint [71]; CRC-VAL-HE-7K and Stanford STARC-9, using 450 images from each dataset [127–129]; random seeds 20260716 and 20260717. | The same four resolution levels; the size and location of the decline between adjacent levels; the shared six-class label set; macro AUC. | **Effective spatial resolution.** Accuracy in the main dataset was 76.0%, 76.4%, 69.3%, and 36.9%, with a slope of −12.44 (CI −14.00 to −10.89) and a total change of −39.11 points. Stanford accuracy was 82.9%, 82.7%, 82.0%, and 68.7%, with a slope of −4.33 (CI −5.68 to −2.98) and a total change of −14.22 points. McNemar tests gave $p = 2.98 \times 10^{-46}$ (182 versus 6 discordant cases) and $p = 6.15 \times 10^{-11}$ (82 versus 18). Macro AUC fell from 0.971 to 0.912 and from 0.976 to 0.932. *Replicated; no comparator model was tested. The generalized estimating equation for the full main dataset did not converge and was excluded.* |
| Opus r1 | The official weights returned an HTTP 401 error, so the run used an unrestricted, byte-consistent mirror. The file contained 802,235,437 bytes, had MD5 `91a630c2...`, loaded with no unexpected keys, and reproduced CONCH's published confusion pattern [71]. Datasets were CRC-VAL [127, 128], LC25000 [130], and PatchCamelyon [131]. PLIP was acquired and tested as a comparator [132]. | Effective resolution in micrometers per pixel across nine levels and three datasets; JPEG compression as a second condition; PLIP tested with the same resolution levels. | **Effective spatial resolution.** CRC-VAL balanced accuracy was 0.769, 0.733, 0.706, 0.513, 0.400, 0.278, 0.144, 0.081, and 0.074 as resolution changed from 0.50 to 7.00 $\mu$m pixel$^{-1}$. At least 90% of performance was retained through 1.0 $\mu$m pixel$^{-1}$; performance fell to half at 1.63, with a logistic midpoint of 1.70, Hill slope of 2.12, and $R^2 = 0.996$. PatchCamelyon reached half performance at 2.48 ($R^2 = 0.982$), placing its sharp decline in the same 1.6–2.5 range as CRC-VAL. LC25000 was much more tolerant. JPEG compression produced a similar decline, from 0.776 at quality 95 to 0.111 at quality 5. The PLIP and CONCH curves crossed: CONCH was better at 20× magnification (0.769 versus 0.688), but PLIP was better below about 1.4 $\mu$m pixel$^{-1}$ (0.660 versus 0.513). *Replicated; PLIP's result shows that information needed for classification remained in the degraded images, so CONCH's representation caused the additional failure.* |

*Supplementary Table 10 (continued)*

| Run | Data used | Conditions tested | Main result |
|---|---|---|---|
| Opus r2 | MahmoodLab CONCH `pytorch_model.bin`, a 765 MB file from a local cache, with image size fixed at 224 pixels [71]; CRC-VAL and three additional datasets for replication [127, 128]; a PLIP comparison script that was written and partially run [132]. | An initial screen using $n = 450$: resolution, blur, JPEG compression, saturation, hue, and Gaussian noise. Resolution was then tested with $n = 1800$ across twelve levels and four datasets. Changes in the image embedding were measured to investigate the cause. | **Downsampling factor.** Accuracy was 0.786, 0.776, 0.757, 0.747, 0.737, 0.709, 0.652, 0.471, 0.242, and 0.137 as the downsampling factor increased from 1.0 to 10.0. The main decline began at a factor of 3.5, the first significant drop occurred at 2.5, and half the performance loss occurred at 6.3 (about 3.2 $\mu$m pixel$^{-1}$), with steepness 4.1 and $R^2 = 0.998$. Three additional datasets had decline points at 4.5, 4.5, and 6.0 and half-loss points at 6.3, 11.4, and 9.6. Embedding drift increased from 0 to 0.53 and was correlated with accuracy ($r = -0.88$, $p = 2 \times 10^{-4}$), placing the source of the decline in the image encoder. *Replicated; because the PLIP test did not finish, the run could not determine whether this failure was specific to CONCH.* |
| Kimi r1 | Unrestricted CONCH mirror [71], validated by reproducing 78.3% zero-shot accuracy compared with the published 79.1%, and 95.5% linear-probe accuracy compared with the published 93.0%; NCT-CRC-HE-100K as the discovery dataset, with 18,000 patches balanced at 2,000 per class; CRC-VAL-HE-7K as the independent dataset, with 7,180 patches from a different institution, scanner, and patient population [127, 128]. The run used one A100 40 GB GPU. | Downsampling across nine levels in both datasets; Gaussian blur as a matched control; a logistic probe trained on clean features as an internal comparator; conversion toward grayscale. | **Downsampling factor.** Discovery accuracy was 0.835, 0.847, 0.848, 0.843, 0.828, 0.598, 0.316, 0.113, and 0.077. Accuracy in the independent dataset was 0.783, 0.797, 0.804, 0.807, 0.824, 0.615, 0.354, 0.125, and 0.096. Performance fell to half at factors of 6.76 and 7.15, a difference of only 0.08 $\log_2$ units; the curves had a correlation of 0.998. At factor 16, accuracy was 7.7%, below the 11.1% chance rate, because at factor 8 the model predicted background for 12,388 of 18,000 images. A probe trained on clean CONCH features declined more gradually, reaching 65.7% rather than 31.6% at factor 8. Moving toward grayscale caused a linear decline with no sharp break. *Replicated; image–text alignment declined about twice as quickly as the visual representation itself.* |

*Supplementary Table 10 (continued)*

| Run | Data used | Conditions tested | Main result |
|---|---|---|---|
| Kimi r2 | Balanced discovery dataset of 9,000 NCT images, with 1,000 images per class and random seed 0; the full CRC-VAL-HE-7K dataset, which was not used until the protocol was fixed [127, 128]; CONCH [71] under two protocols: official prompts with the standard 448-pixel pipeline, and seven custom prompt templates with images passed directly at 224 pixels. | Downsampling across six levels under both protocols and in both datasets; JPEG quality across seven levels; whether mild downsampling improves performance. | **Downsampling factor.** Under Protocol A, discovery accuracy was 0.801, 0.765, 0.405, 0.141, 0.053, and 0.044, while replication accuracy was 0.793, 0.777, 0.472, 0.157, 0.034, and 0.009. Slopes per doubling were −1.10 (CI −1.12 to −1.07) and −1.19 (CI −1.22 to −1.17), and performance reached half-degradation at factors of 3.6 and 4.2. Under Protocol B, slopes were −0.995 and −0.925, with half-degradation near a factor of 7.8. The pattern of decline appeared in all four dataset–protocol combinations, but preprocessing shifted the half-degradation point by about a factor of two. JPEG quality produced the same pattern, with accuracy falling from 0.801 to 0.111 and from 0.793 to 0.116. The apparent improvement from mild downsampling disappeared when the protocol changed. *Replicated; the decline occurred under both protocols, but its threshold depended on the protocol.* |

**Supplementary Table 11: Level 2 audit detail: ESMFold2.** Empirical audit. The protocol did not require novelty screening or a model-independent baseline. The model name did not identify a single implementation: two runs tested an unversioned public folding service, one ran the released ESMFold2 checkpoint, and three tested the 2022 predecessor. Absolute results should therefore not be compared across rows.

| Run | Data used | Conditions tested | Main result |
|---|---|---|---|
| GPT r1 | Public ESM Atlas fold-on-demand endpoint [101]; RCSB structures first released in 2024 for discovery and in 2025 for replication [133]. Structures were determined by X-ray crystallography at 2.5 Å resolution or better, contained a single chain and entity, and were 40–699 residues long. Targets were ordered by SHA-256. The endpoint returned coordinates for 291 of 391 candidate sequences, from which 200 targets were used. | Length below 200 residues, the boundary specified by the published training mixture, tested separately for accuracy and confidence; the gap between confidence and accuracy; nine sequence-only features explored after the main analysis with Benjamini–Hochberg correction. | **Length below 200 residues.** Accuracy did not differ consistently by length. The 2024 difference was −0.0048 (CI −0.0175–0.0107, $p = 0.915$), while the 2025 difference was 0.0022 (CI −0.0115–0.0200, $p = 0.652$). The result was unchanged after requiring 80% structure coverage and after a Huber adjustment. Confidence did differ: −0.0799 (CI −0.1278 to −0.0506) in 2024 and −0.0526 (CI −0.0781 to −0.0165) in 2025. The confidence–accuracy gap shifted by −0.0677 and −0.0507. None of the additional features passed a false discovery rate of 0.05. *The accuracy result failed. The remaining effect was lower confidence, not lower accuracy: short proteins were just as accurate but were more likely to be rejected by a fixed confidence threshold.* |
| GPT r2 | The same public endpoint, evaluated as it existed on a fixed date [101]; three non-overlapping cohorts of 60 entities from 2022, 2024, and 2025, with 15 targets in each of four length bands from 60 to 500 residues [133]. | Seven sequence-only features screened in the 2022 cohort. The maximum fraction of any single residue type was selected as the measure of sequence composition bias. Tertile cutoffs and a degraded-fold threshold of lDDT below 0.90 were fixed before testing the 2025 cohort. The analysis also tested the feature continuously, tested mean accuracy, and removed closely related sequences. | **Sequence composition bias.** In the most biased versus the most balanced tertile, degraded folds occurred in 3 of 20 versus 2 of 20 targets in 2022, 7 of 21 versus 1 of 20 in 2024, and 3 of 25 versus 1 of 25 in 2025. The pooled difference in the independent cohorts was 17.3 percentage points (CI 4.2–30.4), with a risk ratio of 4.11 and a one-sided permutation $p = 0.015$. The direction remained the same after removing sequences above 30% identity and requiring 90% resolved residues. The continuous analysis reproduced poorly ($\rho = -0.045$, $p = 0.366$), and the prespecified cohort alone was not significant. Median lDDT remained between 0.961 and 0.967, so composition bias changed the risk of rare bad predictions rather than average performance. *Replicated as a threshold comparison; the run graded the result reproducible but moderate.* |

*Supplementary Table 11 (continued)*

| Run | Data used | Conditions tested | Main result |
|---|---|---|---|
| Opus r1 | Original ESMFold [101]. The run explicitly noted that no model had been officially released under the name ESMFold2. It used two separate, nonredundant PDB samples: 218 chains first released in 2022 and 219 first released in 2023 [133]. Structures were determined by X-ray crystallography at 2.5 Å resolution or better and were 40–500 residues long. Each sample was reduced to one representative per 30%-identity cluster, with no sequences shared between samples. | Masked pseudo-perplexity from the language-model component, calculated over 48 sampled positions per protein; length as a possible confound; predicted confidence as a possible explanation of the effect; a confidence threshold as a practical filter. | **Sequence pseudo-perplexity.** Its Spearman correlation with TM-score [134] was −0.275 (CI −0.41 to −0.14, $p = 3.8 \times 10^{-5}$) in the 2022 sample and −0.396 (CI −0.51 to −0.27, $p = 1.2 \times 10^{-9}$) in the 2023 sample. Mean TM-score in the top quartile was 0.929 and 0.935, compared with 0.824 and 0.819 in the bottom quartile. The decline was concentrated in the fourth quartile, which contained 87% of all failures. Length had no detectable effect ($p = 0.84$ and 0.47). Predicted confidence fully explained the relationship: the adjusted effect fell from −0.065 to 0.002 and from −0.000 to zero. Rejecting predictions with confidence below 0.7 retained 93% of predictions and reduced the wrong-fold rate from 3.4% to 0.5%. *Replicated; the model's confidence score identified the problem rather than hiding it.* |
| Opus r2 | The only run that evaluated the released ESMFold2 checkpoint [97]. It ran the 6B backbone, diffusion head, three loops, and confidence head locally on one A100 40 GB GPU, using two random seeds per target and 20 sampling steps. Homolog counts came from a public MMseqs2 service [135]. The discovery sample used 2022 structures, and the replication sample used structures from 2024–2025. Both were drawn after global clustering at 30% sequence identity [133]. | Evolutionary depth in three homolog-count groups; later release years for replication; length and disorder as possible confounds; an intervention that varied the number of alignment sequences from 1 to 1,024 for the same target; predicted confidence as a filter. | **Evolutionary depth.** For targets with fewer than 100 homologs, mean C$\alpha$-lDDT [136] was 0.50 and the catastrophic-failure rate was 73%. For 100–1,000 homologs, mean lDDT was 0.46 and the failure rate was 75%. At 1,000 or more homologs, mean lDDT was 0.94 and no catastrophic failures occurred. Spearman correlation was 0.69 ($p = 5.4 \times 10^{-10}$), with a sharp threshold near 1,000 homologs. The result replicated in the later samples at $\rho = 0.73$ and 0.65, with failure rates of 83% and 60% falling to zero. Length had no detectable relationship ($\rho = -0.01$). Confidence identified every failure with about 1% false positives. *Replicated. Increasing alignment depth during inference did not rescue the same targets (mean lDDT change 0.007). This suggests that the limitation lies in what the language model learned about the protein family, not in the amount of alignment data supplied during one prediction.* |

*Supplementary Table 11 (continued)*

| Run | Data used | Conditions tested | Main result |
|---|---|---|---|
| Kimi r1 | ESMFold v1 weights and the 3B language model from the original release source [101], used with the original inference library. The evaluation included 524 chains in two non-overlapping and mutually nonredundant samples: 266 structures released from mid-2022 to mid-2023 and 258 released in 2024. Any chain in a discovery cluster at 30% identity was excluded from the later sample. Each sample included 45 chains in each of six length bands from 40 to 400 residues [133]. | Chain length below 80 residues; a logistic failure model fit in the discovery sample and tested out of sample; consistency of the effect across cohorts; resolved coverage and crystal context as possible confounds; whether the length normalization in TM-score created the effect. | **Chains shorter than 80 residues.** In discovery, mean TM-score was 0.774 and the failure rate was 14.0%, compared with 0.927 and 0% for chains of at least 300 residues. In replication, the corresponding values were 0.794 and 14.3% versus 0.921 and 0%. Spearman correlation with length was 0.318 ($p = 1.2 \times 10^{-7}$) and 0.330 ($p = 5.6 \times 10^{-8}$), with a pooled value of 0.323. A logistic model fit on the discovery data predicted a 12.6% failure rate at 60 residues, compared with 14.3% observed. Coverage did not explain the result. Two checks ruled out TM-score normalization: predicted confidence showed the same gradient, and the failures formed a separate group at TM-scores of 0.24–0.30. *Replicated; the result was linked to the published rejection-sampling procedure, which downweighted chains shorter than 256 residues.* |
| Kimi r2 | ESMFold 3B weights [101]; CASP14 as discovery, with 54 targets and a median length of 197 residues, and CASP15 as replication, with 72 targets and a median length of 292 [111]; reference database built from 1,142,466 PDB chain sequences, reduced to 556,659 sequences released before the 2020-05-01 cutoff [133]; MMseqs2 for sequence search [135]. | Maximum sequence identity to any structure released before the cutoff, tested continuously and with a parametric model; a model fit on CASP14 and tested on CASP15; four definitions of identity and coverage; predicted confidence as a possible filter. | **Identity to structures released before the cutoff.** In discovery, $\rho = 0.282$ ($p = 0.039$), and mean TM-score was 0.652 without a homolog versus 0.760 with a homolog ($p = 0.051$). In replication, $\rho = 0.422$ ($p = 2.2 \times 10^{-4}$), and mean TM-score was 0.668 without a homolog versus 0.822 with one ($p = 0.0074$). The fraction reaching TM-score 0.5 rose from 0.66 to between 0.92 and 1.00. In the pooled data, $\rho = 0.390$ ($p = 6.5 \times 10^{-6}$), and the slope was 0.025 TM-score per ten percentage points of identity ($p = 0.002$). The out-of-sample correlation was 0.347 ($p = 0.003$). Rank correlation remained between 0.368 and 0.414 across the four definitions. *Replicated. Confidence tracked accuracy at 0.846 and 0.553 but failed on specific named examples, so it was not a reliable filter.* |

**Supplementary Table 12: Level 2 audit detail: STATE.** Empirical audit. The protocol did not require novelty screening or a model-independent baseline. All six runs tested perturbation effect size and found a similar relationship between effect size and measured performance. They differed in whether they accounted for the lower signal-to-noise ratio of weak perturbations.

| Run | Data used | Conditions tested | Main result |
|---|---|---|---|
| GPT r1 | Released STATE evaluation outputs [3] from three model directories: chemical perturbations at three doses in five held-out cell lines [7]; CRISPR interference in four cell lines [4, 5]; and cytokine perturbations in five immune cell types [6]. The analysis covered 11,893 perturbation–context pairs. Every repository revision was fixed, and every input had a recorded SHA-256 checksum. | Observed response breadth, measured as the number of genes differentially expressed at a 5% false discovery rate; the same condition under a cosine-based discrimination metric; consistency across held-out contexts; administered dose as a predictor chosen before observing the response; the same comparison under absolute-error metrics. | **Observed response breadth.** Change in delta-Pearson per within-context standard deviation was 0.0223 (CI 0.0118–0.0329) for chemical perturbations, 0.1837 (CI 0.1791–0.1884) for CRISPR, and 0.0488 (CI 0.0335–0.0640) for cytokines. The gaps between the lowest and highest quintiles were 0.0487, 0.5150, and 0.1466. The cosine metric reproduced the pattern at 0.0350, 0.1783, and 0.0225. The relationship appeared in all four CRISPR contexts (0.711–0.816) and all five cytokine contexts (0.075–0.539), but not throughout the chemical dataset, where two of five cell lines had small effects in the opposite direction. A tenfold increase in dose produced 29.6 more truly differentially expressed genes (CI 20.0–39.2). Under absolute-error metrics, the direction reversed: one standard deviation improved the cosine score by 0.178 but worsened L1 and L2 discrimination by 0.112 and 0.121. *Replicated, but not consistently across all contexts. Because no matched baseline was analyzed by effect size, the result shows where STATE's performance changes but not whether the pattern is unique to STATE.* |
| GPT r2 | The same released evaluation outputs [3], including both the gene-space and embedding-space versions of the model. | The lowest within-context quartile of the same differential-expression count, using thresholds of 231–422 genes in the chemical dataset and 20–83.5 genes in the CRISPR dataset; rate of predicting the correct direction; change in mean squared error; the embedding-space model; performance across all four quartiles; dose as a basic validity check. | **Weak-response quartile.** In the chemical dataset, performance was 0.482 in the weak quartile and 0.555 in the other quartiles, a gap of 0.074 (CI 0.057–0.090). In the CRISPR dataset, the corresponding values were 0.050 and 0.354, a gap of 0.304 (CI 0.292–0.315). The gap was positive in all nine held-out contexts. Correct-direction gaps were 0.044 and 0.174. The embedding-space model showed gaps of 0.081 and 0.209, but the run treated this as a sensitivity check rather than an independent replication because both models used the same ground-truth conditions. The shape across all four quartiles did not replicate: the chemical dataset peaked in the third quartile (0.481, 0.553, 0.590, 0.523), while the CRISPR result increased monotonically. *Underperformance for weak responses replicated, but the full quartile pattern weakened. The run limited its claim because correlations between two nearly zero vectors are inherently noisy, making model error hard to separate from low signal-to-noise. It also noted that the hypothesis was chosen after reviewing the available evaluation fields rather than specified in advance.* |

*Supplementary Table 12 (continued)*

| Run | Data used | Conditions tested | Main result |
|---|---|---|---|
| Opus r1 | Released checkpoints, held-out predictions, ground-truth files, and per-perturbation metric tables for the four-cell-line CRISPR dataset [4, 5] and the cytokine dataset [6] [3]. The main metric was reimplemented and matched the released table with a correlation of 0.99999999. Rerunning the released checkpoint reproduced the packaged predictions at a correlation of 0.986. | Perturbation effect size; cell count as a possible confound; two simple baselines exposed to the same measurement noise—predicting the average perturbation and predicting no change; an independent cytokine dataset; transfer to an unseen cell type. | **Perturbation effect size.** Spearman correlation between the number of differentially expressed genes and performance per perturbation was 0.86 across 945 held-out perturbations, 0.82 pooled across four cell lines, and 0.83 after adjusting for cells per perturbation. The result replicated in the cytokine dataset, which used a different laboratory, perturbation type, and cell system, at 0.41 and 0.35. *The relationship replicated, but the baselines changed its interpretation. Mean advantage over predicting the average perturbation was 0.005, 0.062, 0.083, and* −0.039 *in the four cell lines. In the weakest quartile, STATE outperformed predicting no change for 0 of 242 perturbations. When transferring to a held-out cell type, its advantage was negative in every effect-size group in all four cell lines, and discrimination moved toward chance as effect size increased (*−0.53, −0.68*). STATE therefore added the least in the transfer setting emphasized as a main use case.* |
| Opus r2 | The same released files across four cell lines and two generalization settings, covering 9,817 held-out perturbations, plus the cytokine dataset [3–6]; released inference command run on a CPU. The main metric was reimplemented and matched the released table exactly, with mean absolute difference 0.0000. | Perturbation strength in all eight released conditions; a split-half noise ceiling for each perturbation, with performance reported as a fraction of the maximum any predictor could achieve; the average-perturbation baseline; an independent cytokine dataset; cells per perturbation; predicted direction and magnitude as possible explanations. | **Perturbation strength.** Spearman correlation was 0.83, 0.80, 0.81, and 0.86 within context and 0.81, 0.71, 0.72, and 0.82 in transfer. The cytokine dataset reproduced the result at 0.83 and 0.87, for a total of ten independent held-out evaluations. *The difference in measurement noise was real but did not explain the relationship. The noise ceiling rose from 0.32 to 0.95 across quintiles, but the fraction of achievable signal captured by the model still fell from 0.79 to 0.10. The strength term remained after joint regression on ceiling-normalized performance ($t$ =* 50.2*). Advantage over the average-perturbation baseline was* −0.019, −0.005*, 0.068, 0.158, and 0.211 across quintiles. The win rate was 0.43 in the weakest quintile. Cosine similarity with the true expression shift was 0.04 in the weakest quintile and 0.75 in the strongest. Replicated.* |

*Supplementary Table 12 (continued)*

| Run | Data used | Conditions tested | Main result |
|---|---|---|---|
| Kimi r1 | Released checkpoints for two cell lines, 472 MB each; their held-out ground-truth files, about 1.1 GB each; and released per-perturbation metric tables for 18 evaluations across two datasets and 18,120 perturbation rows [3–5, 7]. Inference was rerun on 12 CPU cores because the available A100 GPU was in use. The reimplementation matched the released tables with rank correlations of 0.97–0.99. | True effect magnitude, measured both as the norm of the true expression shift and as the number of differentially expressed genes; performance relative to predicting no change; a split-half reliability ceiling; predicted shift magnitude; replication across all 18 released evaluations. | **True effect magnitude.** Spearman correlation was 0.734 ($p = 2 \times 10^{-164}$) and 0.813 ($p = 5 \times 10^{-251}$) in the two cell lines, and remained 0.73 and 0.79 after adjusting for cells per perturbation. In the weak quartile, the median variance explained relative to predicting no change was −0.31, and only 4% and 18% of perturbations beat that baseline. In the strong quartile, the corresponding values were 0.50 and 0.74, and 94% and 99% beat the baseline. The reliability ceiling was 0.19 for weak and 0.88 for strong perturbations. STATE captured 37% of the achievable signal for weak perturbations and 92% for strong ones. The predicted magnitude remained about two thirds of the true magnitude even when variance explained was negative, suggesting that weak-response predictions were incorrect rather than simply conservative. *The result replicated in all eight CRISPR evaluations (0.71–0.86) and all five within-context chemical evaluations (0.11–0.44). It was weaker in transfer: two of five effects were not significant, and the relationship largely disappeared because context shift became a larger limit than signal strength.* |
| Kimi r2 | Released transfer checkpoint, containing 49.4M parameters in a 471,699,039-byte file; 16 released result files covering four cell lines, two settings, and 9,881 perturbation evaluations [3–5]; two public screens from a curated repository as independent data—one CRISPR interference screen and one CRISPR activation screen. Both used the same cell line as the checkpoint's held-out context but came from different laboratories [137]. | True effect size in each of eight released settings; new inference on an external screen; a corrupted-label control that assigned the control representation to every perturbation; perturbations absent from the training vocabulary; the discrimination metric. | **True effect size.** Spearman correlation was 0.846, 0.789, 0.836, and 0.793 within context and 0.822, 0.677, 0.763, and 0.387 in transfer, with a pooled value of 0.644. For perturbations affecting ten or fewer genes ($n = 607$), median performance was 0.004 and 47.9% were at or below zero. For perturbations affecting at least 100 genes ($n = 5{,}003$), median performance was 0.423 and 8.4% were at or below zero ($p = 9.8 \times 10^{-234}$). The result replicated on a screen from an external laboratory at $\rho = 0.646$ ($p = 5.6 \times 10^{-8}$), near the low end of the released results as expected for a shallow screen. Advantage over the corrupted-label control increased with effect size at $\rho = 0.652$. *Replicated. Two additional findings were reported. First, perturbations absent from the training vocabulary had median performance of* −0.007 *with no relationship to effect size, because one-hot encoding assigned them the control representation; only 6 of 237 targets in the activation screen were in the vocabulary. Second, the discrimination metric was negatively correlated with effect size at* −0.86*, making the model appear strongest where it had learned the least.* |

**Supplementary Table 13: Level 3 audit detail: AlphaMissense.** Empirical audit with a search for prior publication. GPT r1 and GPT r2 reached different conclusions about whether the cysteine result was already known. Kimi r1 and Kimi r2 used the same main source of labels but reached different conclusions about protein length.

| Run | Data used | Conditions tested | Main result |
|---|---|---|---|
| GPT r1 | All 96 human assays in ProteinGym v1.3 [70]; AlphaMissense scores from the redistributed Zenodo release [50]; AlphaFold DB coordinates for validation [68]; comparison scores from ESM-1b, GEMME, EVE, and ESM2-650M [101, 138, 139]. Proteins were included only when at least 80% of unique substitutions matched the canonical coordinates. This left 269,506 variants across 53 proteins and 66 assays. Discovery and replication were separated by the year in which each protein was first assayed. | Canonical substitutions accessible through a single nucleotide change, which did not pass the initial screen. The outermost 5% of a protein sequence and cysteine-introducing substitutions, which were excluded as already documented; a 2025 rhodopsin mutational-map study reported the same cysteine relationship. The run then screened all 20 destination residues in discovery, selected the strongest remaining negative result before opening the replication data, and applied five robustness restrictions, four comparator models, and a check for measurement noise. | **Methionine as the destination residue.** In discovery, across 24 proteins and 32 assays, performance was 0.284 versus 0.427, a paired difference of −0.142 (CI −0.226 to −0.060, $p = 0.0021$; Bonferroni-adjusted $p = 0.040$ across 20 tests). In replication, across 18 proteins and 22 assays, performance was 0.373 versus 0.485, a difference of −0.112 (CI −0.164 to −0.067, $p = 0.00015$). The result survived all five restrictions, including comparison with destination-specific means, where the differences were −0.115 and −0.079. Greater measurement noise did not explain the result. *Replicated, but not specific to AlphaMissense: all four comparators showed the same deficit, with replication means from* −0.092 *to* −0.150*. The run also confirmed a stronger cysteine result but excluded it because it found that the relationship had already been published.* |

*Supplementary Table 13 (continued)*

| Run | Data used | Conditions tested | Main result |
|---|---|---|---|
| GPT r2 | AlphaMissense authors' supplementary ProteinGym predictions, containing 1,622,429 scores across 87 assays and 72 proteins [50, 70]. The analysis used 353,782 variants across 75 assays and 61 proteins: 17,908 cysteine-introducing substitutions and 335,874 other substitutions. Comparison scores came from ProteinMPNN, GEMME, ESM1b, a position-independent baseline, ESM-IF1, and EVmutation [101, 138, 140–142]. Proteins were assigned to discovery or replication by hashing the UniProt entry name before any outcomes were read, keeping each protein in only one group. | Thirteen conditions defined before analysis and screened together: codon reachability; introduction and removal of cysteine, glycine, and proline; direction of change in volume, hydropathy, and charge; terminal position; local entropy; and local 5-mer uniqueness. Twelve did not pass the discovery screen. The cysteine result was then tested within taxonomic groups, using four versions of the metric, as percentile-rank error, and against six comparators. Tryptophan was also difficult but was not tested in the fixed confirmation analysis. | **Cysteine-introducing substitutions.** In discovery, across 25 proteins, performance was 0.527 versus 0.374, with a reported difference of −0.153 (CI −0.204 to −0.094; Holm-adjusted $p = 2.10 \times 10^{-4}$). The effect was negative in 21 of 25 proteins. In replication on 36 different proteins, performance was 0.537 versus 0.365, with a difference of −0.172 (CI −0.217 to −0.128, $p = 8.38 \times 10^{-9}$), negative in 34 of 36 proteins. The result held in all four taxonomic groups and under AUROC, decile-matched Spearman correlation, and exact position matching. *Replicated, with two qualifications noted by the run. First, the difference reversed when expressed as percentile-rank error. Second, all six comparators showed the same deficit, from* −0.103 *to* −0.168*, compared with* −0.172 *for AlphaMissense. The effect therefore applies to the model class, with AlphaMissense near the severe end. This is the same result GPT r1 excluded as already published.* |
| Opus r1 | Precomputed AlphaMissense predictions [50]; ProteinGym v1, retaining 83 of 96 human assays after exact substring alignment and verification of the wild-type residue at every mapped position, for 308,922 variants [70]; ClinVar dated 2026-07-15, using GRCh38 single-nucleotide variants with at least one gold review star and no conflicts, for 1,429,046 keys [123]; AlphaFold DB v4 and v6 for 67 proteins [68]. | Four conditions excluded as already documented before measurement: disorder and low pLDDT, buried residues and interface positions, gain-of-function and dominant-negative variants, and sensitivity to alignment depth. The run then compared ranking across protein positions with ranking among substitutions at the same position, using two independent label sources. BLOSUM62 and Grantham served as comparators. It also tested whether tolerant positions explained the result and separated results by solvent exposure, pLDDT band, and assay type. | **Ranking within positions versus across positions.** This separates two parts of the prediction task rather than testing a property of the input. Across 71 assays and 16,833 positions, ranking positions achieved 0.51 (CI 0.46–0.56), while ranking different substitutions at the same position achieved 0.27 (CI 0.24–0.29). The difference appeared in 61 of 71 assays (paired Wilcoxon $p = 8 \times 10^{-9}$). On clinical labels with no overlap with the assay data, pairwise accuracy was 0.960 across positions and 0.826 within the same position, so error increased from 4.0% to 17.4%. Removing uninformative positions raised within-position performance only to 0.31. *Replicated. AlphaMissense still outperformed BLOSUM62 and Grantham in the setting where its own performance was weakest. The clinical analysis contained only 353 same-position pathogenic–benign pairs, as the run acknowledged.* |

*Supplementary Table 13 (continued)*

| Run | Data used | Conditions tested | Main result |
|---|---|---|---|
| Opus r2 | Precomputed AlphaMissense scores [50]; ClinVar with review status retained for stratified analysis [123]; UniProt Humsavar as an external label set [126]; gnomAD chromosome 21 extract as a population check [69]. | Gene-level selective constraint, excluded as already documented and used only as a positive control to show that the analysis recovered a known relationship. CpG transitions initially appeared important, but 48% were arginine substitutions and the effect disappeared after removing arginine, so they did not pass the screen. The run then tested aromatic reference residues on separate sets of genes, across ClinVar review levels, on an external label set, with a calibration control, against indicators for the replacement residue, and after removing the worst-performing gene. | **Identity of the reference residue**, meaning the wild-type amino acid being replaced. At the published threshold of 0.564, the false-positive rate was 0.262 (CI 0.234–0.290) for aromatic residues and 0.047 (CI 0.044–0.049) for other residues, a 5.6-fold difference. Rates ranged from about 2% for glutamine to 40% for phenylalanine. The ratio was 5.0 and 4.5 in two non-overlapping gene sets, 4.1 for expert-panel labels, 4.0 in Humsavar, and 5.63 after excluding CFTR, covering 249 genes. After adjusting for the identity of the replacement residue, the direction reversed, showing that the original residue carried the effect. *Replicated at the published operating point. Specificity was 0.953 for other residues but 0.738 for aromatic residues. Restoring 95% specificity for aromatic residues required a threshold of 0.957, which reduced sensitivity by half.* |
| Kimi r1 | ClinVar, with 193,160 labeled missense variants across 1,129 proteins that each had at least five variants in both classes [123]; 69 human ProteinGym deep mutational scanning assays as an independent label source [70]; precomputed AlphaMissense scores [50]. | Four conditions excluded as already documented before measurement: disorder and low pLDDT, solvent accessibility, transmembrane and mitochondrial proteins, and asymmetry in substitution scores. Protein length was then tested in pooled ClinVar, by review status, and in the assay data. For the null assay result, the run calculated the smallest effect that the dataset had enough power to detect. | **Protein length, interpreted as a property of the benchmark rather than the model.** Per-protein AUROC fell from 0.963 for proteins 300–500 residues long to 0.930 for proteins longer than 2,500 residues. Spearman correlation was −0.218 ($p = 1.2 \times 10^{-13}$); specificity fell from 0.925 to 0.857, while sensitivity remained stable. After adjusting ranks for variant count, class prevalence, and label quality, the correlation weakened to −0.071. Both independent checks failed: the relationship appeared only in one-star labels, was flat in two-star labels, was absent in expert-panel labels, and was null in deep mutational scanning (−0.075, $p = 0.54$). *Failed. The run suggested that benign labels in large genes often come from population-frequency rules used by single submitters—the same kind of evidence used to fine-tune the model. It also noted that the assay dataset could detect only a large effect.* |

*Supplementary Table 13 (continued)*

| Run | Data used | Conditions tested | Main result |
| --- | --- | --- | --- |
| Kimi r2 | ClinVar with review levels and submission dates [123]; UniProt HUMSAVAR [126]; ProteinGym [70]; complete proteome-wide set of about 71 million AlphaMissense predictions, used for a check that required no labels [50]. | Disorder was excluded as already documented. It was also measured and showed an effect in the opposite direction, so it could not explain the length result. Protein length was then tested in pooled and per-gene analyses, after controlling label quality and gene composition, at the published thresholds, in three independent label sets, across the full proteome without labels, in the assay data, and against twelve other predictors. | **Protein length.** AUROC was 0.9665 for proteins of at most 2,000 residues, 0.9606 for 2,000–3,000, 0.9399 for 3,000–6,000, and 0.8566 for more than 6,000. Across 528 genes, the per-gene correlation with log length was $\rho = -0.281$ ($p = 4.7 \times 10^{-11}$). The effect remained after restricting to two-star labels, excluding TTN and five other very large genes, restricting to ordered or structured regions, and matching the reference residue. Sensitivity fell from 0.82 to 0.623, and the fraction of ambiguous predictions rose from 5% to 10.3%. *Replicated and supported without labels: across the complete proteome, the ambiguous fraction rose from 13.6% to 18.3%, and the mean score fell from 0.52 to 0.45. A label-submission process cannot produce this label-free pattern. The effect was concentrated above 3,000 residues, a range the other Kimi run did not analyze separately; this explains the different conclusions.* |

**Supplementary Table 14: Level 3 audit detail: CONCH.** Empirical audit with a search for prior publication. All six runs considered effective image resolution or magnification. Three excluded it as already documented, while two treated it as a new finding and reproduced it in two datasets each.

| Run | Data used | Conditions tested | Main result |
|---|---|---|---|
| GPT r1 | Public results by medical center from a 2026 pathology robustness benchmark [73], joined by center to tissue-occupancy measurements from the benchmark's open Camelyon and TCGA tiles. The official CONCH weights require individual license acceptance. The run did not bypass that requirement and recorded this decision. It therefore evaluated CONCH [71] as a fixed patch encoder inside published multiple-instance slide classifiers, using published center-level performance rather than running its own inference. | Stain intensity and color, scanner and site, ten image corruptions, and patch size and context, all excluded as already documented. The run then tested tissue occupancy, defined as one minus the fraction of nearly white pixels. It varied the near-white threshold and excluded the survival endpoint from the pooled fit. | **Tissue occupancy.** Change in AUC per 10 percentage points of occupancy, estimated within each task across 13 task–center observations, was −0.125 across 7 centers for breast homologous-recombination deficiency, −0.114 across 3 centers for Camelyon, and −0.078 across 3 centers for non-small-cell lung cancer. The pooled estimate was −0.114, with Pearson correlation −0.457 and Spearman correlation −0.555. However, the exact stratified permutation test gave $p = 0.131$, the center-level bootstrap interval was −0.350 to 0.137, and the direction reversed to +0.086 when the near-white threshold was 235. *The result weakened and was reported as inconclusive. Its direction—more tissue associated with worse performance—was unexpected. The run suggested that occupancy may represent a center-specific preprocessing pattern rather than the actual amount of tissue.* |
| GPT r2 | Official repository at revision `141cc09c`; checkpoint SHA-256 `40a9644b...` [71]; official colorectal class-name and prompt template ensemble with six exact ontology matches; CRC-VAL-HE-7K for discovery [127, 128]; Stanford STARC-9 for replication [129]. | Nine documented groups of conditions were excluded: stain, contamination, blur, noise, illumination, medical center, tissue type, scale, and resolution. The run then tested distortion caused by forcing rectangular images into a square at five original aspect ratios. It also tested crop direction, results by class, and a threshold-independent outcome. Before scoring, it fixed the confirmation rule: both datasets had to show a negative slope whose two-sided 95% confidence interval excluded zero. | **Distortion from forcing a rectangular image into a square.** Discovery accuracy was 85.6%, 82.2%, 81.1%, 76.7%, and 74.4%. The slope was −2.22 percentage points per 0.1 change in aspect ratio (CI −3.34 to −1.11), and the endpoint difference was −11.11 (CI −16.11 to −6.11). McNemar $p = 8.8 \times 10^{-5}$, and 17.8% of labels changed. Stanford accuracy was 76.7%, 73.9%, 71.1%, 69.4%, and 58.3%. Its slope was −3.29 per 0.1 (CI −4.39 to −2.19), the endpoint difference was −18.33, McNemar $p = 1.0 \times 10^{-8}$, and 25.0% of labels changed. Both crop directions had negative effects, and all four confidence intervals excluded zero. Macro AUROC fell from 0.987 to 0.962 and from 0.978 to 0.885, showing that the result did not come only from the zero-shot decision rule. *Replicated under a confirmation rule fixed in advance, the only CONCH run at this level to do so. The run noted that cropping removes image content while resizing changes geometry, so it could not separate these two explanations.* |

*Supplementary Table 14 (continued)*

| Run | Data used | Conditions tested | Main result |
|---|---|---|---|
| Opus r1 | Unrestricted public mirror `liubw/conch`, with the same 802,235,437-byte size as the restricted checkpoint. The pipeline was validated using the repository's example [71]. Datasets were CRC-VAL-HE-7K; a 495-tile subset of NCT-CRC-HE-100K [127, 128]; an LC25000 lung subset [130]; and PatchCamelyon [131]. | Magnification and resolution, stain normalization, scanner and medical center, and rotation, all excluded as already documented. The run then changed color saturation while preserving brightness. It repeated the test in two datasets, by class, and in a different tissue task. Hue rotation and brightness served as controls. PatchCamelyon was excluded before perturbation because baseline performance was near chance. | **Color saturation.** Balanced accuracy fell from 0.731 to 0.552 across the tested range, a loss of 17.9 percentage points with retention of 0.755. Spearman correlation was 1.00. McNemar $p = 1.3 \times 10^{-32}$, with 239 correct-to-wrong changes and 46 changes in the opposite direction. The result replicated in a separate 495-tile dataset, where accuracy fell from 0.792 to 0.610. Mucus drove much of the result, with recall falling from 0.900 to 0.058, while stroma and background improved slightly. Saturation and hue caused changes of −0.203 and −0.220, compared with −0.033 for brightness, indicating a color-specific rather than general brightness effect. *Replicated, but the size depended on the task. In a lung adenocarcinoma-versus-squamous task, the loss was only 3.3 points and was not significant, although embedding cosine similarity still fell to 0.869.* |
| Opus r2 | The same unrestricted mirror and checkpoint [71]; balanced subsets of CRC-VAL-HE-7K with $n = 72$ [127, 128] and LC25000 with $n = 100$ [130]. | Color and grayscale were measured and then excluded after causing only a 21% loss. Gaussian blur was also excluded because tile-size differences made the datasets difficult to compare. Scale invariance of the embeddings was excluded as already documented, but the earlier evidence used newly trained prediction heads. The run therefore treated the fixed zero-shot pathway as untested. It then measured effective spatial resolution in a second tissue and dataset, tested digital zoom beyond the native magnification range, and analyzed results by class. | **Effective spatial resolution.** Each patch was downsampled to $r \times r$ pixels and then resized to its original size before preprocessing. This kept geometry and field of view fixed while reducing image detail. Balanced accuracy was 0.722 at 20× magnification, followed by 0.667, 0.597, 0.431, 0.306, 0.181, 0.125, and 0.056 at 1.8×. In the lung dataset, accuracy fell from 0.910 to 0.190. Digital zoom beyond the native range also caused a decline, from 0.722 to 0.181, suggesting that the model works best within a limited magnification range rather than always benefiting from more detail. Classes defined by fine structure failed first, while classes defined by coarse texture remained accurate longer. *Replicated. The run considered the result new because published scale-invariance results used adaptable prediction heads, which can retrain on changed embeddings. The zero-shot pathway instead uses fixed text references and cannot adapt in this way.* |

*Supplementary Table 14 (continued)*

| Run | Data used | Conditions tested | Main result |
|---|---|---|---|
| Kimi r1 | The same unrestricted 802,235,437-byte mirror [71]; full NCT-CRC-HE-100K with $n = 100{,}000$ for discovery; CRC-VAL-HE-7K with $n = 7{,}180$ from 50 different patients and no stain normalization for replication [127, 128]; one A100 40 GB GPU. Before any image change, the pipeline achieved 0.7948 (CI 0.7859–0.8041), compared with the published 0.791. | Prompt wording [83], magnification and input resolution, and medical center, stain, and scanner effects, all excluded as already documented. For magnification, the run cited a 2025 study that reported nearly unchanged performance. The run then tested JPEG quality in both datasets and by class, including the unusual behavior of the background class. It did not run a matched-information control with the same peak signal-to-noise ratio. | **JPEG quality.** Balanced accuracy was 0.8331 without compression, 0.7937 at q30, 0.7802 at q25, 0.7227 at q15, 0.6289 at q10, and 0.3257 at q5. Spearman correlation was 0.235 with $p < 10^{-300}$. Three ranges appeared: a plateau at q30 or higher with a loss of at most 4.0 points, a transition from q15 to q25, and a sharp decline at q10 or lower. Macro AUC fell from 0.970 to 0.928 to 0.842. Accuracy at each quality level correlated with discovery accuracy at Pearson 0.987 in the separate-patient dataset. Changes in recall by class at q10 had Pearson correlation 0.946, with the same direction in all nine classes. Background recall was not monotonic: it fell to 0.61 around q60–q75 and rose to 0.976 at q10. At q10, 17–21% of tumor, debris, muscle, and mucus tiles were reclassified as background. *Replicated. Without a control containing the same amount of information, the run could not separate general information loss from artifacts specific to JPEG compression.* |
| Kimi r2 | The same mirror and datasets: NCT-CRC-HE-100K with $n = 100{,}000$ and CRC-VAL-HE-7K with $n = 7{,}180$ [71, 127, 128]. Baseline balanced accuracy in the replication dataset was 0.7926, compared with the published 0.791. | Stain and scanner color effects and prompt sensitivity, excluded as already documented. An initial screen ranked four image changes by their damage in a 10,000-tile subset: downsampling −67 points, compression −53, noise −50, and blur −35. Only downsampling was selected for further testing; the others were not tested in an independent dataset. Effective resolution was then tested in both datasets, with three resizing methods, against matched Gaussian blur, by class, and against a known compression artifact in discovery. | **Effective spatial resolution.** Downsampling factors of 1, 1.5, 2, 3, 4, 6, 8, and 16 produced accuracies of 0.8119, 0.8023, 0.7603, 0.6134, 0.4065, 0.2044, 0.1209, and 0.0493. Spearman correlation was −1.000, and every adjacent change was significant. The largest change occurred from $s = 3$ to 4, with 21,586 correct-to-wrong changes and 891 in the opposite direction. In replication, accuracy fell from 0.7926 to 0.0294, Spearman correlation was −1.000, and every adjacent change except 1 to 1.5 was significant. Performance was relatively stable at or below 1.0 $\mu$m pixel$^{-1}$, with a loss of at most 5.2 points. It then fell by 32–35 points as resolution changed from 1.0 to 2.0 $\mu$m pixel$^{-1}$ and failed above 3.0. At $s = 4$, bilinear, bicubic, and nearest-neighbor resizing produced 0.397, 0.406, and 0.285. Gaussian blur at $\sigma = 2$ produced 0.447. This indicates loss of high-frequency image detail rather than an artifact of one resizing method. *Replicated. This run considered the resolution result new for CONCH. It distinguished earlier work on low-resolution natural images, research that measured embedding invariance rather than accuracy, and a study that trained a separate model. The other Kimi run made the opposite decision about whether the same issue was already known.* |

**Supplementary Table 15:** **Level 3 audit detail: ESMFold2.** Empirical audit with a search for prior publication. Three runs evaluated the 2026 release, while three evaluated the 2022 predecessor. Because the relevant prior literature differs, every row identifies the system tested.

| Run | Data used | Conditions tested | Main result |
| --- | --- | --- | --- |
| GPT r1 | *2026 release through a hosted service.* Public endpoint of the release's hosted atlas [97, 101], queried using the MD5 checksum of the exact sequence; FoldBench for discovery, with 309 of 334 attempted sequences retained [98]; 150 structures first released by RCSB in 2025 for replication, determined by X-ray crystallography at 2.5 Å resolution or better [133]. The endpoint did not report the checkpoint hash, random seed, loop count, or use of alignments, so its exact configuration was unknown. | Nine conditions excluded as already documented in the model's preprint or limitations page: pseudo-perplexity, alignment depth, homology and temporal holdout, length and long-context scaling, modality, inference budget, confidence calibration, underrepresented lineages, and short or disordered proteins. The run then tested glycine fraction, controlling for length and the rest of the amino-acid composition. It also tested a linear form of the relationship and folded isolated constructs within their natural full-length sequences. | **Glycine fraction.** Spearman correlation with C$\alpha$-lDDT [136] was 0.199 (CI 0.087–0.309; permutation $p$ = 0.00050), which survived Bonferroni correction across 65 screened conditions. Median lDDT was 0.946 above and 0.925 below 7% glycine. The result replicated in the 2025 sample at 0.203 (CI 0.034–0.356, $p$ = 0.0128) and remained 0.193 and 0.187 after controlling for length. *The glycine-specific effect did not replicate after adjusting for the other amino acids: the correlation was 0.175 in discovery and 0.004 in replication. The run therefore reported a general relationship with amino-acid composition rather than a mechanism specific to glycine. Placing constructs in their natural full-length sequence also failed to improve accuracy and slightly reduced it.* |

*Supplementary Table 15 (continued)*

| Run | Data used | Conditions tested | Main result |
|---|---|---|---|
| GPT r2 | *2026 release run locally.* Checkpoint revision `1ebf0e34`, weight SHA-256 `138fd435...`, language-model component `45b0fa5d`, and inference library `ba4d7124` [97]; discovery and replication datasets containing 50 short and 50 control targets from 2024 and 2025 RCSB releases [133]; four additional datasets for testing sensitivity to the checkpoint configuration; MMseqs2 for homology control [135]. | Three initial ideas were abandoned. A difference in composition between the two halves of a sequence disappeared after shuffling while preserving length and composition. A viral/disulfide condition failed in the independent dataset. An envelope-like disulfide and hydrophobic condition was explained by two shared alphavirus clusters. The run then tested sequences shorter than 200 residues, the boundary of the distillation dataset that supplies 70% of structural training. It tested coordinate accuracy, confidence, a continuous length effect, homology control, four checkpoint configurations, and the practical effect of a confidence filter. | **Sequences shorter than 200 residues: calibration rather than accuracy.** Coordinate accuracy showed no effect: −0.0048 (CI −0.0176 to +0.0112) in 2024 and +0.0022 (CI −0.0114 to +0.0201) in 2025. Confidence did change. The pLDDT differences were −0.0799 (CI −0.1266 to −0.0496) and −0.0526 (CI −0.0780 to −0.0165), while the confidence–accuracy gap shifted by −0.0677 and −0.0507. Spearman correlation between $\log_2$ length and this gap was 0.567 ($p = 7.97 \times 10^{-10}$) and 0.449. After removing every replication target that shared a sequence cluster across years, the confidence shift remained −0.0524 across 88 targets, while coordinate accuracy remained unchanged. All four released-checkpoint datasets showed the same direction. *Replicated. In practice, a pLDDT threshold of 0.9 rejected 70.5% of accurate short targets but only 19.0% of accurate controls.* |
| Opus r1 | *2022 predecessor.* `facebook/esmfold_v1` [101]. The run reported that it could not find a separately released model with the name in the task. It used 900 recent structures released from 2024-01 to 2026-04 and 400 early structures from 2010–2011. Each group contained no pairs above 30% identity, and every early structure with a match to a recent structure was removed. This reduced 1,788 candidates to 1,238 [133]. ESM-2 650M and 3B were used to calculate pseudo-perplexity [101]. | Language-model pseudo-perplexity in both time periods; length, contact density, secondary structure, and resolved coverage as possible confounds; predicted confidence as a possible explanation; native contact density; release period; comparison between the 650M and 3B language models; enrichment of designed proteins among high-perplexity sequences. | **Language-model pseudo-perplexity.** Spearman correlation with TM-score [134] was −0.397 ($p = 3 \times 10^{-35}$) in the recent sample and −0.294 ($p = 2 \times 10^{-9}$) in the homology-filtered early sample. It remained between −0.387 and −0.392 after adjustment. Performance stayed near TM-score 0.95 until perplexity reached about 7 and then declined sharply. Mean TM-score fell from 0.946 to 0.806 across quartiles, while failure increased from 1.8% to 22.2%. *Replicated, but already documented. This run did not perform the prior-publication screen, while two other runs explicitly excluded pseudo-perplexity as known. A secondary finding may be newer: native contact density had a raw correlation of* +0.414 *and changed the odds of failure by 16.7- to 23.3-fold. Unlike pseudo-perplexity, it remained* +0.303 *and* +0.372 *after controlling for pLDDT and did not appear on either exclusion list.* |

*Supplementary Table 15 (continued)*

| Run | Data used | Conditions tested | Main result |
|---|---|---|---|
| Opus r2 | *2022 predecessor.* RCSB monomeric X-ray structures at 2.0 Å resolution or better and 80–300 residues long. After selecting one representative per 30%-identity cluster, 5,689 structures were shuffled with seed 1234 and split into two separate halves; 450 were sampled from each [101, 133]. | Perplexity, length, and pLDDT were intentionally recovered to validate the analysis and then excluded as already documented; their correlations with TM-score were −0.38, +0.17, and +0.66. The run then tested elongation in both halves, local versus global accuracy, confidence, a mechanism check that did not depend on an accuracy metric, secondary structure, contact order, and low complexity as alternative explanations, and very elongated fibrous proteins in relation to prior research. | **Elongation**, measured as the observed radius of gyration divided by the expected radius for a globular chain of the same length. Partial Spearman correlation with TM-score was −0.35 after controlling for length and perplexity, and −0.27 after also controlling for pLDDT; both had $p < 10^{-8}$. The multivariable coefficient was −0.21 ($p = 4.5 \times 10^{-17}$). Failure increased from 5% in the most compact group to 33% in the most elongated group. The relationship replicated in the separate half at −0.31 and −0.33. Local structure remained accurate: C$\alpha$-lDDT changed only from 0.91 to 0.89, while the global TM-score fell from 0.94 to 0.78. Controlling for lDDT strengthened the TM-score relationship to −0.45. Confidence changed little, from 86 to 82, so it did not reveal the failure. *Replicated. A metric-independent check showed the same mechanism: predicted elongation regressed toward the mean, with slopes of 0.80–0.88, and 62% of predictions were more compact than their native structures.* |
| Kimi r1 | *2022 predecessor.* `esmfold_v1` through fair-esm 2.0.0, run as a single sequence without alignments or templates on one A100 40 GB GPU [101]. The evaluation used two datasets of X-ray structures at 2.5 Å resolution or better, 50–350 residues long, clustered at 30% identity. Each dataset contained 316 analyzed structures: discovery structures were deposited in 2022, and replication structures from 2024-07 to 2025-07 [133]. All 640 folding runs finished in 54 minutes. | Chain length and similarity to the training data by identity and $k$-mer coverage, excluded as already documented. An initial composition screen covered low complexity, charge, aromatic residues, glycine and proline, repeats, hydropathy, and secondary structure; none passed. Relative contact order also had no relationship with lDDT. The run then tested the aspect ratio of the C$\alpha$ gyration tensor, separating error by residue distance, examining confidence, excluding repeat-rich sequences, and testing whether predicted shape could flag risk during inference. | **Gyration-tensor aspect ratio.** Spearman correlation with TM-score was −0.341 ($p = 4.7 \times 10^{-10}$) in discovery and −0.324 ($p = 3.6 \times 10^{-9}$) in the separate replication sample. Partial correlations were −0.280 and −0.300, and the pooled correlation was −0.331 ($p = 1.4 \times 10^{-17}$). Failure occurred in 22.8% of the elongated half and 5.7–10.1% of the compact half. Out-of-sample $R^2$ was 0.733, compared with 0.758 in sample. Local pair error at residue separations of 4–12 remained stable, while long-range error at separations above one third of the chain increased at +0.212 and +0.224. Confidently wrong predictions, defined as pLDDT at least 80 and TM-score below 0.8, were about ten times more common in the elongated half. The result remained −0.286 and −0.298 after removing repeat-rich sequences. *Replicated and usable during inference: predicted and native aspect ratios correlated at 0.78, allowing risk to be flagged with a predicted aspect-ratio threshold near 2.5.* |

*Supplementary Table 15 (continued)*

| Run | Data used | Conditions tested | Main result |
|---|---|---|---|
| Kimi r2 | *2026 release in its fast configuration.* Snapshot `b28d8ace`, with three recurrent loops, 50 sampling steps, and one diffusion sample, run on one A100 40 GB GPU [97]. Cohort A contained 1,293 scored structures from 2017-01-01 to 2021-06-30, within the training period. Cohort B contained 1,291 scored structures from 2023-01-01 to 2024-12-31, after training. Both used X-ray structures at 2.5 Å resolution or better, 40–500 residues long, with one representative per 40%-identity cluster [133]. A stricter replication set contained 1,159 structures after removing every cohort B cluster represented in cohort A. The design was fixed before analysis. | Length, alignment depth, disorder and low complexity, repeated single-residue runs, and confidence calibration, all excluded as already documented. Low complexity, repeated residues, and sequence entropy were also nonsignificant in this dataset. The run then screened 31 sequence-only conditions using Benjamini–Hochberg correction. Seven passed, and the other 24 were not tested in replication. Glycine was selected for analysis with confounder adjustment, dose–response modeling, residue-level localization, robustness across tails and subgroups, and a check for other small amino acids. | **Glycine fraction.** In cohort A, the correlation was +0.160 ($p = 7 \times 10^{-9}$, CI +0.104–+0.214), with a slope of +0.40 lDDT points per 0.01 increase in glycine fraction. It replicated at +0.143 in the post-training sample and +0.136 in the strictly cluster-separated sample. After adjusting for length, experimental resolution, observed fraction, and experimental secondary structure, the effect remained between +0.111 and +0.147. Glycine was negatively correlated with helices and positively correlated with sheets, the opposite of what would create the observed result as a secondary-structure artifact. The bottom glycine decile had mean lDDT of 0.862–0.889 and 12–16% of targets below 0.80, compared with mean lDDT of 0.92–0.94 and 3–5% below 0.80 at 8–10% glycine. *Replicated, with four limitations noted by the run: the direction was unexpected, the datasets contained only crystallizable ordered single-chain proteins, the model used a fast single-sample configuration, and the effect was modest. Alanine also replicated, and glycine correlated with mean residue mass at −0.65, suggesting that the broader factor may be small amino acids rather than glycine alone.* |

**Supplementary Table 16: Level 3 audit detail: STATE.** Empirical audit with a search for prior publication. Three runs reanalyzed the released evaluation tables, and three downloaded the checkpoints and ran inference.

| Run | Data used | Conditions tested | Main result |
|---|---|---|---|
| GPT r1 | *Released outputs only.* Best public few-shot checkpoint for the chemical atlas, its configuration, and official per-perturbation evaluation tables [3, 7], verified by SHA-256. Each held-out cell-line file was 65–75 GB, while the workspace had 553 MB, so inference was not rerun. The configuration represents each drug-and-concentration pair as a separate one-hot category; the model does not treat dose as a continuous value. The analysis used 199 drugs measured at both 0.05 and 5 $\mu$M in every cell line, with one melanoma line for discovery and four others for replication. | Nominal concentration, tested with delta mean absolute error and its squared-error version; delta correlation; agreement on the direction of differential expression; biological effect size at the high dose, retained as a known confounder. | **Nominal concentration.** Delta mean absolute error was 0.00701 at 0.05 $\mu$M and 0.00947 at 5 $\mu$M, a ratio of 1.351 (CI 1.244–1.460). Error was worse at the high dose for 125 of 199 drugs (sign-test $p = 0.000368$). In four additional cell lines, Holm-corrected ratios were 1.275, 1.206, 1.266, and 1.298; all were significant, and 59.8–68.8% of drugs were worse at high dose in every line. Error increased monotonically across all three doses. Delta correlation was nearly unchanged, suggesting that the model identified which genes changed but misestimated the size of the change. *Replicated, with an important confound: the number of truly differentially expressed genes increased by 46–66 at the high dose, making the high-dose conditions intrinsically more difficult.* |
| GPT r2 | *Released outputs only.* The same official few-shot evaluation tables [3, 7]; molecular weights from PubChem, matched by exact benchmark name for 369 of 379 compounds [143]. Dose was fixed at 0.5 $\mu$M. Compounds were divided by the parity of a hash of the drug name, and the two groups were evaluated in separate cell lines. Replication therefore used both different compounds and different biological contexts. | Cell-set size and nominal dose, excluded as already documented; dose was also considered too similar to perturbation strength. The run then tested molecular weight in both partitions, adjusted for biological effect size, fixed thresholds using the discovery group, tested a separately trained zero-shot checkpoint, and excluded salt forms and very high molecular weights in sensitivity analyses. | **Molecular weight at a fixed dose.** Spearman correlation with delta correlation was −0.181 (CI −0.340 to −0.017, $p = 0.0315$) across 139 compounds in three discovery cell lines and −0.181 (CI −0.338 to −0.017, $p = 0.0473$) across 121 compounds in two different replication lines. After adjusting for observed effect size, correlations strengthened to −0.211 and −0.208, so the result was not simply caused by stronger or weaker drug effects. With the discovery thresholds held fixed, performance was 0.948 below 298 g/mol and 0.929 above 490 g/mol in discovery, and 0.946 versus 0.925 in replication. The direction was negative in all five cell lines. *Replicated, but limited to one checkpoint. The pooled correlation was* −0.004 *for the separately trained zero-shot checkpoint, so the relationship applied to the underrepresented-context checkpoint rather than the entire released model family.* |

*Supplementary Table 16 (continued)*

| Run | Data used | Conditions tested | Main result |
|---|---|---|---|
| Opus r1 | *Checkpoints rerun.* Two released few-shot checkpoints: one trained on a CRISPR-interference atlas across four cell lines and one on cytokine-stimulated memory B cells [3–6], run with the official inference command. Each control cell's expression profile was converted back to counts by binomial thinning and passed through the same normalization at retained fractions from 1 to 0.01. Only control cells were changed; the ground truth remained at its original sequencing depth. | Training-time cell-set size and inference-time set size, measured and then excluded as two versions of the same known relationship. The run then tested reference-cell sequencing depth in four CRISPR lines and in an independent assay and tissue, combined results by absolute depth, tested the full released metric set, separated bias from residual error, used three thinning seeds, and validated its pipeline against the official scorer. | **Reference-cell sequencing depth.** As depth fell from its original value to about 34 UMI per cell, delta correlation fell from 0.373 to 0.299, 0.346 to 0.328, 0.515 to 0.372, and 0.502 to 0.415 across the four lines. Mean absolute error increased 4.4- to 5.0-fold, and paired tests gave $p < 10^{-10}$ in three of four lines. In memory B cells, which began at about 390 UMI per cell, delta correlation fell from 0.601 to 0.308 and error increased 11.6-fold. Across all five contexts, absolute read depth mattered more than the fraction removed: performance retained at least 95% of its original level down to about 250 UMI per cell and fell below 90% under about 74 UMI. Every metric in the released suite was affected. *Replicated; variation across thinning seeds was about 0.003, compared with effects of 0.06–0.29.* |
| Opus r2 | *Checkpoints rerun.* Two per-cell-line checkpoints from the same CRISPR atlas, with public raw counts rebuilt from the original files [3–5]. Processing reproduced the model's steps: library-size normalization, log transformation, and selection of its exact 2,000 genes. Only control cells were downsampled and renormalized, and predictions were compared with a real reference kept at full sequencing depth. | The claim that normalization absorbs sequencing depth, which the paper assumed but did not test, was treated as already documented. The run then tested depth in one in-distribution line and a separate per-cell-line checkpoint, recalculated the same outcome using a reference built from degraded cells, tested rank-based discrimination, and reproduced the published per-perturbation metrics at full depth. | **Reference-cell sequencing depth.** At a median depth of 11,674 UMI per cell, delta correlation was 0.483 and error was 0.063. At 2,920 UMI, they were 0.421 and 0.072; at 584 UMI, 0.254 and 0.133. This was a 47% decline in correlation and a 2.1-fold increase in error, following a log-linear relationship with $R^2 = 0.88–0.94$. The result replicated in a separate checkpoint across seven depth levels and 400 perturbations: delta correlation fell from 0.288 to 0.156, while error increased 3.67-fold. *Replicated. When the reference was recomputed from the same degraded cells, the metric remained nearly flat, from 0.498 to 0.479; it declined sharply only when compared with a fixed full-depth reference. Normalization removes differences in total scale but cannot restore information lost at low sequencing depth. This explains why a pipeline can appear insensitive to depth even when its predictions are not.* |

*Supplementary Table 16 (continued)*

| Run | Data used | Conditions tested | Main result |
|---|---|---|---|
| Kimi r1 | *Released outputs only.* Evaluation outputs for three CRISPR cell lines, an embedding-space version, memory B cells in both settings, and chemical-atlas tables for screening [3–7]. The analysis used only a CPU and took less than one core-hour. Before measurement, the run inspected the data loader and prediction writer. It verified that the same control-cell index generated both the input embedding and recorded barcode, with rows kept aligned. This allowed legitimate pairing by cell even though the files were organized as aggregate outputs. | Performance versus true effect size and performance versus on-target knockdown efficacy, both excluded as already documented; knockdown efficacy had also been largely filtered out before the released data were distributed. Within-drug dose did not pass the screen after accounting for effect size. The run then tested per-cell input sensitivity in six evaluations, reconstructed control rows, checked marginal calibration, and examined where performance crossed a simple baseline. | **Per-cell input sensitivity.** This was measured as the slope of the predicted cell's deviation from its perturbation average against the matching input cell's deviation from the control average. A faithful cell-specific map would have a slope near 1; a population-level sampler would have a slope near 0. In discovery, the slope was 0.0029 (CI 0.0024–0.0039) across 945 perturbations. *The run reported replication of low input–output sensitivity in five additional conditions: Jurkat (slope* −0.0118*), K562 (0.0002), an alternative model variant (0.0025), and independent Parse data under few-shot (0.135) and zero-shot (0.103) evaluation. Paired pre- and post-perturbation biological ground truth was unavailable, so these results do not establish true per-cell predictive accuracy. A slope near 1 represents preservation of input-cell deviations, not a universally required property of a biologically correct perturbation response.* |

| | | | |
|---|---|---|---|
| Kimi r2 | *Checkpoints rerun.* All four released few-shot CRISPR checkpoints, covering 4,058 held-out perturbations across four cell lines [3–5], with package versions fixed. Public raw counts were processed independently, and the authors' released predictions were downloaded as a cross-check. | Effect size and differentially expressed gene count, zero-shot and few-shot context transfer, cell-set size, architecture comparisons, and baseline comparisons, all excluded as already documented. Baseline expression of the perturbed gene did not pass the screen: its effect fell from +0.168 to +0.019 after controlling for effect size and was reproduced by a simple baseline. The run then tested whether the model captured the directly targeted gene in four cell lines, compared seen and unseen perturbations, measured a downstream consequence, checked the authors' own predictions, and tested consistency across cell lines. | **On-target capture**, measured as the slope of predicted versus true log fold change for the gene directly perturbed. The true median was −0.44, while the predicted median was −0.011. The slope was 0.045 (CI 0.027–0.064; Wilcoxon $p = 1.9 \times 10^{-29}$), and was 0.067, 0.105, and 0.094 in the other three cell lines. The predicted absolute fold change of the target gene ranked near the 34th percentile among all genes, while its true change ranked above the 99.9th percentile. This occurred even though the same predictions captured 87% of the overall response norm. Performance was better for perturbations seen during training but remained weak: 0.170 for seen versus 0.045 for unseen perturbations. Among 266 genes held out in at least two cell lines, 79% had median capture below 0.15. *Replicated and confirmed in the authors' released predictions, where median predicted target log fold change was* −0.008 *versus a true value of* −0.439. *The result therefore reflects the model rather than this run's inference implementation.* |

**Supplementary Table 17:** **Level 4 audit detail: AlphaMissense.** AutoSPECTRA-guided. Each run fixed a condition before scoring, tested a model-independent baseline on the same variants, and made an explicit decision. All runs reanalyzed released score tables because the model weights are not public.

| Run | Data used | Conditions tested | Prespecified condition, result, and decision |
|---|---|---|---|
| GPT r1 | ClinVar [123] joined to the AlphaMissense hg38 release [50] using exact GRCh38 coordinates and GENCODE v32 transcripts [144]: 200,901 variants across 14,119 genes. Genes were assigned by hash to discovery, with 84,436 variants, or confirmation, with 116,465 variants, with no genes shared between them. **Baseline:** BLOSUM62 [145]. | Disorder was excluded because it was already documented. Interface position and gene evolutionary age were deferred without a conclusion. The fixed condition was also tested with calibration outcomes, exact matching on gene and substitution, covariate adjustment, ClinVar review levels, and a separate ProteinGym dataset. | **Distance to the nearest internal exon boundary.** Four levels were fixed by hash, with median distances of 135, 33, 11, and 1 bp. Confirmation AUROC was 0.9602, 0.9592, 0.9589, and 0.9168, a decline of 0.0435 (CI 0.0306–0.0572). Brier score increased from 0.0643 to 0.1303, showing that the result did not depend only on the classification threshold. *The decline was also observed in the baseline: BLOSUM62 declined by 0.0432, only 0.0002 different from AlphaMissense and within the prespecified* ±0.03 *margin. On ProteinGym, the same substitutions did not decline (*−0.0208*), and residues near exon boundaries had higher AlphaFold confidence. The clinical decline therefore appears to reflect an unmodeled splicing mechanism rather than more difficult amino-acid chemistry.* |
| GPT r2 | The same two sources acquired independently [50, 123], with GENCODE 32 exon features [144]: 182,240 variants across 13,913 genes. Hash partitioning produced 71,333 discovery and 110,907 confirmation variants with no shared genes. **Baseline:** BLOSUM62 [145]. | Disorder was excluded as already documented. Interface and binding positions were tested in a separate ProteinGym dataset [70] and rejected when the direction reversed across two confirmation assays. Gene age was deferred because it depended on the choice of biological ontology. | **Minimum exonic distance to an internal exon boundary.** Median distances across the four groups were 84, 20, 7, and 1 nt. The discovery difference was 0.03818, below the protocol's 0.05 requirement. Before opening the confirmation data, the run fixed an additional statistical-power rule, which the result met. Confirmation AUROC was 0.9662, 0.9625, 0.9624, and 0.9184. The difference between L3 and L0 was −0.04788 (CI −0.06159 to −0.03549): performance was stable through L2 and fell only at L3. The result remained after standardizing amino acid identity, separating review levels and protein-length quartiles, and using a within-gene analysis across 184 genes. *The decline was also observed in the baseline: BLOSUM62 declined by 0.05984, and the paired difference between models included zero.* |

*Supplementary Table 17 (continued)*

| Run | Data used | Conditions tested | Prespecified condition, result, and decision |
|---|---|---|---|
| Opus r1 | 271,201 variants across 73 ProteinGym assays and 64 human proteins [50, 70], without clinical labels. Alignments were built with phmmer against SwissProt [126] to measure coverage, column entropy, and a position-specific scoring matrix (PSSM). **Baselines:** BLOSUM62 [145] for amino-acid chemistry and the PSSM for evolutionary conservation. | Whole-protein alignment depth was invalid as originally defined: accuracy rose as depth fell ($\rho = -0.30$) because stability assays had both the deepest alignments and the lowest accuracy. This required analysis within each assay. Coverage at the variant's position showed a real relationship, but not one specific to AlphaMissense. | **Evolutionary constraint at the variant position**, measured within each assay as negative alignment-column entropy. The ordering was verified at 2.92, 2.17, and 1.08 bits. AlphaMissense performance fell from 0.410 to 0.291, a decline of +0.119 (CI +0.078–+0.161). The decline was positive in 50 of 70 assays ($p = 1.1 \times 10^{-6}$) and replicated in protein-separated groups at +0.138 and +0.099. *The decline was larger than for the chemistry baseline by* +0.057 *(CI* +0.012–+0.102*), but was indistinguishable from the PSSM by* −0.018*. The PSSM itself outperformed the chemistry baseline by* +0.075*. AlphaMissense therefore added no robustness beyond a conventional evolutionary profile where evolutionary information was weak.* |
| Opus r2 | The same dataset as Opus r1 [50, 70], using different supporting evidence: the ProteinGym multiple-sequence-alignment archive and AlphaFold residue-level confidence for 61 of 64 proteins from the prediction API [68]. **Baselines:** BLOSUM62 [145] and a PSSM. | The run independently reproduced the reversal for whole-protein alignment depth at $\rho = -0.297$, confirming that the original definition was invalid. Coverage was fixed first and showed a real decline, from 0.381 to 0.245, but confidence explained it in a joint model: confidence remained at $t = 8.26$, while coverage became nonsignificant. The run therefore reassigned the result and fixed a second condition. | **AlphaFold residue-level confidence.** Four levels were fixed again, with means of 94.7, 83.8, 61.3, and 39.9. AlphaMissense performance was 0.419, 0.319, 0.175, and 0.143, a paired decline of +0.297 (CI +0.207–+0.386). The decline appeared in 24 of 27 assays ($p = 3.1 \times 10^{-6}$) and replicated in protein-separated groups at +0.208 and +0.359. Scores in disordered regions were compressed toward benign, with a mean of 0.29 versus 0.67. *The decline exceeded the chemistry baseline by* +0.134 *(*$p = 0.002$*) but was shared with the PSSM; their difference was* +0.024 *(*$p = 0.46$*). The run also acknowledged that this condition had already been documented but was missed during its novelty search.* |

*Supplementary Table 17 (continued)*

| Run | Data used | Conditions tested | Prespecified condition, result, and decision |
|---|---|---|---|
| Kimi r1 | ClinVar GRCh38 VCF, reduced to 164,792 variants after joining AlphaMissense scores and verifying every wild-type residue [50, 123]; GENCODE v50 exon coordinates [144]; ortholog counts from Ensembl BioMart [146]. **Baselines:** BLOSUM62 [145] and Grantham [124]. | Four tests with sufficient statistical power were carried to a decision: local sequence complexity, distance to the nearest protein end, protein length, and ortholog count. All four ran opposite to the predicted direction: AlphaMissense improved by 0.0088–0.0161, while the fixed baselines declined on the same data. Population observability was reported only as exploratory. | **Nucleotide distance to the nearest exon boundary.** Four groups were defined from exon coordinates before scoring. AUROC declined monotonically from 0.9679 to 0.9639, 0.9609, and 0.9252. The contrast was 0.0427 (CI 0.0355–0.0504, $p < 10^{-4}$). It increased to 0.0658 among variants with two review stars and remained within every Grantham third and entropy quartile. *The decline was smaller than in both baselines: BLOSUM62 declined by 0.0541 and Grantham by 0.0598 on the same variants. The four sufficiently powered negative tests declared first reduce the chance that the successful condition was selected only because it happened to look favorable.* |
| Kimi r2 | ClinVar dated 2026-07-20, with 209,344 variants, including 67,958 pathogenic [50, 123]. Features were created before scoring and did not use model outputs: phyloP [147], Grantham [124], AlphaFold DB v6 confidence [68], and CpG context. **Baselines:** Grantham and phyloP in the same paired analysis. | Pooled conservation across genes was invalid as originally defined: accuracy appeared higher at low conservation, but the direction reversed after separating genes. The run then completed four sufficiently powered negative tests: within-gene conservation, gene-level mean conservation, Grantham severity, and CpG context. Each was null for AlphaMissense even though one baseline declined. | **AlphaFold residue-level confidence within each gene.** Four levels, the within-gene rule, and the estimator were fixed together before testing 365 genes. The paired contrast was +0.115 (CI +0.087–+0.144), above the prespecified +0.05 threshold. Across 41 genes with complete data, the more detailed curve was 0.856, 0.959, 0.939, and 0.954. The result remained within both conservation groups. *The decline was shared with the chemistry baseline (*+0.097*) and much larger than for the conservation baseline (*+0.010*). The run noted that residue-level confidence was already documented; its new contribution was the within-gene analysis, because the pooled contrast was only* +0.009 *while the paired within-gene contrast was* +0.115. |

**Supplementary Table 18: Level 4 audit detail: CONCH.** AutoSPECTRA-guided. All runs evaluated zero-shot classification of nine colorectal tissue classes against an NCT-CRC-HE-100K training reference. Five runs could transform and rescore images; one could only analyze a fixed table of predictions.

| Run | Data used | Conditions tested | Prespecified condition, result, and decision |
|---|---|---|---|
| GPT r1 | Licensed checkpoint at revision `8391de42`, verified as 802,235,437 bytes before scoring [71]; new inference on 7,180 CRC-VAL-HE-7K images and 12,000 Stanford STARC-9 images [127–129]. A rule fixed in advance required 50 images per class at every level. Four CRC classes, with 3,514 images, and five Stanford classes, with 9,951 images, met it. **Baselines:** two classifiers trained only on NCT, one using handcrafted features and one using MobileNet [148]. | Joint color-and-texture distance was separated before scoring. Only texture was retained because stain was a documented control. Stain, blur, tissue fraction, and JPEG compression were included as measured nuisance variables. Kather-2016 was rejected before scoring the target model because no class met the sample-size rule and the condition was strongly mixed with focus ($\rho = -0.711$). | **Texture-only distance from the NCT reference.** The condition was fixed by hash, and its ordering passed from 0.232 to 0.639. CRC balanced accuracy was 0.627, 0.636, 0.643, 0.619, and 0.574. The decline was 0.0537 (CI −0.0056 to 0.1129), with one-sided $p = 0.0415$ against a prespecified 0.025 threshold. After exact matching on class, stain, focus, and tissue fraction, the direction reversed to −0.0346, and only two of four classes declined. Stanford showed an unadjusted decline of 0.0253 and a matched decline of 0.0115. *Both baselines declined more—0.0873 and 0.1959 versus 0.0537 for CONCH. The condition separated the images, but CONCH was not the model most harmed by it.* |

*Supplementary Table 18 (continued)*

| Run | Data used | Conditions tested | Prespecified condition, result, and decision |
|---|---|---|---|
| GPT r2 | The same checkpoint [71]; CRC-VAL-HE-7K as the main dataset and the normalized Stanford archive for confirmation [127–129]; a deterministic, class-balanced NCT reference containing 4,860 images, with 540 per class. The smallest detectable declines were 4.18 and 2.80 percentage points; a 5-point decline was defined as meaningful before scoring. **Baselines:** three models trained only on NCT: handcrafted nearest-centroid, handcrafted logistic regression, and ImageNet ResNet-18 with an NCT-trained head. | Magnification was closed without a test because the external archive had no valid per-image resolution field. The run did not replace a true acquisition setting with a value inferred from the image. Stain, blur, tissue fraction, and compression were included as adjusted covariates and excluded from construction of the condition. | **Joint color-and-texture distance from the NCT reference.** Five equal-sized levels were fixed by hash. CRC accuracy was 0.793, 0.786, 0.809, 0.775, and 0.714, a decline of 7.88 percentage points (CI 3.86–11.63). Stanford accuracy was 0.681, 0.702, 0.654, 0.608, and 0.586, a decline of 9.49 points (CI 6.72–12.20). Both lower confidence limits were above zero and both exceeded the prespecified 5-point threshold. *The decline was confirmed independently, but the baseline comparison depended on the dataset. In CRC, the handcrafted logistic model declined by 34.62 points versus 7.88 for CONCH. In Stanford, CONCH declined by 9.49 points versus 5.15 and 6.33 for the baselines.* |
| Opus r1 | No checkpoint was available. CONCH predictions came from a per-image table, fixing the analyzed population at 7,180 images [71, 127, 128]. A bounded, class-stratified sample of the 11.69 GB NCT archive was downloaded through HTTP range requests, with 300 images per class. **Baseline:** a classifier trained on this reference sample using the color-feature family. | Overall distance without class adjustment was invalid: accuracy increased with distance ($\rho = +0.082$) because color distance also identified tissue class. After class balancing, the curve changed to a monotonic decline. Image sharpness was tested in parallel and controlled against the fixed condition in both directions. | **Color and stain distance from the reference within each tissue class.** The data were divided into thirds so that all nine classes contributed to every level. Macro accuracy was 0.848, 0.785, and 0.723, a decline of 0.125, or 0.112 after removing the relationship with blur. The class-adjusted odds ratio was 0.13, compared with 2.03 without adjustment ($p = 1.4 \times 10^{-38}$). The result replicated in a held-out half using thresholds fixed in discovery. *The effect was limited. Only debris and stroma remained after blur adjustment, and the baseline declined more than CONCH, 0.177 versus 0.112. Image sharpness had the larger effect in this dataset, at 18.6 points, and was reported as image corruption rather than novelty relative to training.* |

*Supplementary Table 18 (continued)*

| Run | Data used | Conditions tested | Prespecified condition, result, and decision |
|---|---|---|---|
| Opus r2 | Licensed CONCH checkpoint and fresh zero-shot inference [71]. The 7,180 CRC-VAL-HE-7K tiles were used only for development. Confirmation used 1,080 NCT-CRC-HE-100K tiles, 120 per class, acquired from a Hugging Face mirror [127, 128]. A separate 2,700-tile NCT sample trained a fixed ImageNet ResNet-50 plus logistic-regression baseline. Confirmation tiles were excluded from baseline fitting; patient-level separation between these subsets could not be verified. | Development screening and a blur pilot selected a known image-quality axis. The analysis plan, code, prompts and weights were frozen before a single confirmation execution. Each confirmation tile was evaluated at Gaussian-blur standard deviations of 0, 1, 2 and 3 pixels. Sharpness and reference-feature proximity were checked across levels; paired comparisons held tissue composition fixed. This tested controlled blur, not a natural or cross-site shift. | **Controlled Gaussian blur.** CONCH accuracy was 0.789, 0.730, 0.460 and 0.246 across the four levels. The endpoint decline was 0.543 (paired tile-bootstrap 95% CI 0.512–0.574), versus 0.323 for the fixed baseline, whose accuracy fell from 0.932 to 0.609. The excess decline was 0.219 (95% CI 0.182–0.257). *Supported finding: greater blur sensitivity than this comparator under the tested conditions.* Freezing preceded confirmation, but the result is limited to the same source family. Cross-site generalization and patient-level separation from baseline training were not established. Intervals are tile-level, not patient-cluster intervals. Blur was a known axis, not a novel discovery. |
| Kimi r1 | Full CRC-VAL-HE-7K, containing 7,180 images, scored with the official zero-shot protocol and prompt ensemble at the native 224-pixel input size [71, 127, 128]. A class-stratified reference pool of 25,002 NCT images was used only to train the baseline. It had no pixel-hash overlap with the test set and was patient-separated by design. **Baseline:** a fixed ImageNet ResNet-50 with a $k$-nearest-neighbor classifier, kept fixed to match CONCH's zero-shot use, plus a retrained version as an upper bound. | No candidate was abandoned. One condition was chosen from the proposed mechanism: CONCH pretraining was dominated by images near 0.5 $\mu$m pixel$^{-1}$. The condition was specified in writing before CONCH was scored and was evaluated on the first and only examination of the data. No additional analysis was needed to rescue the result. | **Effective acquisition magnification.** Five levels ranged from 0.5 to 8.0 $\mu$m pixel$^{-1}$. CONCH and the baseline received identical pixels at every level, and ordering was verified with a measure independent of CONCH. CONCH accuracy was 0.803, 0.806, 0.778, 0.276, and 0.118, compared with chance at 0.111. The decline was 52.7 percentage points (CI 51.3–54.1), a relative loss of 65.6%; McNemar counts were 4,168 versus 384. Performance was stable from 20× to 10×, declined slightly at 5×, and fell sharply at 2.5× or below. *The decline was substantially larger for CONCH. The baseline received the same degraded images and lost 29.1%. Even after retraining, it retained 0.62 accuracy where CONCH achieved 0.28. The transformed images therefore still contained class information; CONCH made less effective use of it.* |

*Supplementary Table 18 (continued)*

| Run | Data used | Conditions tested | Prespecified condition, result, and decision |
|---|---|---|---|
| Kimi r2 | A 900-image CRC-VAL pilot for discovery; 3,600 held-out NCT images, with 400 per class, for confirmation; full 7,180-image CRC-VAL as an independent-center check [71, 127, 128]. The official CRC prompt ensemble was used unchanged for CONCH and the zero-shot baseline. **Baselines:** OpenCLIP with the same prompts and images, and a DINOv2 encoder with a fixed probe to test whether any model could still recover class information after the image change. | Stain shift was negative at pilot scale, from 0.751 to 0.727, and was kept as a control showing that CONCH was not equally fragile to every image change. Site shift was nearly absent at native resolution, 0.771 versus 0.756, so it did not provide an ordered condition to test. | **Effective spatial resolution.** Six levels from 0.5 to 4.0 $\mu$m pixel$^{-1}$ were fixed after the pilot and confirmed on images not used in the pilot. Ordering was checked with two independent measures, and power was defined in advance. CONCH accuracy fell at every adjacent level: 0.771, 0.697, 0.502, 0.343, 0.188, and 0.138, ending near chance. McNemar counts between the extremes were 2,306 versus 25. The same curve appeared at the independent center, from 0.756 to 0.139. *The decline was substantially larger for CONCH. The general vision–language baseline remained near its low starting level (*$-0.033$ *change), while the information-availability baseline retained 0.667 accuracy where CONCH reached 0.138. Its own decline of 0.302 was retained as a real effect shared across models.* |

**Supplementary Table 19: Level 4 audit detail: ESMFold2.** AutoSPECTRA-guided. Each run predicted post-cutoff RCSB structures, removed related sequences with MMseqs2, tested a model-independent baseline, and applied prespecified decision criteria. Kimi r1 evaluated ESMFold v1 rather than ESMFold2.

| Run | Data used | Conditions tested | Prespecified condition, result, and decision |
|---|---|---|---|
| GPT r1 | RCSB entries released from 2021-10-01 through 2026-06-30 at 3.5 Å resolution or better. MMseqs2 clustering produced 2,530 representatives: 2,000 primary targets and 530 controls [97, 133, 135]. Predictions used three loops, 68 diffusion steps, seed 0, and one A100 GPU; accuracy was measured by TM-score [134]. **Baseline:** coordinate transfer from the best matching PDB chain deposited before the cutoff, scored on the same target chains. | Chain length and homology support were included as documented controls; homology also supplied the baseline. Predicted disorder, coiled-coil content, and low complexity were measured as possible confounders. Membrane topology was considered but not measured, and the scope limitation was stated. | **Novelty of the Pfam-A domain repertoire.** Four ordered levels had median similarities of 1.000, 0.750, 0.667, and 0.250. Statistical power was sufficient at the most novel level. TM-scores were 0.857, 0.796, 0.777, and 0.761. The L0 minus L3 difference was 0.096 (CI 0.075–0.116), and the prespecified monotonic isotonic test passed. The fully adjusted coefficient was 0.053 ($p$ = 0.00044). Homology and experimental resolution caused the largest changes when individually removed from the adjustment, but neither removed the result. *The decline was also observed in the baseline. ESMFold2 declined by 0.096, while template transfer declined much more, by 0.473; the difference was* −0.378. |
| GPT r2 | 46,571 post-cutoff RCSB entries reduced to 2,530 chains after excluding FoldBench targets and examples displayed in the paper [97, 98, 133, 135]. Discovery and confirmation were separated strictly by deposition date. The run folded 435 chains with three loops and 14 steps, the checkpoint's default rather than the paper's setting, and limited the claim accordingly. **Baseline:** template coordinate transfer. | Four conditions were fixed in sequence and all failed: internal sequence-repeat burden, low composition entropy, isolated-chain oligomeric context, and dominance of a single residue type. The entropy result remained negative even after the largest allowed follow-up, and two conditions were significantly opposite to the predicted direction. Cysteine, hydrophobic content, and glycine were rejected in discovery; length and low complexity were treated as documented controls. | **Net positive charge among cryo-EM chains solved in isolation.** This condition was identified by examining the 19 catastrophic failures among the 285 predictions already produced for the four negative tests. The shared feature was named before any additional chain was folded. The condition was then fixed again and tested on 150 sequence clusters not used in any prior prediction. TM-scores were 0.8454, 0.7804, and 0.7437. The high-minus-low difference was −0.1017 (CI −0.1768 to −0.0247), exceeding the prespecified −0.09 threshold. Catastrophic-failure rates were 4%, 14%, and 16%. *The decline was also observed in the baseline: template transfer showed a gap of 0.1152, compared with 0.1017 for ESMFold2.* |

*Supplementary Table 19 (continued)*

| Run | Data used | Conditions tested | Prespecified condition, result, and decision |
|---|---|---|---|
| Opus r1 | 15,648 RCSB candidates reduced by MMseqs2 to 3,137 nonredundant chains. Clusters were separated into 700 discovery, 1,400 confirmation, and 1,037 reserve chains [97, 133, 135]. The run folded 648 discovery chains, 518 confirmation chains, and another 293 chains enriched for orphans. **Baselines:** AlphaFold2 structures from the database, scored identically, and template transfer used only as a measure of intrinsic difficulty. | Length was tested first and was null through about 600 residues. A positive relationship between length and TM-score was explained by TM-score's own length normalization. Predicted disorder was fixed as the first main condition and met the discovery criterion at −0.139 but shrank to −0.032 in the independent confirmation set. The run treated this as an exaggerated discovery effect and did not promote the discovery estimate. | **Coverage-weighted sequence identity to the pre-cutoff PDB**, fixed again after the disorder result failed. Mean TM-score fell from 0.897 for high-homology targets to 0.756 for orphan targets, a decline of 0.141. Spearman correlation was +0.224 in discovery and +0.243 in confirmation. After controlling for length and experimental resolution, it was +0.250 ($p = 3.5 \times 10^{-7}$). *The decline was larger for ESMFold2 than for AlphaFold2. The run specified that template transfer could not fairly judge a homology condition because it necessarily fails on orphan sequences. AlphaFold2 instead declined by 0.063 where ESMFold2 declined by 0.141, a difference of* +0.078 *(CI 0.006–0.150).* |
| Opus r2 | 10,045 RCSB candidates reduced to 2,795, then fixed as 1,250 primary chains and 100 chains reserved for an extreme-length group [97, 133, 135]. The test specification was hashed and written before any prediction was compared with experimental coordinates. **Baseline:** coordinate transfer from the best pre-cutoff chain with coverage at least 0.5, available for 1,192 of 1,350 targets. | Homology to the pre-cutoff PDB was designated as a possible alternative explanation but was null in this dataset (raw $\rho \approx -0.06$; adjusted +0.001). Experimental resolution, composition entropy, and oligomeric copy number were nonsignificant. Very long chains did have a failure tail, but it was shared: the baseline failed for 39% and ESMFold2 for 21%. | **Input sequence length.** Six ordered levels were evaluated against four criteria defined in advance. TM-scores were 0.836, 0.922, 0.922, 0.933, 0.915, and 0.875, peaking at L4 and reaching their minimum at L1. Rank correlation was +0.176, opposite the predicted direction. The difference between the pooled L1–L3 group and L6 was 0.019, below the required 0.05. The adjusted coefficient was +0.019 ($p = 0.11$), and ESMFold2 outperformed the baseline at every level. *None of the four prespecified criteria was met. The planned 768–1,023-residue group could not be evaluated because the shared 40 GB GPU ran out of memory above about 760 residues. That range remains unanswered rather than negative.* |

*Supplementary Table 19 (continued)*

| Run | Data used | Conditions tested | Prespecified condition, result, and decision |
|---|---|---|---|
| Kimi r1 | *ESMFold v1, the predecessor.* 13,225 PDB entries filtered to 10,254 candidates and clustered into 2,145 representatives, with 80 chains per level matched by length group [101, 133, 135]. TM-score was checked against an independent implementation to within $10^{-4}$. **Baseline:** template coordinate transfer. | Low-complexity fraction was fixed as a second condition and failed, running opposite the prediction with only 16 chains in the most extreme group. The original four-level sequence-identity condition also failed its prespecified requirements: it was not monotonic, its extreme difference was 0.081 rather than the required 0.15, and the trend came entirely from the lowest group. | **Orphan status, defined as less than 30% identity to the training set.** This condition was fixed again in a written record and powered in advance at 170 chains per group. It was then confirmed using 340 sequence clusters separate from discovery. Mean TM-score was 0.789 for orphan and 0.875 for non-orphan targets, a gap of 0.086 (effect size 0.43, $p = 4.7 \times 10^{-5}$), with the same direction in all three length groups. The 10th percentile was 0.42 for orphans and 0.69 for non-orphans. *The decline was smaller for ESMFold than for the baseline. Template transfer scored 0.593 on orphans and could not produce a prediction for 143 of 170. The broader graded identity test failed, but redefining it as a prespecified orphan threshold produced a confirmed result.* |
| Kimi r2 | 182,461 pre-cutoff PDB entries yielding 599,471 reference sequences; 32,788 qualifying test entities subsampled and clustered into 1,312 representatives [97, 133, 135]. The test period began after the validation period used in the paper. A pilot with 8 targets per level set the final sample size at 40 per level before confirmation, providing more than 95% power. **Baseline:** coordinate transfer from the best pre-cutoff match. | No condition was abandoned: one condition was fixed and tested at the declared power. Missing electron density and dependence on a protein complex were defined in advance as sensitivity analyses. Removing those cases strengthened the result, so neither explained it. | **Maximum sequence identity to the pre-cutoff PDB.** Four levels followed the paper's homology filter, with observed identities of 0.961, 0.615, 0.319, and 0.000 and balanced lengths. TM-scores were 0.914, 0.894, 0.936, and 0.828. The I1-minus-I4 difference was 0.086 ($p = 3.0 \times 10^{-2}$). Performance was stable from I1 through I3 and then fell at I4. *The decline was also observed in the baseline. At high homology, ESMFold2 and template transfer were similar. At I3, ESMFold2 led by 0.170, and at I4 the template baseline was undefined. The pilot gap of 0.19 shrank to 0.086 in the adequately powered confirmation analysis, and the run reported this discovery overestimate explicitly.* |

**Supplementary Table 20:** **Level 4 audit detail: STATE.** AutoSPECTRA-guided. Released benchmarks hold out cell context, not perturbation identity. Conditions based on the cell context generally failed, while conditions based on the perturbation more often met the prespecified criteria.

| Run | Data used | Conditions tested | Prespecified condition, result, and decision |
|---|---|---|---|
| GPT r1 | Tahoe-100M with both released checkpoints, five held-out contexts, 379 compounds, and 1,136 drug–dose conditions [3, 7]. Compounds were hashed by normalized parent structure so that every dose and context for a compound remained together. Confirmation contained 268 compounds and 4,020 rows, with power defined in advance by Monte Carlo simulation. **Baseline:** an identity predictor that always returned zero fold change, paired within the same bootstrap samples. | S-phase variation in the held-out population produced the largest regression coefficient in the run and exceeded the baseline by +0.119, but did not pass. With only five held-out contexts, the ordering test was coarse, and the five context averages had rank correlation 0.30 with the prespecified order, below the required 0.70. Scarcity of a cell lineage failed twice: its direction reversed after removing the single liver context, and it was defined only after the checkpoint had been examined. | **Baseline transcriptomic distance from the training contexts.** The prespecified ordering was verified against an independent expression atlas at rank correlation 1.0. *STATE improved as distance increased, contrary to the hypothesized decline.* The raw slope was −0.032 and the adjusted slope was −0.150, while the identity baseline became worse over the same rows at +0.163. Every dose group and every leave-one-context-out fit was negative. The contexts were genuinely different, but that difference did not cause STATE to fail. *The run ended without a confirmed failure condition.* |
| GPT r2 | Two evaluation families under one protocol: Replogle–Nadig CRISPR interference across four contexts for the primary condition, and Tahoe-100M across five held-out contexts for the successful condition [3–5, 7]. Code, evaluation and data-loading revisions, four checkpoints, two baseline expression atlases, and official split files were fixed by hash. Each condition was fixed under its own written protocol before its target representation was examined. | Baseline distance across cell lines failed and reversed direction (four-context permutation $p = 0.583$). Weak perturbation magnitude could not be tested: none of 1,385 candidates met the prespecified independence requirement, and the run recorded an empty result rather than relaxing the rule. An external Tahoe analysis was blocked by an unpublished column-to-gene mapping and by an unfinished scan of a 4.09-billion-row table. Three reserve conditions were not scored, and perturbation modality could not be identified from the data. | **Low administered dose.** The condition was fixed before the representation was examined and included 199 drugs with measurements at all three doses in five contexts. Performance was 0.4738 at 5 $\mu$M, 0.4399 at 0.5, and 0.4417 at 0.05. The fixed slope was −0.01692 per standard deviation, the adjusted slope was −0.01592, and the low-minus-high difference was −0.03204. *A baseline that did not use any perturbed outcome was nearly flat at* +0.00173*, so the established decline appeared only in STATE. However, the difference between STATE and the baseline remained within the prespecified comparability margin.* |

*Supplementary Table 20 (continued)*

| Run | Data used | Conditions tested | Prespecified condition, result, and decision |
|---|---|---|---|
| Opus r1 | Released precomputed evaluation package containing 39,808 scored predictions across all three model families [3–7]. Discovery used one Tahoe cell line. Confirmation used a second representation, the few-shot protocol, and each of five held-out lines separately, with the condition fixed first. **Baseline:** a placebo using random gene sets matched to the size of each drug's target list, plus a label permutation. | An initial variance decomposition changed the direction of the search: perturbation identity explained 78–86% of Replogle variance, while held-out context explained less than 1%. True effect strength was the strongest predictor in the run, with $\rho$ as high as 0.93, but could not be used as a condition because it was calculated from the outcome. Chemical novelty was null because all 379 drugs appeared in training. Novelty of the cell context was also null at +0.012. | **Coverage of a drug's targets in the training knockout panel**, calculated from annotations alone. Discovery performance was 0.236, 0.308, and 0.291 across levels, with Spearman +0.263 ($p = 1.5 \times 10^{-5}$). The absolute difference between no targets covered and any target covered was +0.063, or +27% relative. After joint adjustment for all three confounders, the effect remained +0.118 (CI 0.003–0.235). It replicated in both other representations and all five held-out cell lines at +0.19 to +0.29. *The association was specific to STATE relative to the baseline. The size-matched random-gene placebo did not change, separating an effect of training coverage from a simple effect of target count.* |
| Opus r2 | The same package restricted to Replogle–Nadig CRISPR interference: 2,023 genes, four contexts, two protocols, and two representations [3–5]. The trend was fit in two discovery contexts and tested in two held-out contexts, with additional gene-holdout and cross-family confirmation. **Baseline:** none could be calculated because the package contained only final metrics, not raw expression or predictions from other models. The run recorded this as a limitation rather than substituting an unsuitable baseline. | Drug chemical novelty was null; mechanism coverage had the opposite direction; GO annotation count was null, ruling out the explanation that well-annotated genes were simply easier. Cross-context experimental coverage produced only a weak binary effect. Applying the same definition to Tahoe drug targets and Parse ligands produced no effect; this was reported as a limit on the claim's scope. | **Coverage of the perturbed gene's functional neighborhood in training**, measured as GO-BP Jaccard density among the 1,911 annotated panel genes and split into quartiles. Performance was 0.090, 0.108, 0.164, and 0.185 in one representation and increased from 0.040 to 0.111 in the other. Spearman correlations were +0.295 and +0.285 at $n$ = 5,440. After controlling for effect size, the partial correlation was +0.178. The condition was unrelated to cross-context coverage and replicated in all four contexts and both gene-holdout halves. *The association was supported only for direct genetic knockdowns. Four of five prespecified conditions were met; replication in a different data family failed. The claim is therefore limited to this perturbation type. The absence of a computable baseline remains a gap.* |

*Supplementary Table 20 (continued)*

| Run | Data used | Conditions tested | Prespecified condition, result, and decision |
|---|---|---|---|
| Kimi r1 | Four separately trained few-shot checkpoints for Replogle–Nadig, scored for each gene–context pair. The metric was validated against the official per-perturbation table supplied with the weights [3–5]. Discovery used one context. The condition was fixed in an addendum before confirmation in the other three. **Baselines:** two predictors calculated only from training cells: the context's mean training perturbation and the gene's mean across the other three cell lines. | A four-level exposure condition was fixed first and failed. Its curve was not monotonic, and controlling for effect magnitude removed the apparent trend because genes tested in more cell lines tended to be essential genes with larger responses. Continuous embedding similarity was reassigned as partly explained by effect magnitude. Cells per observation were excluded as a nuisance variable. Two other prespecified candidates were left unscored after the stopping rule was reached. | **Novelty of the gene–context pair**, defined only from membership in the fixed training split. Differences between seen and unseen pairs were +0.110, +0.118, +0.118, and +0.117 in the four contexts, with pooled Stouffer $Z = 11.46$. After adjustment for effect magnitude and cell count, coefficients ranged from +0.109 to +0.137. Both baselines changed by at most 0.019 under the same comparison. *The association was specific to STATE relative to the baselines. On unseen pairs, STATE still outperformed both simple baselines in most contexts but fell below one of them in two contexts. The claim concerns the cost of few-shot generalization, not a complete absence of generalization.* |
| Kimi r2 | New inference for every held-out cell in four contexts: 643,413 cells across 2,024 perturbations, seed 42 on one A100 GPU [3–5]. The pipeline reproduced the official per-perturbation table at $r = 0.955$. The confirmation protocol was fixed before scoring and required at least two of three criteria. **Baselines:** cross-context mean transfer and the global training mean, both calculated only from training cell lines. | The initial four-level training-support condition produced mixed results: the bootstrap comparison passed, while the gradient criterion failed at $\rho = -0.013$. This suggested a step change rather than a gradual trend, and the run reported both results. Amount of support among trained perturbations failed: one exposure was enough to fill the perturbation code. Overall performance below the mean baseline was separated as previously published background. | **Novelty of the perturbation across contexts.** This condition was fixed again after the discovery analysis and was explicitly labeled as informed by the observed outcome. It was confirmed in three new contexts with separate checkpoints and different cells. Performance declines were 0.152, 0.114, 0.055, and 0.045 across the four contexts. With no prior exposure, STATE fell below the global-mean baseline in all four. *Part of the decline was shared with the transfer baseline, but the fall below the global-mean baseline was specific to STATE. The mechanism also matched the step-shaped result: untrained one-hot columns had norm 0.233, compared with 5.3–6.1 for trained columns, so the model received an almost empty perturbation code. The conclusion is limited to models that use one-hot perturbation encoding.* |

### References

1. Ektefaie, Y., Shen, A., Bykova, D., Marin, M. G., Zitnik, M. & Farhat, M. Evaluating generalizability of artificial intelligence models for molecular datasets. *Nature Machine Intelligence* **6,** 1512–1524. doi:10.1038/s42256-024-00931-6 (2024).
2. Starace, G. *et al. PaperBench: Evaluating AI's Ability to Replicate AI Research* 2025. arXiv: 2504.01848.
3. Adduri, A. K. *et al.* Predicting cellular responses to perturbation across diverse contexts with State. *bioRxiv.* doi:10.1101/2025.06.26.661135 (2025).
4. Replogle, J. M. *et al.* Mapping information-rich genotype-phenotype landscapes with genome-scale Perturb-seq. *Cell* **185,** 2559–2575.e28. doi:10.1016/j.cell.2022.05.013 (2022).
5. Nadig, A. *et al.* Transcriptome-wide analysis of differential expression in perturbation atlases. *Nature Genetics* **57,** 1228–1237. doi:10.1038/s41588-025-02169-3 (2025).
6. Parse Biosciences. *10 Million Human PBMCs in a Single Experiment* Dataset, Parse Biosciences GigaLab. 2024.
7. Zhang, J. *et al.* Tahoe-100M: A Giga-Scale Single-Cell Perturbation Atlas for Context-Dependent Gene Function and Cellular Modeling. *bioRxiv.* doi:10.1101/2025.02.20.639398 (2025).
8. Srivatsan, S. R. *et al.* Massively multiplex chemical transcriptomics at single-cell resolution. *Science* **367,** 45–51. doi:10.1126/science.aax6234 (2020).
9. McFarland, J. M. *et al.* Multiplexed single-cell transcriptional response profiling to define cancer vulnerabilities and therapeutic mechanism of action. *Nature Communications* **11,** 4296. doi:10.1038/s41467-020-17440-w (2020).
10. Wei, Z. *et al.* Fleming: An AI Agent for Antibiotic Design for Mycobacterium tuberculosis. *bioRxiv.* doi:10.1101/2025.04.01.646719 (2026).
11. Kwa, T. *et al. Measuring AI Ability to Complete Long Software Tasks* in *Advances in Neural Information Processing Systems* (eds Belgrave, D. *et al.*) **38** (Curran Associates, Inc., 2025), 92213–92266. doi:10.52202/085713-3086.
12. Heijl, S., Vroling, B., van den Bergh, T. & Joosten, H.-J. Mind the gap: preventing circularity in missense variant prediction. *bioRxiv.* doi:10.1101/2020.05.06.080424 (2020).
13. Howard, F. M. *et al.* The impact of site-specific digital histology signatures on deep learning model accuracy and bias. *Nature Communications* **12,** 4423. doi:10.1038/s41467-021-24698-1 (2021).
14. Sasse, A. *et al.* Benchmarking of deep neural networks for predicting personal gene expression from DNA sequence highlights shortcomings. *Nature Genetics* **55,** 2060–2064. doi:10.1038/s41588-023-01524-6 (2023).
15. Molotkov, I., Koboldt, D. C. & Artomov, M. SNPred outperforms other ensemble-based SNV pathogenicity predictors and elucidates the challenges of using ClinVar for evaluation of variant classification quality. *medRxiv.* doi:10.1101/2023.09.07.23295192 (2023).

16. Liu, T., Li, K., Wang, Y., Li, H. & Zhao, H. Evaluating the Utilities of Foundation Models in Single-Cell Data Analysis. *Advanced Science* **13,** e14490. doi:10.1002/advs.202514490 (2026).

17. Huang, C. *et al.* Personal transcriptome variation is poorly explained by current genomic deep learning models. *Nature Genetics* **55,** 2056–2059. doi:10.1038/s41588-023-01574-w (2023).

18. Edmunds, N. S., Genc, A. G. & McGuffin, L. J. Benchmarking of AlphaFold2 accuracy self-estimates as indicators of empirical model quality and ranking: a comparison with independent model quality assessment programmes. *Bioinformatics* **40,** btae491. doi:10.1093/bioinformatics/btae491 (2024).

19. Hermann, L., Fiedler, T., Nguyen, H. A., Nowicka, M. & Bartoszewicz, J. M. *Beware of Data Leakage from Protein LLM Pretraining* in *Proceedings of the 19th Machine Learning in Computational Biology meeting* (eds Knowles, D. A. & Mostafavi, S.) **261** (PMLR, 2024), 106–116.

20. Kathail, P., Shuai, R. W., Chung, R., Ye, C. J., Loeb, G. B. & Ioannidis, N. M. Current genomic deep learning models display decreased performance in cell type-specific accessible regions. *Genome Biology* **25,** 202. doi:10.1186/s13059-024-03335-2 (2024).

21. Chakravarty, D. *et al.* AlphaFold predictions of fold-switched conformations are driven by structure memorization. *Nature Communications* **15,** 7296. doi:10.1038/s41467-024-51801-z (2024).

22. Rastogi, R., Reddy, A. J., Chung, R. & Ioannidis, N. M. Fine-tuning sequence-to-expression models on personal genome and transcriptome data. *Genome Biology* **27,** 234. doi:10.1186/s13059-026-04091-1 (2026).

23. Vedula, S., Bronstein, A. M. & Marx, A. Improving prediction accuracy in chimeric proteins with windowed multiple sequence alignment. *Computational and Structural Biotechnology Journal* **27,** 3292–3298. doi:10.1016/j.csbj.2025.07.039 (2025).

24. Hernández Trapero, R., Badonyi, M., Gerasimavicius, L. & Marsh, J. A. Complementary Roles of Structure and Variant Effect Predictors in RyR1 Clinical Interpretation. *Human Mutation* **2025,** 1834898. doi:10.1155/humu/1834898 (2025).

25. Rafi, A. M., Kiyota, B., Yachie, N. & de Boer, C. Detecting and avoiding homology-based data leakage in genome-trained sequence models. *bioRxiv.* doi:10.1101/2025.01.22.634321 (2025).

26. Spiro, A. E. *et al.* A scalable approach to investigating sequence-to-function predictions from personal genomes. *Nature Methods* **23,** 1308–1312. doi:10.1038/s41592-026-03124-8 (2026).

27. Joeres, R., Blumenthal, D. B. & Kalinina, O. V. Data splitting to avoid information leakage with DataSAIL. *Nature Communications* **16,** 3337. doi:10.1038/s41467-025-58606-8 (2025).

28. Kedzierska, K. Z., Crawford, L., Amini, A. P. & Lu, A. X. Zero-shot evaluation reveals limitations of single-cell foundation models. *Genome Biology* **26,** 101. doi:10.1186/s13059-025-03574-x (2025).

29. Livesey, B. J. & Marsh, J. A. Variant effect predictor correlation with functional assays is reflective of clinical classification performance. *Genome Biology* **26,** 104. doi:10.1186/s13059-025-03575-w (2025).

30. Csendes, G., Sanz, G., Szalay, K. Z. & Szalai, B. Benchmarking foundation cell models for post-perturbation RNA-seq prediction. *BMC Genomics* **26,** 393. doi:10.1186/s12864-025-11600-2 (2025).

31. Atti, S. & Subramaniam, S. Fundamental Limitations of Foundation Models in Single-Cell Transcriptomics. *bioRxiv.* doi:10.1101/2025.06.26.661767 (2025).

32. Ahlmann-Eltze, C., Huber, W. & Anders, S. Deep-learning-based gene perturbation effect prediction does not yet outperform simple linear baselines. *Nature Methods* **22,** 1657–1661. doi:10.1038/s41592-025-02772-6 (2025).

33. Ribeiro-dos-Santos, A. M. & Maurano, M. T. Iterative improvement of deep learning models using synthetic regulatory genomics. *bioRxiv.* doi:10.1101/2025.02.04.636130 (2025).

34. Perkins-Jechow, B. H. *et al.* Challenging AlphaFold in predicting proteins with large-scale allosteric transitions. *Communications Chemistry* **8,** 378. doi:10.1038/s42004-025-01763-0 (2025).

35. Drusinsky, S., Whalen, S. & Pollard, K. S. Deep-learning prediction of gene expression from personal genomes. *Genome Biology* **27,** 19. doi:10.1186/s13059-025-03926-7 (2026).

36. Thiringer, E., Gustafsson, F. K., Eriksson, K. L. & Rantalainen, M. *Scanner-Induced Domain Shifts Undermine the Robustness of Pathology Foundation Models* 2026. doi:10.48550/arXiv.2601.04163. arXiv: 2601.04163v1.

37. Souza, H. & Mehta, P. *Parameter-free representations outperform single-cell foundation models on downstream benchmarks* 2026. arXiv: 2602.16696v1.

38. Feldman, J., Brogi, M. & Skolnick, J. Adversarial Sequence Mutations in AlphaFold and ESMFold Reveal Nonphysical Structural Invariance, Confidence Failures, and Concerns for Protein Design. *Computational and Structural Biotechnology Journal* **35,** 0142. doi:10.34133/csbj.0142 (2026).

39. Mahtha, S. K., Venkadesan, S. & Mohanty, D. Comparative evaluation of the prediction accuracy of AlphaFold and ESMFold for monomeric and dimeric proteins. *NAR Genomics and Bioinformatics* **8,** lqag002. doi:10.1093/nargab/lqag002 (2026).

40. Lu, T., Lu, A. X. & Moses, A. M. *Random Embeddings and Linear Regression can Predict Protein Function* 2021. doi:10.48550/arXiv.2104.14661. arXiv: 2104.14661v1.

41. Tabet, D. R. *et al.* Benchmarking computational variant effect predictors by their ability to infer human traits. *Genome Biology* **25,** 172. doi:10.1186/s13059-024-03314-7 (2024).

42. Kömen, J., Marienwald, H., Dippel, J. & Hense, J. *Do Histopathological Foundation Models Eliminate Batch Effects? A Comparative Study* 2024. doi:10.48550/arXiv.2411.05489. arXiv: 2411.05489v1.

43. Haley, O. C. *et al.* Why do some predicted protein structures fold poorly? Benchmarking AlphaFold, ESMFold, and Boltz in maize. *bioRxiv.* doi:10.1101/2025.07.05.663230 (2025).

44. Lu, B., Liu, X., Lin, P.-Y. & Brandes, N. Genomic heterogeneity inflates the performance of variant pathogenicity predictions. *bioRxiv.* doi:10.1101/2025.09.05.674459 (2025).

45. Hýsková, A., Maršálková, E. & Šimeček, P. Balancing speed and precision in protein folding: a comparison of AlphaFold2, ESMFold, and OmegaFold. *Frontiers in Genetics* **16,** 1715037. doi:10.3389/fgene.2025.1715037 (2026).

46. Garcia, M., Dixit, S. M. & Rocklin, G. J. Evaluating zero-shot prediction of monomeric protein design success by AlphaFold, ESMFold, and ProteinMPNN. *Protein Science* **35,** e70453. doi:10.1002/pro.70453 (2026).

47. Hoffing, R. *et al.* Ancestry-specific performance of variant effect predictors in clinical variant classification. *bioRxiv.* doi:10.64898/2026.02.14.705914 (2026).

48. Henriksen, A. L. *et al. Enabling clinical use of foundation models for computational pathology* 2026. doi:10.48550/arXiv.2602.22347. arXiv: 2602.22347v2.

49. Mathur, V. & Sachidanandam, R. Benchmarking DNA Foundation Models: Biological Blind Spots in Evo2 Variant-Effect Prediction. *bioRxiv.* doi:10.64898/2026.03.10.710786 (2026).

50. Cheng, J. *et al.* Accurate proteome-wide missense variant effect prediction with AlphaMissense. *Science* **381,** eadg7492. doi:10.1126/science.adg7492 (2023).

51. Rastogi, R. *et al.* Critical assessment of missense variant effect predictors on disease-relevant variant data. *Human Genetics* **144,** 281–293. doi:10.1007/s00439-025-02732-2 (2025).

52. Kong, S. W. *et al.* Discordance between a deep learning model and clinical-grade variant pathogenicity classification in a rare disease cohort. *npj Genomic Medicine* **10,** 17. doi:10.1038/s41525-025-00480-w (2025).

53. Pejaver, V. *et al.* Calibration of computational tools for missense variant pathogenicity classification and ClinGen recommendations for PP3/BP4 criteria. *The American Journal of Human Genetics* **109,** 2163–2177. doi:10.1016/j.ajhg.2022.10.013 (2022).

54. Curtis, D. Assessment of ability of AlphaMissense to identify variants affecting susceptibility to common disease. *European Journal of Human Genetics* **32,** 1419–1427. doi:10.1038/s41431-024-01675-y (2024).

55. Luppino, F., Lenz, S., Chow, C. F. W. & Toth-Petroczy, A. Deep learning tools predict variants in disordered regions with lower sensitivity. *BMC Genomics* **26,** 367. doi:10.1186/s12864-025-11534-9 (2025).

56. Wang, Y.-H., Masson, E., Liao, Z., Férec, C., Zou, W.-B. & Chen, J.-M. Strengths and limitations of AlphaMissense in CPA1 missense variant classification. *Gut* **73,** e42. doi:10.1136/gutjnl-2024-332120 (2024).

57. Ljungdahl, A. *et al.* AlphaMissense is better correlated with functional assays of missense impact than earlier prediction algorithms. *bioRxiv.* doi:10.1101/2023.10.24.562294 (2023).

58. Murali, H., Wang, P., Liao, E. C. & Wang, K. Genetic variant classification by predicted protein structure: A case study on IRF6. *Computational and Structural Biotechnology Journal* **23,** 892–904. doi:10.1016/j.csbj.2024.01.019 (2024).

59. McDonald, E. F., Oliver, K. E., Schlebach, J. P., Meiler, J. & Plate, L. Benchmarking AlphaMissense pathogenicity predictions against cystic fibrosis variants. *PLOS ONE* **19,** e0297560. doi:10.1371/journal.pone.0297560 (2024).

60. Chong, R. *et al.* A Multiplexed Assay for Exon Recognition Reveals that an Unappreciated Fraction of Rare Genetic Variants Cause Large-Effect Splicing Disruptions. *Molecular Cell* **73,** 183–194.e8. doi:10.1016/j.molcel.2018.10.037 (2019).

61. Richards, S. *et al.* Standards and guidelines for the interpretation of sequence variants: a joint consensus recommendation of the American College of Medical Genetics and Genomics and the Association for Molecular Pathology. *Genetics in Medicine* **17,** 405–423. doi:10.1038/gim.2015.30 (2015).

62. Tordai, H., Torres, O., Csépi, M., Padányi, R., Lukács, G. L. & Hegedűs, T. Analysis of AlphaMissense data in different protein groups and structural context. *Scientific Data* **11,** 495. doi:10.1038/s41597-024-03327-8 (2024).

63. Fawzy, M. & Marsh, J. A. Assessing variant effect predictors and disease mechanisms in intrinsically disordered proteins. *bioRxiv.* doi:10.1101/2025.04.01.646619 (2025).

64. Pillai, J., Sridhar, A., Sung, K., Shi, L. & Wu, C. Deciphering gain-of-function from loss-of-function variants with AlphaMissense: A case study with the mechanosensitive PIEZO1 ion channel protein. *bioRxiv.* doi:10.1101/2025.07.03.662957 (2025).

65. Yazaki, S., Pei, X., Powell, S., Khan, A., Setton, J. & Riaz, N. AlphaMissense for Identifying Pathogenic Missense Mutations in DNA Damage Repair Genes in Cancer. *JCO Precision Oncology* **9,** e2400908. doi:10.1200/PO-24-00908 (2025).

66. Yang, Y., Sun, W., Zhang, J., Liu, Y. & Li, M. An interpretable molecular framework for predicting cancer driver missense mutations. *bioRxiv.* doi:10.1101/2025.09.03.674130 (2025).

67. Radjasandirane, R., Diharce, J., Gelly, J.-C. & de Brevern, A. G. Insights for variant clinical interpretation based on a benchmark of 65 variant effect predictors. *Genomics* **117,** 111036. doi:10.1016/j.ygeno.2025.111036 (2025).

68. Jumper, J. *et al.* Highly accurate protein structure prediction with AlphaFold. *Nature* **596,** 583–589. doi:10.1038/s41586-021-03819-2 (2021).

69. Karczewski, K. J. *et al.* The mutational constraint spectrum quantified from variation in 141,456 humans. *Nature* **581,** 434–443. doi:10.1038/s41586-020-2308-6 (2020).

70. Notin, P. *et al.* ProteinGym: Large-scale benchmarks for protein fitness prediction and design. *Advances in Neural Information Processing Systems* **36** (2023).

71. Lu, M. Y. *et al.* A visual-language foundation model for computational pathology. *Nature Medicine* **30,** 863–874 (2024).

72. Neidlinger, P. *et al.* Benchmarking foundation models as feature extractors for weakly supervised computational pathology. *Nature Biomedical Engineering* **10,** 1113–1123. doi:10.1038/s41551-025-01516-3 (2026).

73. Kömen, J. *et al.* Towards robust foundation models for digital pathology. *Nature Communications* **17,** 5218. doi:10.1038/s41467-026-73923-2 (2026).

74. Vaidya, A. *et al.* Demographic bias in misdiagnosis by computational pathology models. *Nature Medicine* **30,** 1174–1190. doi:10.1038/s41591-024-02885-z (2024).

75. Gustafsson, F. K. & Rantalainen, M. *Evaluating Computational Pathology Foundation Models for Prostate Cancer Grading under Distribution Shifts* 2024. doi:10.48550/arXiv.2410.06723. arXiv: 2410.06723.

76. Lee, J., Lim, J., Byeon, K. & Kwak, J. T. *Benchmarking Pathology Foundation Models: Adaptation Strategies and Scenarios* 2024. doi:10.48550/arXiv.2410.16038. arXiv: 2410.16038.

77. Sharma, V., Alagha, A., Khellaf, A., Trinh, V. Q.-H. & Hosseini, M. S. *Investigating Zero-Shot Diagnostic Pathology in Vision-Language Models with Efficient Prompt Design* 2025. doi:10.48550/arXiv.2505.00134. arXiv: 2505.00134.

78. Alhamoud, K. *et al. Vision-Language Models Do Not Understand Negation* in *Proceedings of the IEEE/CVF Conference on Computer Vision and Pattern Recognition (CVPR)* (2025), 29612–29622.

79. De Jong, E. D., Marcus, E. & Teuwen, J. *Current Pathology Foundation Models are unrobust to Medical Center Differences* 2025. doi:10.48550/arXiv.2501.18055. arXiv: 2501.18055v2.

80. Mishra, V. & Lotter, W. *Comparing Computational Pathology Foundation Models using Representational Similarity Analysis* 2025. doi:10.48550/arXiv.2509.15482. arXiv: 2509.15482.

81. Carloni, G. *et al. Pathology Foundation Models are Scanner Sensitive: Benchmark and Mitigation with Contrastive ScanGen Loss* in *Foundation Models for General Medical AI (MedAGI), MICCAI 2025 Workshop* **16112** (Springer, 2025), 44–53. doi:10.1007/978-3-032-07845-2_5.

82. Chai, B. *et al.* Impact of tissue staining and scanner variation on the performance of pathology foundation models: a study of sarcomas and their mimics. *The Journal of Pathology: Clinical Research* **12,** e70080. doi:10.1002/2056-4538.70080 (2026).

83. Lu, M. Y. *et al. Visual Language Pretrained Multiple Instance Zero-Shot Transfer for Histopathology Images* in *Proceedings of the IEEE/CVF Conference on Computer Vision and Pattern Recognition (CVPR)* (2023), 19764–19775.

84. Ding, T. *et al.* A multimodal whole-slide foundation model for pathology. *Nature Medicine* **31,** 3749–3761. doi:10.1038/s41591-025-03982-3 (2025).

85. Ma, J. *et al. PathBench: A comprehensive comparison benchmark for pathology foundation models towards precision oncology* 2025. doi:10.48550/arXiv.2505.20202. arXiv: 2505.20202.

86. Ramchandani, L., Tinaikar, A., Das, D. K., Garg, R. & Thomas, T. *Benchmarking Computational Pathology Foundation Models for Semantic Segmentation* 2026. doi:10.48550/arXiv.2602.18747. arXiv: 2602.18747.

87. Bilal, M. *et al.* Benchmarking pathology foundation models for predicting microsatellite instability in colorectal cancer histopathology. *Computerized Medical Imaging and Graphics* **127,** 102680. doi:10.1016/j.compmedimag.2025.102680 (2026).

88. Huang, Y. *et al.* Knowledge-guided adaptation of pathology foundation models effectively improves cross-domain generalization and demographic fairness. *Nature Communications* **16.** doi:10.1038/s41467-025-66300-y (2025).

89. Lin, S.-Y. *et al.* Contrastive learning enhances fairness in pathology artificial intelligence systems. *Cell Reports Medicine* **6,** 102527. doi:10.1016/j.xcrm.2025.102527 (2025).

90. Zhang, X. *et al. PMC-VQA: Visual Instruction Tuning for Medical Visual Question Answering* 2023. doi:10.48550/arXiv.2305.10415. arXiv: 2305.10415.

91. Zhang, S. *et al.* A Multimodal Biomedical Foundation Model Trained from Fifteen Million Image–Text Pairs. *NEJM AI* **2.** doi:10.1056/AIoa2400640 (2025).

92. Baghbanzadeh, N., Ashkezari, S., Dolatabadi, E. & Afkanpour, A. *Open-PMC-18M: A High-Fidelity Large Scale Medical Dataset for Multimodal Representation Learning* 2025. doi:10.48550/arXiv.2506.02738. arXiv: 2506.02738.

93. Gallagher-Syed, A., Pontarini, E., Lewis, M. J., Barnes, M. R. & Slabaugh, G. *Going Beyond H&E and Oncology: How Do Histopathology Foundation Models Perform for Multi-stain IHC and Immunology?* in *Workshop on Advancements in Medical Foundation Models, NeurIPS 2024* (2024). doi:10.48550/arXiv.2410.21560. arXiv: 2410.21560.

94. Campanella, G. *et al. A Clinical Benchmark of Public Self-Supervised Pathology Foundation Models* 2024. doi:10.48550/arXiv.2407.06508. arXiv: 2407.06508.

95. Radford, A. *et al. Learning Transferable Visual Models From Natural Language Supervision* in *Proceedings of the 38th International Conference on Machine Learning (ICML)* **139** (2021), 8748–8763.

96. Yu, J., Wang, Z., Vasudevan, V., Yeung, L., Seyedhosseini, M. & Wu, Y. CoCa: Contrastive Captioners are Image-Text Foundation Models. *Transactions on Machine Learning Research* (2022).

97. Chan Zuckerberg Biohub. *ESMFold2* https://github.com/Biohub/esm. 2026.

98. Xu, S. *et al.* Benchmarking all-atom biomolecular structure prediction with FoldBench. *Nature Communications* **17,** 442. doi:10.1038/s41467-025-67127-3 (2026).

99. Chakravarty, D., Lee, M. & Porter, L. L. Proteins with alternative folds reveal blind spots in AlphaFold-based protein structure prediction. *Current Opinion in Structural Biology* **90,** 102973. doi:10.1016/j.sbi.2024.102973 (2025).

100. Buel, G. R. & Walters, K. J. Can AlphaFold2 predict the impact of missense mutations on structure? *Nature Structural & Molecular Biology* **29,** 1–2. doi:10.1038/s41594-021-00714-2 (2022).

101. Lin, Z. *et al.* Evolutionary-scale prediction of atomic-level protein structure with a language model. *Science* **379,** 1123–1130. doi:10.1126/science.ade2574 (2023).

102. Zhang, Z., Wayment-Steele, H. K., Brixi, G., Wang, H., Kern, D. & Ovchinnikov, S. Protein language models learn evolutionary statistics of interacting sequence motifs. *Proceedings of the National Academy of Sciences* **121,** e2406285121. doi:10.1073/pnas.2406285121 (2024).

103. Gopalan, S. & Narayanan, S. *Hallucinations in AlphaFold3 for Intrinsically Disordered Proteins with disorder in Biological Process Residues* 2025. doi:10.48550/arXiv.2510.15939. arXiv: 2510.15939.

104. Hou, C., Zhao, H. & Shen, Y. Protein language models trained on biophysical dynamics inform mutation effects. *Proceedings of the National Academy of Sciences* **123.** doi:10.1073/pnas.2530466123 (2026).

105. Yin, R. & Pierce, B. G. Evaluation of AlphaFold antibody–antigen modeling with implications for improving predictive accuracy. *Protein Science* **33,** e4865. doi:10.1002/pro.4865 (2024).

106. Kretsch, R. C. *et al.* Assessment of Nucleic Acid Structure Prediction in CASP16. *Proteins: Structure, Function, and Bioinformatics* **94,** 192–217. doi:10.1002/prot.70072 (2025).

107. Wittmann, B. J. *et al.* Strengthening nucleic acid biosecurity screening against generative protein design tools. *Science* **390,** 82–87. doi:10.1126/science.adu8578 (2025).

108. Jing, X., Wu, F., Luo, X. & Xu, J. Single-sequence protein structure prediction by integrating protein language models. *Proceedings of the National Academy of Sciences* **121,** e2308788121. doi:10.1073/pnas.2308788121 (2024).

109. Hegedűs, T., Geisler, M., Lukács, G. L. & Farkas, B. Ins and outs of AlphaFold2 transmembrane protein structure predictions. *Cellular and Molecular Life Sciences* **79,** 73. doi:10.1007/s00018-021-04112-1 (2022).

110. Verkuil, R. *et al.* Language models generalize beyond natural proteins. *bioRxiv.* doi:10.1101/2022.12.21.521521 (2022).

111. Kryshtafovych, A., Schwede, T., Topf, M., Fidelis, K. & Moult, J. Critical assessment of methods of protein structure prediction (CASP) — Round XIV. *Proteins: Structure, Function, and Bioinformatics* **89,** 1607–1617. doi:10.1002/prot.26237 (2021).

112. Qi, S.-a. & Chapfuwa, P. Mechanisms Matter: Transportability of Cellular Perturbation Effects. *bioRxiv.* doi:10.64898/2026.05.08.723625 (2026).

113. Dibaeinia, P. *et al.* Virtual Cells Need Context, Not Just Scale. *bioRxiv.* doi:10.64898/2026.02.04.703804 (2026).

114. Viñas Torné, R. *et al.* Systema: a framework for evaluating genetic perturbation response prediction beyond systematic variation. *Nature Biotechnology* **44,** 1050–1059. doi:10.1038/s41587-025-02777-8 (2026).

115. Wei, Z. *et al.* Benchmarking algorithms for generalizable single-cell perturbation response prediction. *Nature Methods* **23,** 451–464. doi:10.1038/s41592-025-02980-0 (2026).

116. Mao, X. *et al. Benchmarking virtual cell models for in-the-wild perturbation response* 2026. doi:10.48550/arXiv.2604.27646. arXiv: 2604.27646.

117. Marrakchi, Y., D'Ascenzo, D. & Cultrera di Montesano, S. *Score Distributions, Not Cells: Evaluating Single-Cell Perturbations Under Class Overlap* 2026. doi:10.48550/arXiv.2607.04595. arXiv: 2607.04595.

118. Squair, J. W. *et al.* Confronting false discoveries in single-cell differential expression. *Nature Communications* **12,** 5692. doi:10.1038/s41467-021-25960-2 (2021).

119. Roohani, Y. H. *et al.* Virtual Cell Challenge: Toward a Turing test for the virtual cell. *Cell* **188,** 3370–3374. doi:10.1016/j.cell.2025.06.008 (2025).

120. Wenteler, A. *et al. PertEval-scFM: Benchmarking Single-Cell Foundation Models for Perturbation Effect Prediction* in *Proceedings of the 42nd International Conference on Machine Learning* **267** (2025), 66633–66677.

121. Wu, Y. *et al. PerturBench: Benchmarking Machine Learning Models for Cellular Perturbation Analysis* in *Advances in Neural Information Processing Systems* **38** (2025), 106937–106977.

122. Weidener, L., Brkić, M., Jovanović, M., Ulgac, E. & Meduri, A. VCBench: A Multi-Dimensional Benchmark for Single-Cell Foundation Models. *bioRxiv.* doi:10.64898/2026.06.18.733146 (2026).

123. Landrum, M. J. *et al.* ClinVar: improving access to variant interpretations and supporting evidence. *Nucleic Acids Research* **46,** D1062–D1067. doi:10.1093/nar/gkx1153 (2018).

124. Grantham, R. Amino acid difference formula to help explain protein evolution. *Science* **185,** 862–864. doi:10.1126/science.185.4154.862 (1974).

125. Emenecker, R. J., Griffith, D. & Holehouse, A. S. Metapredict: a fast, accurate, and easy-to-use predictor of consensus disorder and structure. *Biophysical Journal* **120,** 4312–4319. doi:10.1016/j.bpj.2021.08.039 (2021).

126. The UniProt Consortium. UniProt: the Universal Protein Knowledgebase in 2025. *Nucleic Acids Research* **53,** D609–D617. doi:10.1093/nar/gkae1010 (2025).

127. Kather, J. N., Halama, N. & Marx, A. *100,000 histological images of human colorectal cancer and healthy tissue* Zenodo. 2018. doi:10.5281/zenodo.1214456.

128. Kather, J. N. *et al.* Predicting survival from colorectal cancer histology slides using deep learning: A retrospective multicenter study. *PLoS Medicine* **16,** e1002730. doi:10.1371/journal.pmed.1002730 (2019).

129. Subramanian, B. *et al. STARC-9: A Large-scale Dataset for Multi-Class Tissue Classification for CRC Histopathology* in *The Thirty-ninth Annual Conference on Neural Information Processing Systems Datasets and Benchmarks Track* (2025).

130. Borkowski, A. A., Bui, M. M., Thomas, L. B., Wilson, C. P., DeLand, L. A. & Mastorides, S. M. *Lung and Colon Cancer Histopathological Image Dataset (LC25000)* 2019. doi:10.48550/arXiv.1912.12142. arXiv: 1912.12142.

131. Veeling, B. S., Linmans, J., Winkens, J., Cohen, T. & Welling, M. *Rotation Equivariant CNNs for Digital Pathology* in *Medical Image Computing and Computer Assisted Intervention (MICCAI)* (2018), 210–218. doi:10.1007/978-3-030-00934-2_24.

132. Huang, Z., Bianchi, F., Yuksekgonul, M., Montine, T. J. & Zou, J. A visual–language foundation model for pathology image analysis using medical Twitter. *Nature Medicine* **29,** 2307–2316. doi:10.1038/s41591-023-02504-3 (2023).

133. Burley, S. K. *et al.* Updated Resources for Exploring Experimentally-Determined PDB Structures and Computed Structure Models at the RCSB Protein Data Bank. *Nucleic Acids Research* **53,** D564–D574. doi:10.1093/nar/gkae1091 (2025).

134. Zhang, Y. & Skolnick, J. Scoring function for automated assessment of protein structure template quality. *Proteins: Structure, Function, and Bioinformatics* **57,** 702–710. doi:10.1002/prot.20264 (2004).

135. Steinegger, M. & Söding, J. MMseqs2 enables sensitive protein sequence searching for the analysis of massive data sets. *Nature Biotechnology* **35,** 1026–1028. doi:10.1038/nbt.3988 (2017).

136. Mariani, V., Biasini, M., Barbato, A. & Schwede, T. lDDT: a local superposition-free score for comparing protein structures and models using distance difference tests. *Bioinformatics* **29,** 2722–2728. doi:10.1093/bioinformatics/btt473 (2013).

137. Peidli, S. *et al.* scPerturb: harmonized single-cell perturbation data. *Nature Methods* **21,** 531–540. doi:10.1038/s41592-023-02144-y (2024).

138. Laine, E., Karami, Y. & Carbone, A. GEMME: A Simple and Fast Global Epistatic Model Predicting Mutational Effects. *Molecular Biology and Evolution* **36,** 2604–2619. doi:10.1093/molbev/msz179 (2019).

139. Frazer, J. *et al.* Disease variant prediction with deep generative models of evolutionary data. *Nature* **599,** 91–95. doi:10.1038/s41586-021-04043-8 (2021).

140. Dauparas, J. *et al.* Robust deep learning–based protein sequence design using ProteinMPNN. *Science* **378,** 49–56. doi:10.1126/science.add2187 (2022).

141. Hsu, C. *et al. Learning inverse folding from millions of predicted structures* in *Proceedings of the 39th International Conference on Machine Learning (ICML)* **162** (2022), 8946–8970.

142. Hopf, T. A. *et al.* Mutation effects predicted from sequence co-variation. *Nature Biotechnology* **35,** 128–135. doi:10.1038/nbt.3769 (2017).

143. Kim, S. *et al.* PubChem 2025 update. *Nucleic Acids Research* **53,** D1516–D1525. doi:10.1093/nar/gkae1059 (2025).

144. Frankish, A. *et al.* GENCODE: reference annotation for the human and mouse genomes in 2023. *Nucleic Acids Research* **51,** D942–D949. doi:10.1093/nar/gkac1071 (2023).

145. Henikoff, S. & Henikoff, J. G. Amino acid substitution matrices from protein blocks. *Proceedings of the National Academy of Sciences* **89,** 10915–10919. doi:10.1073/pnas.89.22.10915 (1992).

146. Smedley, D. *et al.* The BioMart community portal: an innovative alternative to large, centralized data repositories. *Nucleic Acids Research* **43,** W589–W598. doi:10.1093/nar/gkv350 (2015).

147. Pollard, K. S., Hubisz, M. J., Rosenbloom, K. R. & Siepel, A. Detection of nonneutral substitution rates on mammalian phylogenies. *Genome Research* **20,** 110–121. doi:10.1101/gr.097857.109 (2010).

148. Howard, A. G. *et al. MobileNets: Efficient Convolutional Neural Networks for Mobile Vision Applications* 2017. doi:10.48550/arXiv.1704.04861. arXiv: 1704.04861.
